# Spateo: multidimensional spatiotemporal modeling of single-cell spatial transcriptomics

**DOI:** 10.1101/2022.12.07.519417

**Authors:** Xiaojie Qiu, Daniel Y. Zhu, Jiajun Yao, Zehua Jing, Lulu Zuo, Mingyue Wang, Kyung Hoi (Joseph) Min, Hailin Pan, Shuai Wang, Sha Liao, Yiwei Lai, Shijie Hao, Yuancheng Ryan Lu, Matthew Hill, Jorge D. Martin-Rufino, Chen Weng, Anna Maria Riera-Escandell, Mengnan Chen, Liang Wu, Yong Zhang, Xiaoyu Wei, Mei Li, Xin Huang, Rong Xiang, Zhuoxuan Yang, Chao Liu, Tianyi Xia, Yingxin Liang, Junqiang Xu, Qinan Hu, Yuhui Hu, Hongmei Zhu, Yuxiang Li, Ao Chen, Miguel A. Esteban, Ying Gu, Douglas A. Lauffenburger, Xun Xu, Longqi Liu, Jonathan S. Weissman, Shiping Liu, Yinqi Bai

## Abstract

Cells do not live in a vacuum, but in a milieu defined by cell–cell communication that can be measured via emerging high-resolution spatial transcriptomics approaches. However, analytical tools that fully leverage such data for kinetic modeling remain lacking. Here we present Spateo (aristoteleo/spateo-release), a general framework for quantitative spatiotemporal modeling of single-cell resolution spatial transcriptomics. Spateo delivers novel methods for digitizing spatial layers/columns to identify spatially-polar genes, and develops a comprehensive framework of cell-cell interaction to reveal spatial effects of niche factors and cell type-specific ligand-receptor interactions. Furthermore, Spateo reconstructs 3D models of whole embryos, and performs 3D morphometric analyses. Lastly, Spateo introduces the concept of “morphometric vector field” of cell migrations, and integrates spatial differential geometry to unveil regulatory programs underlying various organogenesis patterns of Drosophila. Thus, Spateo enables the study of the ecology of organs at a molecular level in 3D space, beyond isolated single cells.

## Introduction

Cells exist in complex three-dimensional microenvironments shaped by the collective activities of neighboring cells of many types and in many states, that together dictate tissue-level form and function (Halpern et al., 2017; Lécuyer et al., 2007; Lee et al., 2022; Lohoff et al., 2022; McCaffrey et al., 2022; Parigi et al., 2022; Scadden, 2014) and change over time (Colom et al., 2020; Garcia-Alonso et al., 2021; Mantri et al., 2021; Misra et al., 2021). Within these microenvironments, constituent cells influence one another through direct cell-cell interactions, physical contacts (Kechagia et al., 2019) and biochemical cues derived from secreted factors (Batlle et al., 2002; Scadden, 2006). This multidimensionality has been difficult to assay due to methodological limitations that constrain high-quality profiling of multiple tissue slices across space and time, and the lack of suitable computational analysis tools capable of linking molecular measurements to 4D processes at scale.

The advent of spatial transcriptomics (ST) has enabled characterization of intercellular interactions (Arnol et al., 2019; Cang and Nie, 2020; Ji et al., 2020; Tanevski et al., 2022; Yuan and Bar-Joseph, 2020), investigation of processes underlying abnormal, pathological tissue organization in health and disease (Boyd et al., 2020; Chen et al., 2020; Elmentaite et al., 2021; Hwang et al., 2022; Janosevic et al., 2021; Ji et al., 2020; Kuppe et al., 2022; Ma et al., 2021;Maniatis et al., 2019), and general attribution of sources of cell-to-cell heterogeneity and dynamism (Asp et al., 2019; Fawkner-Corbett et al., 2021; Garcia-Alonso et al., 2021). These methods can be categorized as imaging-based *in situ* approaches, such as FISSEQ (Lee et al.,2014), STARMap (Wang et al., 2018), expansion sequencing (ExSeq) (Alon et al., 2021), seqFISH (Eng et al., 2019; Lubeck et al., 2014) and MERFISH (Chen et al., 2015), or *ex situ* sequencing-based approaches such as Visium (Ståhl et al., 2016), Slide-seq (Rodriques et al.,2019; Stickels et al., 2021), DBiT-seq (Liu et al., 2020), and HDST (Vickovic et al., 2019) with relative low spatial resolution, and the emerging single cell/subcellar resolution approaches, including Seq-Scope (Cho et al., 2021), Pixel-seq (Fu et al., 2021b) and the recent Stereo-seq (Chen et al., 2022; Liu et al., 2022; Wang et al., 2022; Wei et al., 2022; Xia et al., 2022), characterized by a large field of view (up to 13 x 13 cm^2^) that crucially enables simultaneous profiling of multiple adjacent tissue slices while eliminating platform effects, potentiating efforts to characterize the multidimensional environments of tissues and organs.

While there has been rapid advances in the technologies for collecting high resolution spatial transcriptomic data, the computational tools for analyzing such data lag far behind, especially when applied to jointly study cellular resolution and system-level spatiotemporal dynamics because no uniform multi-scale kinetic approaches are available. To best use the generated data, especially those from high resolution, large-field approaches, and to perform kinetic and predictive analyses that result in novel biological insights, a considerable advance in computational and toolkit development is necessary. Several ST analysis methods exist for a number of applications, such as performing spatially-aware clustering (Dong and Zhang, 2022;Fu et al., 2021a; Hu et al., 2021; Zhao et al., 2021; Zhu et al., 2018), finding potential hotspots of ligand-receptor interactions (Arnol et al., 2019; Cang and Nie, 2020; Tanevski et al., 2022; Yuan and Bar-Joseph, 2020), identifying gene expression modules over space (Jerby-Arnon and Regev, 2022) and identifying spatially-variable genes (Edsgärd et al., 2018; Sun et al.,2020; Svensson et al., 2018). Additionally, integrative toolkits (e.g. Giotto (Dries et al., 2021), stLearn (Pham et al., 2020) and Squidpy (Palla et al., 2022)) provide statistical libraries and facile visualization options in an uniformed environment. Although powerful, existing spatial analyses methods and tools are in general not suitable for high-resolution spatial transcriptomics data and kinetic multi-dimensional (2D or 3D spatial and 4D spatiotemporal) spatiotemporal modeling. As a result, the field currently relies on descriptive approaches that are limited in their ability to generate predictions and uncover mechanisms in complex dynamic tissues. Thus one of the biggest challenges now is how to best leverage the advantage of next generation spatial transcriptome techniques such as Stereo-seq and related approaches to build an analytical framework to achieve single cell resolution and 3D spatial transcriptomics modeling over time and space.

To address these unmet needs, here we introduce Spateo, a modular computational platform providing a comprehensive environment for end-to-end analysis of single-cell resolution spatial transcriptomic data that connects molecular observations to two-dimensional patterns and three-dimensional dynamics through advanced spatiotemporal modeling, and demonstrate its integration with next generation spatial transcriptome techniques and application in four different tissues to extract biological insights. Overall, Spateo delivers four major innovations (**Fig. 1, Supplementary Table 1**). First, Spateo identifies spatial polarity/gradient genes (e.g. neuronal layer specific genes) by solving a partial differential equation to digitize layers and columns of a spatial domain. Second, Spateo implements a full suite of spatially-aware modules for differential expression inference, including novel parametric models for spatially-informed prediction of cell-cell interactions and interpretable estimation of downstream effects. Third, Spateo enables reconstruction of 3D whole-organ models from 2D slices, identifying different “organogenesis modes” (patterns of cell migration during organogenesis) for each organ and quantifying morphometric properties (such as organ surface area, volume, length and cell density) over time. Fourth, Spateo brings in the concept of the “morphometric vector field” that predicts migration paths for each cell within an organ in a 3D fashion and reveals principles of cell migration by exploring various differential geometry quantities.

**Figure 1.**
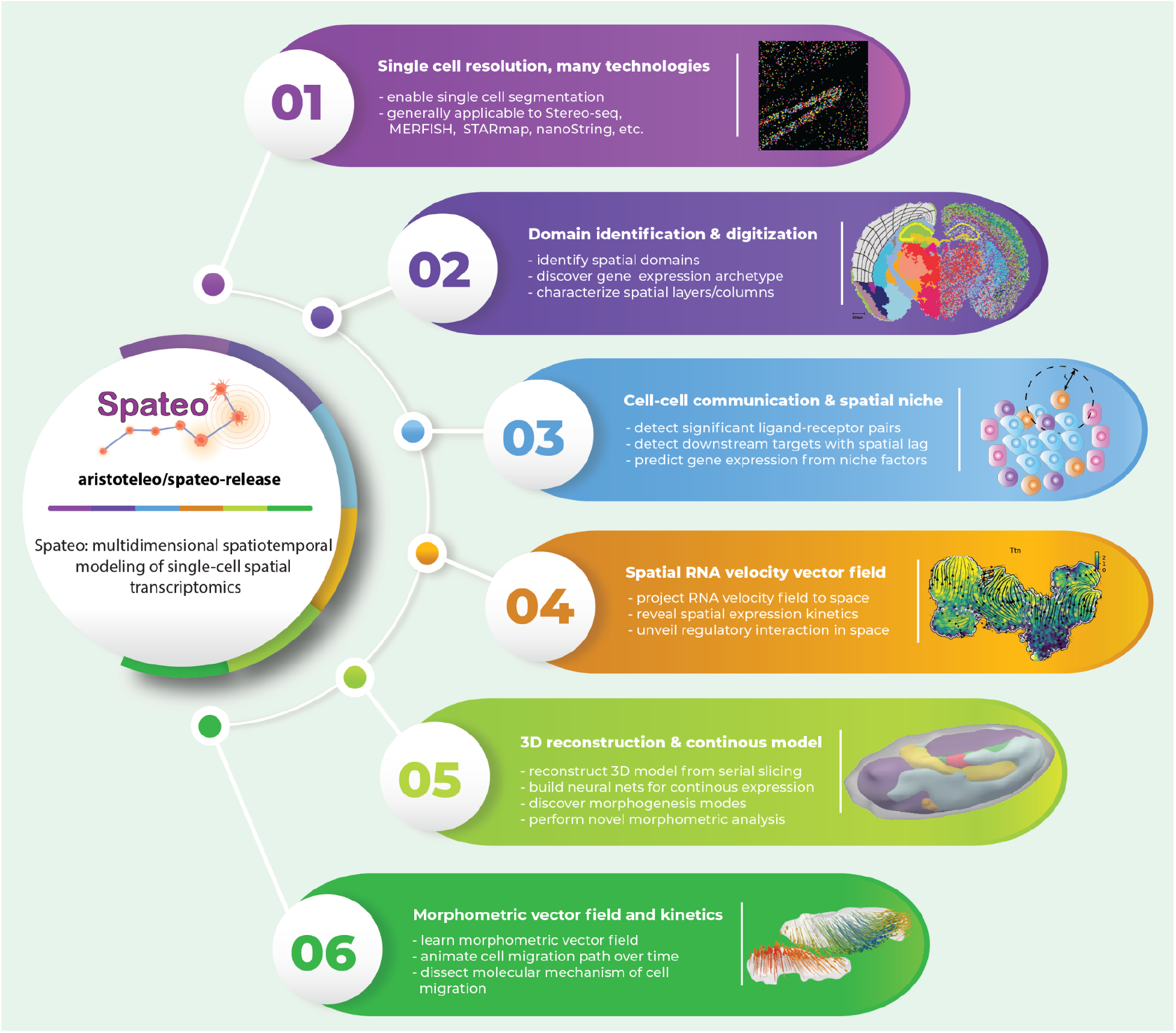
Spateo: advanced spatiotemporal modeling framework of single cell resolution spatial transcriptomics in 3D. **a**) Spateo is a novel computational framework tailored for single cell resolution spatial transcriptomics that enables single cell segmentation based on RNA staining fluorescence or RNA signals, and has general supports to Stereo-seq, MERFISH, seqFISH, STARMap, Nanostring, and others. **b**). Spateo identifies continuous spatial domains, dissects spatial cell type distribution, discovers gene expression archetypes and characterizes spatial polarity genes by digitizing spatial layers or columns. **c**). Spateo models cell-cell communication and spatial niches via a general spatial lag regression approach to detect significant ligand-receptor interactions, unveil signal cascade and predict spatial niche effects. **d**). Spateo enables spatial RNA velocity vector field analyses to reveal spatial expression kinetics and regulations. **e**). Spateo reconstructs 3D models of organs and embryos, builds continuous expression models using a neural network, discovers various morphometric modes and performs sophisticated morphometric analyses. **f**). Spateo learns morphometric vector fields, predicts cell migration paths, and relates morphometric properties to the underlying regulatory genes.

Spateo facilitates a shift in single-cell analysis, from the conventional, reductionist cell-centric focus to embracing the tissue as a whole, allowing ultra-fine, multi-dimensional spatial and temporal examination of molecular mechanisms. Spateo is generally applicable to any sequencing-based or imaging-based spatial transcriptomic readouts and can be used in conjunction with **dynamo** (Qiu et al., 2022), a general framework for RNA velocity vector field analyses, to enable quantitative and predictive analyses of spatiotemporal kinetics of cell fate transitions. Extensive tutorials, workflows and documentation are provided at https://spateo-release.readthedocs.io/en/latest/, and the open-source toolkit can be found at https://github.com/aristoteleo/spateo-release, where community contributions are welcome.

## Results

### Spateo dissects the architecture and composition of tissues domains, identifying domain-specific and spatial polarity genes

A full understanding of tissue architecture depends on the ability to delineate spatial domains (e.g., corresponding to anatomical or histological regions of interest), and to characterize spatial dependencies in gene expression and cell type distribution. Accordingly,we introduce the spatially-constrained clustering (SCC) algorithm which identifies continuous spatial domains using both gene expression similarity and spatial proximity **(STAR Methods)**. To quantify the spatial cell type composition, we developed a cell segmentation method (Starro) that has been integrated as part of Spateo (see our [Cell segmentationwebpage](https://spateo-release.readthedocs.io/en/latest/tutorials/notebooks/cell_segmentation.html), will be reported elsewhere), which segments single cells based on RNA intensity. These segmented single cells can then be clustered using the aggregated expression matrix from the segmentations, annotated based on cluster-specific markers, and finally projected back to the physical space to reveal the spatial distribution of distinct cell types.

Simultaneous spatial domain and cell type annotations enable a broad range of novel analyses, including: first, investigating the contribution of each cell type to different domains and *vice versa;* second, analyzing the spectrum of spatial enrichment or dispersion of different cell types; third, identifying genes or gene archetypes (clusters of genes whose expression shows characteristic spatial distribution) that are specifically enriched in a particular spatial domain; fourth, studying how cell types colocalize and interact with each other; fifth, computationally defining arbitrarily-shaped spatial layers or columns of a tissue domain; and lastly discovering *“polarity genes”* that change along different layers or columns.

The mouse adult coronal hemibrain consists of a complex array of highly specialized neuronal cell types that forms characteristic domains (Narayanan et al., 2017; Radnikow and Feldmeyer, 2018) with distinct functionalities (e.g., the striatum is responsible for regulating motor-functions and reward behaviors; and the ventricle mainly cushions and protects the brain, etc.), thus providing an ideal system to develop, benchmark and test algorithms for the aforementioned tasks. Using Stereo-seq, we obtained a spatial transcriptomic dataset sampled from a mouse adult coronal brain section. We first implemented SCC to identify 18 continuous spatial domains (**Fig. 2a, Fig. S1a, b**, see **STAR Methods**). We next segmented cells based on RNA signal (**Fig. S1c**), resulting in 11,854 cells with about 400 genes and 600 UMIs per cell, sufficient for cell type identification (**Fig. S1d**). We identified a total of 26 cell types, each characterized by specific markers, for example *Slc1a3, Mag, Slc17a7* and *Gad1* for astrocytes (AST), oligodendrocytes (OLIG), excitatory neurons (EX), and inhibitory neurons (IN), respectively (**Fig. 2b,c**). Compared to the Louvain algorithm and SpaGCN, SCC outperforms the Louvain algorithm and performs comparably to SpaGCN in obtaining continuous spatial domains (**Fig. S1a**).

**Figure 2.**
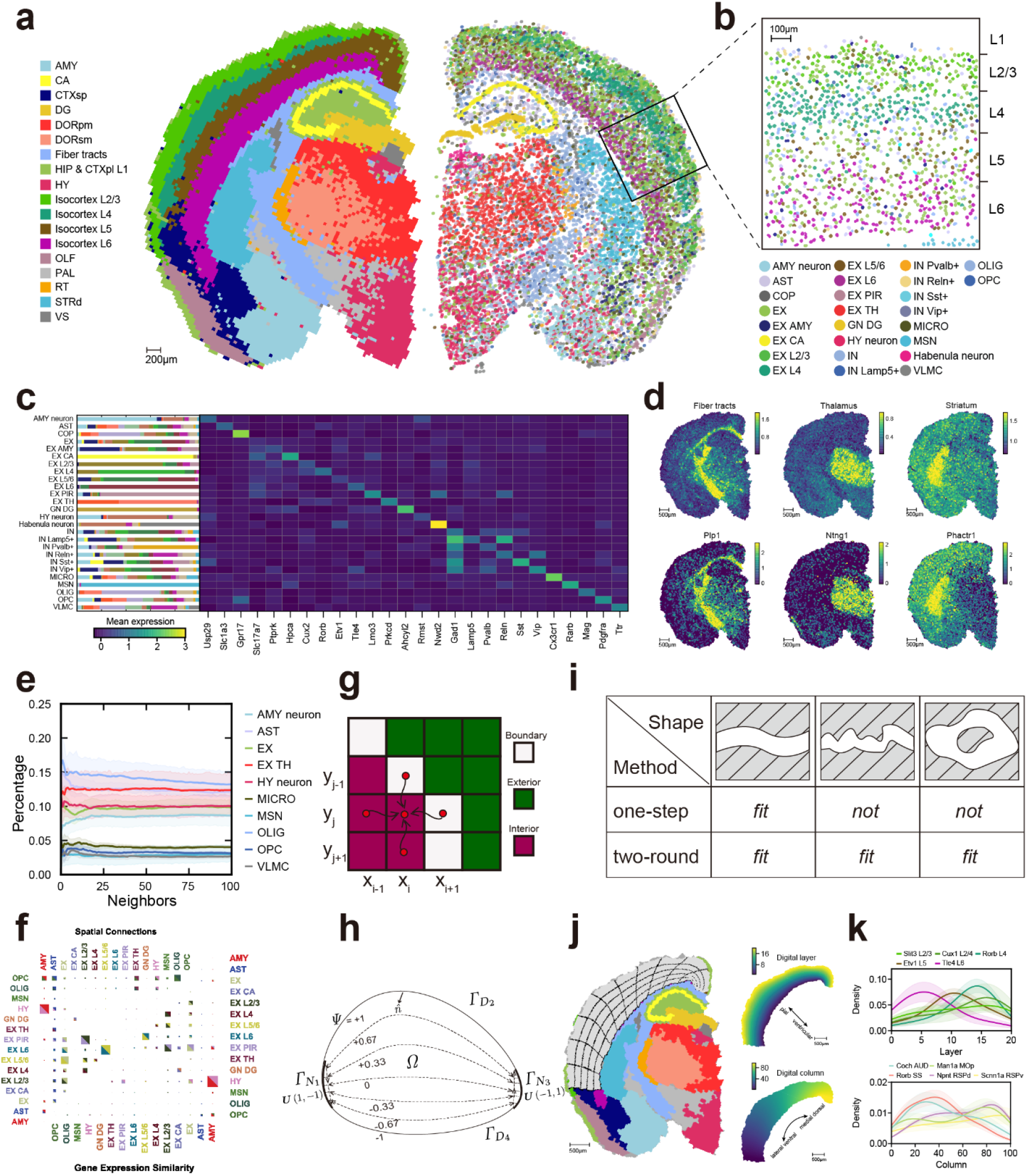
Spateo harmonizes spatial domain and cell type characterization, and unveils expression polarity across spatial layers / columns. **a**). Spatial domains and cell types of the mouse adult coronal hemibrain section. **Left**: the spatial domains identified by the unsupervised SCC method at bin 60 (30 um^2^) resolution. **Right**: cell types annotated based on the clustering result from the segmented cells. Cell shape corresponds to the cell segmentation. The legend indicates the mapping from each color to the corresponding spatial domain or cell type. Same as in Panels **b**, **c, e, and f**. AMY: Amygdalar nucleus; CA: Cornu ammonis; CTXsp: Cortical subplate; DG: Dentate gyrus; DORpm: Thalamus, polymodal association cortex related; DORsm: Thalamus, sensory-motor cortex related; HIP & CTXpl L1: Hippocampus and Cortical plate layer 1; HY: Hypothalamus; OLF: Olfactory area; PAL: Pallidum; RT: Reticular nucleus. STRd: Striatum dorsal region; VS: Ventricular systems. **b**). Magnification inlet showing spatial composition of different cell types within different neuronal layers in the regions squared in Panel **a**. **c**). **Left**: the spatial domain contribution of each cell type. **Right**: marker expression for each cell type. **d**). Identified archetypes (**top**), and corresponding example marker (**bottom**) gene expression in space. **e**). Spatial co-localization relationship between different cell types. **Left**: Pairwise connectivity plot of identified cell types. *Top triangle:* spatial connectivity; *bottom triangle:* expression connectivity. Box size corresponds to the strength of connectivity. **Right**: The percentage of neighbors from OPCs belonging to specific cell types as a function of neighbor size. The shade of each line corresponds to the *90*% confidence interval. **f**). Heatmap of spatial connection and gene expression similarity. Left and top axes display the identified cell types of axolotl brain at 2-DPI. Top-left triangle: cell-type colocalization inferred using actual spatial proximity. Bottom-right triangle: cell-type transcriptomic similarity. Box sizes correspond to the number of spatial colocalization or strength of expression similarity for the top-left and bottom-right triangles, respectively. **g**). The schematic of the Jacobi method (Saad, 2003), showing that the potential of the central grid point is the equal-weighted average of four neighborhoods. **h**). The schematic of the solution of domain digitization boundary problem for irregular bounded domains. See **STAR Methods** for further details. **i**). The schematic of the solution of domain digitization boundary problem for regular/smooth bounded domains. See **STAR Methods** for further details. **j**). Digitization of the neuronal layers (L2-L6) into different layers and columns. **Left**: the select spatial domain in gray color and example borderlines showing the result of region digitization. **Right**: the assignment of layers and columns. **k**). Example genes showing distinct layer and column dependent (or polarity-like) expression. Expression values are firstly density normalized using the *kdeplot* function from *Seaborn* (*Waskom, 2021*). The shade of each line corresponds to the *90%* confidence interval. Celltype abbreviations: (*Domain*) neuron: *Domain-enriched* neuron subtypes; AST: Astrocyte; COP: Differentiation-committed oligodendrocyte precursors; EX (*Domain):* Excitatory neuron or *Domain*-enriched subtypes; GN DG: Granule cells enriched in dentate gyrus; IN (*Gene*+): Inhibitory neuron or *Gene*-expressed subtypes; MICRO: Microglia; MSN: Medium spiny neuron; VLMC: vascular and leptomeningeal cell; OLIG: Oligodendrocyte; OPC: Oligodendrocyte precursor

To investigate the interaction landscape of spatial domain and cell type, we computed the cell type composition of each spatial domain (**Fig. 2c, Fig. S1e**). We found spatial domains characterized by enrichment of specific cells. For example, the isocortex domain is enriched with *Slc17a7*-expressing excitatory neurons, while a *Cux2*-expressing intratelencephalic subpopulation is enriched in the subcortical layer L2/3 domain (**Fig. 2c**). We identified spatial domain-specific genes and gene expression archetypes. Analysis of the fiber tract, thalamus and striatum archetypes reveal different marker genes (*Plp1, Ntng1*, and *Phactr1*, respectively) (**Fig. 2d**). Gene ontology (GO) enrichment analyses of associated genes of identified archetypes are enriched in nervous system development related biological processes (BP), such as myelination (archetypes 11, fiber tract related) and dendrite development (archetypes 5, striatum related) (**Fig. S1f**). To analyze the spatial distribution of each cell type, we used Moran’s *I (Moran, 1950*), a quantification of the tendency of nearby samples to have similar values, to identify cell types that are spatially enriched or dispersed (**Fig. S1g**). We found cell types such as dentate gyrus granule cells (GN DG) show strong spatial enrichment while other cell types, such as *Pvalb*+ inhibitory neurons and microglia, are more dispersed (**Fig. S1h**).

In addition to spatial-aware clustering, Spateo is able to use spatial information to describe additional aspects of tissue architecture, such as cell-type to cell-type colocalization and spatially-dependent gene expression variation along arbitrary axes, constituting means of identifying the most interesting patterns for downstream analyses, e.g. specific cell type pairs that can be used for cell-cell communication analyses or genes that may be important for domain-specific functions. Using the cell-by-cell adjacency matrix and cell-type identity matrix to quantify cell type colocalization (**STAR Methods**), we found that OLIG cells strongly colocalized with oligodendrocyte progenitor cells (OPCs) across different distance scales (**Fig. 2e, Fig. S1i**), which is consistent with the fact that OPCs are potential progenitors for OLIG cells. To define the spatial columns and layers of a spatial domain with any arbitrary shape, we used a digitalization approach (**Fig. 2g-i**, **STAR Methods**) based on solving a potential equation to partition the neuronal cortex into 20 layers and 100 columns and identified genes that significantly change along different layers or columns (**Fig. 2j, Fig. S1j**). For example, we found peak expression of known coronal markers *Tle4* (Isocortex layer 4), *Etv1* (L5), *Rorb* (L6) at digital layers 5, 10 and 15, respectively. We also detected *Coch, Rorb, Man1a, Npnt, Scnn1a* at the lateral, dorsal and medial columns, respectively, consistent with the known order of these markers for cortical functional areas (auditory area, somatosensory area, motor area, retrosplenial area) (Weed et al., 2019) (**Fig. 2k**); these column-specific genes are difficult to identify with other spatially varying gene detection methods (Ma et al., 2022).

In summary, the above analyses demonstrate Spateo’s power in revealing spatial domains, analyzing cell type compositions, characterizing gene expression archetypes and finding spatial polarity genes via spatial domain digitization, each of which is generally applicable to many spatial transcriptomics technologies(Cho et al., 2021; Eng et al., 2019; Stickels et al., 2021; Wang et al., 2018; Xia et al., 2019) and biological systems.

### Spateo predicts cell-cell interactions and characterizes potential downstream effects of intercellular signaling

Tissue phenotypes and intercellular dynamics are shaped by the collective signaling events that occur among constituent cells in spatial microenvironments (Baccin et al., 2020; McCarthy et al.,2020; Rodda et al., 2018); these events are crucial to understanding cellular behavior over 3D space and across time. However, available tools for cell-cell communication analysis are limited in key aspects; for example some methods do not consider spatial distance (Browaeys et al.,2020; Efremova et al., 2020; Wilk et al., 2022), are restricted to potential limited ligand and receptor repertoires known from experimental evidence (Cang and Nie, 2020; Efremova et al.,2020; Wilk et al., 2022; Yuan and Bar-Joseph, 2020), and do not measure cellular communication (Yuan and Bar-Joseph, 2020). Furthermore, methods that characterize potential interaction effects provide limited insight on the cells or cell types involved (Arnol et al., 2019;Browaeys et al., 2020). In Spateo, we unify spatial information and existing statistical approaches for identifying ligand-receptor (L-R) interactions, and apply novel spatial-aware regression methods to model transcriptomic data and reveal potential effects of interaction (**Fig. 3a**).

**Figure 3.**
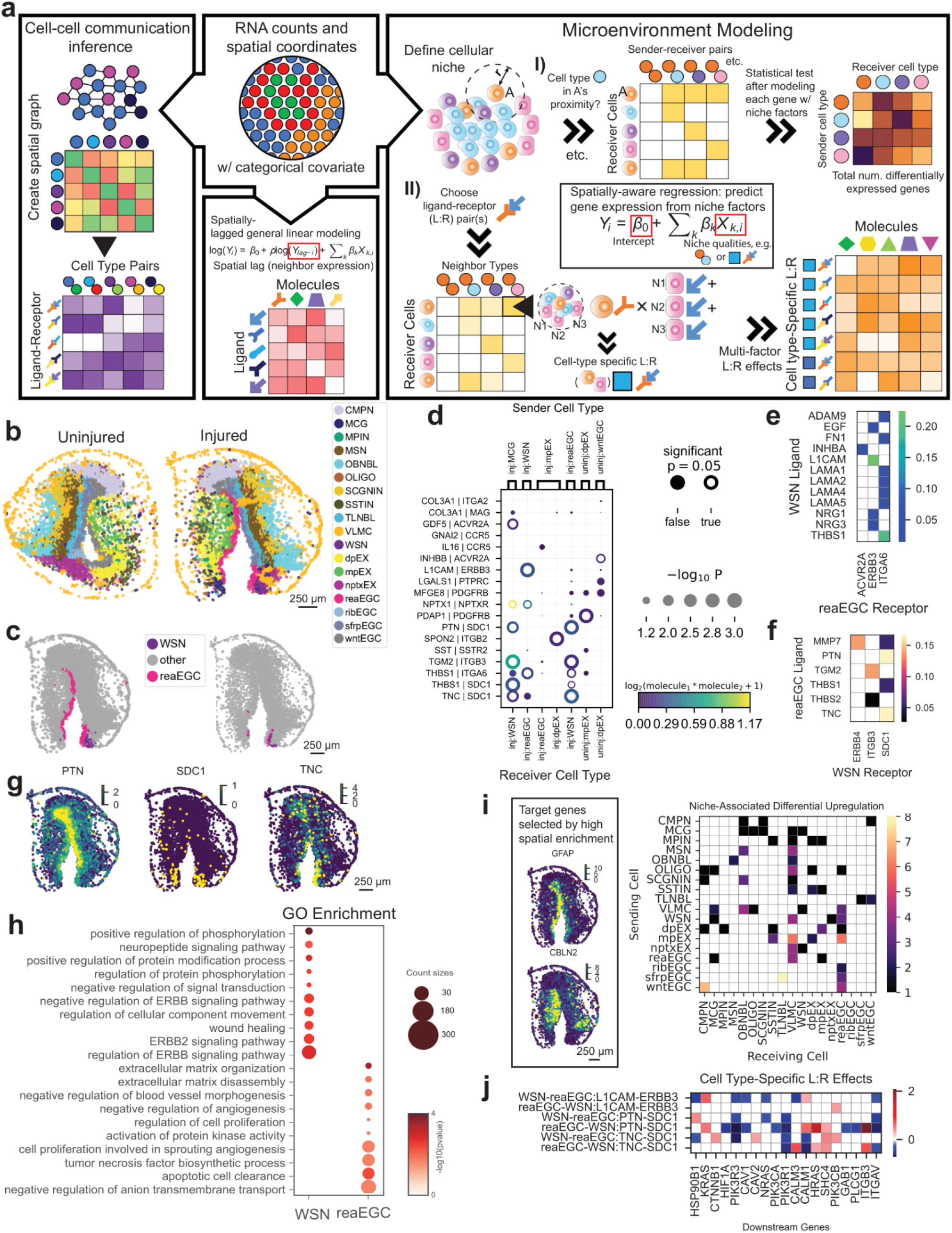
Spateo models intercellular communication and estimates associations with ligand-receptor interaction events. **a**) Scheme of Spateo’s cell-cell communication and microenvironment modeling core. From a spatial transcriptomic dataset with single cell resolution that has been annotated with categorical group labels, Spateo implements cell-cell interaction prediction based on ligand-receptor products conditioned on spatial proximity (**left**). Spateo’s regression suite enables estimation of ligand effects on expression of receptors and other molecules of interest using spatially-lagged (considering dependent variable magnitude in neighboring cells) general log-linear modeling (**middle bottom**). Spateo is also able to estimate the impact of niche factors (**right box, top row**) and cell type-specific ligand-receptor interactions (**right box, bottom row**) on expression of molecules of interest with spatially-aware generalized linear modeling that enables selection of multiple non-normal distribution assumptions. **b**) Cell types of cross-section of both hemispheres of the axolotl telencephalon, two days post-injury where only one hemisphere is subjected to injury. reaEGC: reactive ependymoglial cell; sfrpEGC: *SFRP+* ependymoglial cell; wntEGC: *WNT+* ependymoglial cell; WSN: wound-stimulated neuron; mpEX: medial pallium excitatory neuron; dpEX: dorsal pallium excitatory neuron; nptxEX: *NPTX+* lateral pallium excitatory neuron; VLMC: vascular leptomeningeal cell; MCG: microglia. Cell type labels used here and in all following figures. **c**) Spatial distribution of reaEGCs and WSNs (**left**) and spatial distribution of the same cell types at only the locations where the two cell types are assumed to be in range for interactions, as defined by the nearest-neighbors graph (**right**). **d**) Most significant interactions on both injured and uninjured sides of the telencephalic hemisphere. From all interactions involving wntEGC, nptxEX, mpEX and dpEX in the uninjured side and reaEGC, WSN, mpEX, dpEX, MCG in the injured side, those in which the average ligand-receptor product is not >0.1 for at least one cell type pair are filtered to arrive at this subset. Ligand-receptor pair on left axis, sender and receiver cell types on bottom and top axes, respectively, corrected *p*-value denoted by circle size, ligand-receptor product by circle color, and significance by an open or filled circle. **e**) All identified nonzero interactions resulting from paired testing of WSN-produced ligands and reaEGC receptors, using the method described in subsection **d**. **f**) Subset of identified nonzero interactions resulting from paired testing of reaEGC-produced ligands and WSN receptors, using the method described in subsection **d**. **g**) Expression patterns for *PTN, TNC* and *SDC1*. These ligands (*PTN, TNC*) and the cognate receptor (*SDC1*) colocalize along the reaEGC-WSN interface. **h**) For reaEGC ligands and WSN receptors from **f**, significantly-enriched gene ontology (GO) biological processes (BP). **i**) From outputs of the model labeled **I** in the right box of panel **a,**number of genes that are both differentially expressed and positively associated with proximity of a given pair of cell types (labeled as “differentially upregulated”), with neighboring (sender) cell types along the left axis and receiver (refer to annotation “A” in the “Microenvironment Modeling” box in panel **a**) cell types along the bottom axis, with differential expression assessed by significance testing. Gene set contains all genes with nonzero expression in >5% of cells in addition to Moran’s *I* coefficient that is significant and >0.4, with expression patterns for example genes *GFAP* and *CBLN2* in the left subpanel. The same gene set is used for **Fig. S3. j**) From outputs of the model labeled **II** in the right box of panel **a**, estimated effect sizes when cell type-specific patterns (specifically, reaEGC-WSN and WSN-reaEGC specific patterns) of ligand-receptor interaction are used to predict the expression patterns of select downstream genes. A prior knowledge network database was used to choose genes downstream of *ERBB3* and *SDC1* (see **SLICE model** in **STAR Methods**).

As intercellular communication is distance-limited (Francis and Palsson, 1997), to reduce false positives, Spateo considers spatial proximity for paired cell types by sifting both cell types whose corresponding cells are in each others’ local neighborhoods and determining likely interactions in a data-driven manner (see **STAR Methods**). Phenotypic changes can be induced by the action of subsequentially coordinated gene expression changes downstream of intercellular interaction events. These intercellular dependencies can be attributed to L-R interactions or other diverse molecular mechanisms, such as direct exosomal delivery (Mathieu et al., 2019). To predict in an unbiased manner the downstream effects of intercellular interaction on gene expression, we introduce the niche regression model (model **I** in **Fig. 3a, right panel, Fig. S3a**, see **STAR Methods** for details), which aims to identify genes with altered expression where particular cell type pairs colocalize (henceforth, this will be described as “niche regression”). Although useful to infer potential effects of cellular communication, the niche regression model cannot inform which ligand(s) and receptor(s) caused the observed expression. To address this need, we introduce Spatially-aware Ligand:receptor-based Inference of Cell type-specific Effects (SLICE) (model **II** in **Fig. 3a, right panel**, see **STAR Methods** for details), a spatially-lagged regression approach essentially aiming to identify altered gene expression at locations where particular cell type pairs both colocalize and express specific combinations of ligand and receptor.

We utilized each of these tools to comprehensively characterize the interaction landscapes in the axolotl brain, in the context of the remarkable nervous system regeneration that can occur in these amphibians (Amamoto et al., 2016). Time course Stereo-seq was previously used to collect samples throughout this process, identifying neural-stem-like ependymoglial cell (EGC) states that may mediate this process (Wei et al., 2022) (**Fig. S2a**). We used Spateo to describe the interaction landscape of one such state, reactive EGCs (reaEGCs). Eighteen cell types were found to exist two days post-injury (**Fig. 3b, Fig. S2b-d**), including reaEGCs and an emergent population of adjacent, immature wound-stimulated neurons (WSNs) appearing only in the injured hemisphere (**Fig. 3c**). Several additional cell types could be found in proximity to reaEGCs, including medial- and dorsal-pallium excitatory neurons (mpEX and dpEX, respectively), *WNT+* EGCs (wntEGC), and microglia (MCG), constituting clusters potentially capable of inducing or promoting regenerative processes through cell-cell communications (**Fig. S2e**).

We first derived enriched ligand-receptor interactions for paired cell types, finding that many of the significantly predicted interactions for the injured hemisphere involved either reaEGCs or WSNs, although many cell types were predicted to be involved as either ligands or receptors (**Fig. 3d**). Although the interaction landscape was heterogeneous over time as various cell types changed in prevalence (**Fig. S2g-h**), for the injured hemisphere two days post-injury, pleiotrophin (*PTN*)-syndecan 1 (*SDC1*), tenascin C (*TNC)-SDC1*, and L1 cell adhesion molecule (*L1CAM)-ERBB3* emerged as the most notable interactions, with ligands enriched in reaEGCs and receptors enriched in WSNs. Matrix metalloproteinase-7 (*MMP7)-SDC1* emerged as a sparser but highly-specific interaction between neighboring reaEGCs and WSNs (**Fig. 3d-f Fig. S2f**). In addition to involvement in known ontogenic and regenerative processes(Faissner et al.,2017; Iseki et al., 2002; Mouthon et al., 2020; Rojas-Muñoz et al., 2009), these molecules are enriched in reaEGCs, WSNs and adjacent radial glia, constituting a potential mutually activating relationship between reaEGC and neural progenitors. Notably, expression of *PTN, TNC* and *SDC1* in reaEGCs (**Fig. S6b**) suggests a unique pleiotropic effect in actively proliferating as well as promoting differentiation and mitogenesis in surrounding cells in response to traumatic brain injury (**Fig. 3g**). Having identified multiple potential ligand-receptor mechanisms, we fitted niche regression models and SLICE models to identify potential downstream effects of cellular interaction unconstrained by particular molecular pairs and specific to the *PTN-SDC1, TNC-SDC1* and *L1CAM-ERBB3* interactions, respectively. From our niche regression models, we queried the effect on gene expression of cell type composition in the local neighborhood of each cell across the tissue, revealing the positive effect of WSNs on regenerative factors such as *ATF3 (Seijffers et al., 2007*) and vimentin (Perlson et al., 2005) in reaEGCs (**Fig. S3b**) and the pervasive influence of fibroblastic vascular leptomeningeal cells (VLMCs), perhaps attributable to the fundamental importance of extracellular matrix interaction to growth, proliferation, differentiation and response to signals (Chaudhuri et al., 2020; Discher et al., 2005) (**Fig. 3i, Fig. S3f-g**). We also revealed a potential axis of communication between reaEGCs and other radial glial cells (sfrpEGC, wntEGC), with the upregulation of *WNT*-pathway modulator *SFRP1* at the reaEGC-sfrpEGC/wntEGC interface lending support to the hypothesized origin of reaEGCs from the latter two cell populations(Wei et al., 2022). SLICE models revealed an upregulation of integrins and growth-promoting, axonogenic molecules such as *HRAS (Fivaz et al., 2008*) (**Fig. 3j**) in association with significant cell type-specific signaling (**Fig. S6**); previously characterized biological relationships between *PTN* and identified molecules(Himburg et al., 2014) lend support to the hypothetical effect of *PTN* signaling on growth and differentiation.

Spateo’s spatially-aware models offer a flexible framework to connect gene expression patterns to cell-cell interaction. Any number of genes of interest can be modeled with either the niche regression or SLICE models, enabling identification of putative mechanisms driving pathways hypothesized to have some effect on the phenotype, e.g. glial differentiation, Wnt signaling, and upregulation of translation in the case of the injured brain (**Fig. S3b-e, Fig. S4**), or, starting from all genes measured, enabling unbiased identification of genes strongly influenced by the local microenvironment. Applied to the axolotl sample, the latter approach revealed genes highly dependent on proximity of other sender cells to reaEGCs, including many genes nearly completely polarized along the interface with another cell type (**Fig. S5**). Using Spateo’s cell-cell interaction inference and gene expression modeling capabilities, we identified robust regeneration signatures within the wounded axolotl telencephalon, related these signatures to one another in the context of the physical space, and demonstrated Spateo’s broad applicability to numerous questions involving intercellular interactions.

### Spateo reveals spatially dependent gene regulation via RNA velocity vector field analyses

RNA velocity vector field analyses are powerful tools to explore RNA kinetics and underlying gene regulatory networks during dynamical biological processes (Qiu et al., 2022). However, RNA velocity has been difficult to calculate from previous spatial transcriptomic data due to their low spatial resolution and poor RNA capture efficiency (Asp et al., 2019). With the high RNA capture and high field-of-view of single-cell-resolved next generation spatial transcriptomic techniques, such as Stereo-seq, it is possible to perform single cell RNA velocity vector field analyses at the scale of an organ.

The heart, one of the first organs to appear during mammalian embryogenesis, is a highly structured organ which consists of a relatively small number of cell types, making it suitable for comprehensive RNA velocity vector field characterization. Using Stereo-seq, we previously profiled mice embryos that sampled cells from the heart extensively from E9.5 (embryonic day 9.5) to E16.5 in (Chen et al., 2022).

We used this dataset to investigate the spatial distribution of cardiac cell types over time and space and performed RNA velocity vector field analyses based on the intron and exon information of the mouse heart during mouse organogenesis (**Fig. S7a**).

We used three cardiac specific markers (*Myl2, Myl7, Actc1*) (Chen et al., 2022) to identify the region corresponding to the heart on each embryo slice and then manually extracted the cardiac cells from these regions (**Fig. S7b**). Next we leveraged Starro to segment single cells in these regions using RNA intensity. The segmentation resulted in a spatially-resolved heart cell atlas of 21,075 cells from a total of 16 slides from E9.5 to E16.5, collected at one day intervals. After preprocessing, clustering and marker calling, we identified a total of ten cell types, including cardiomyocyte progenitor cells (CMPs) and several cardiomyocyte subtypes (CMSs), smooth muscle cells, epicardical / fibroblast cells, macrophages and others (**Fig 4a-c and STAR Methods**). We found strong spatial as well as temporal enrichment of these cell types. For example, CMPs are enriched at earlier time points (E9.5-E12.5) and epicardical cells are enriched only in the outer-layer of the heart, surrounding the CMSs (**Fig. 4a**). Furthermore, the identified region-specific CMSs, such as atrial cardiomyocytes and ventricle cardiomyocytes, are characterized by spatially-restricted expression of *Nppa* and *Ckm*, respectively (**Fig. 4c, Fig. S7c, d**) (Litviňuková et al., 2020). We next performed spatially resolved RNA velocity vector field analyses of the maturation of either cardiomyocytes, or that of the fibroblast cells from epicardial cells (Meilhac and Buckingham, 2018) (**Fig. 4d-h, Fig. S7e**). For the cardiomyocytes, this revealed a bifurcation trajectory starting from the CMP cells with low expression of *Ttn* (a marker of mature CMs) (Litviňuková et al., 2020) to either a mixture of ventricular or atrial cardiomyocytes on the left side of the bifurcation or to right ventricular cardiomyocytes on the right side with high expression of *Ttn* (**Fig. 4d**). In order to reveal the spatiotemporal regulation of the cardiomyocyte maturation, we collected a set of 36 cardiac TFs (transcription factors) for RNA Jacobian analysis (Herrmann et al., 2012), to reveal how the increase of a regulator (e.g. a TF) would affect the velocity of a potential target ((Qiu et al., 2022) and see **STAR Methods**). We found, for example, a widespread activation from *Tbx5* to *Gata4* (Herrmann et al., 2012) during the early and middle stages of the cardiomyocyte maturation **(Fig. 4e left)**. Of note, this activation is restricted to the CMs but not present in the surrounding epicardial cells **(Fig. 4 middle)**. On the contrary, regulation from *Tbx5* to *Mef2c* (Herrmann et al., 2012) is positive in the epicardial cells but repressive in the CMs, revealing an interesting inverse pattern of regional specificity of this regulation **(Fig. 4e right)**. The vector field approach enables *in silico* prediction of the effects from genetic perturbations at scale. We therefore investigated the potential resultant cell fate changes following genetic perturbation over space; computationally, this strategy enables genome-wide predictions in spatial data that would otherwise be inaccessible experimentally (Jin et al., 2020) **(Fig. 4f)**. To demonstrate this, we first simulated *Tbx5* knockdown, predicting that knockdown of *Tbx5* would lead to dedifferentiation of mature cardiomyocytes **(Fig. 4f top)**. Secondly, we found that although the expression of both the direct target *Gata4* and the indirect target *Ttn* had repression after *Tbx5* knockout, the prior showed more uniform repression **(Fig. 4f bottom)**. Furthermore, *Tbx5* knockdown seemed to activate the expression of *Gata4* / *Ttn* in epicardial cells, again revealing the heterogeneity of spatial regulation, and implies that *Tbx5* may be an activator in CMs but a repressor in the surrounding epicardial cells **(Fig. 4f bottom)**.

**Fig. 4.**
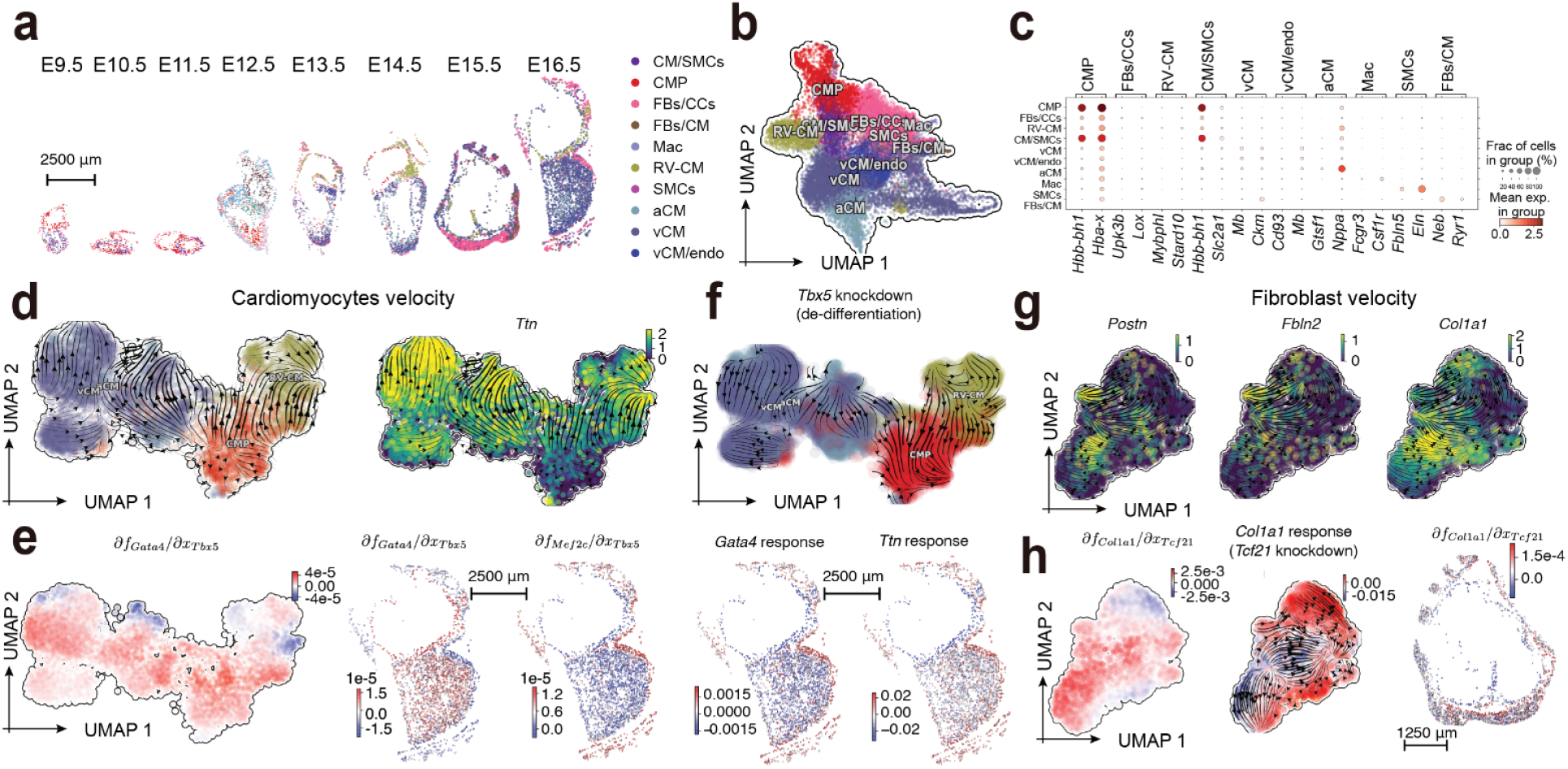
RNA velocity vector field analyses of spatially resolved atlas of cardiac cells during mouse organogenesis. **a**). Spatial geometry plots (geometry here refers to the cell shape, identified through cell segmentation) of distinct cell types of the heart on typical embryo slices from E9.5 to E16.5, profiled with Stereo-seq, from the MOSTA dataset (Chen et al., 2022). CM/SMCs: cardiomyocytes and smooth muscle cells; CMP: cardiomyocyte progenitor cells; FBs/CCs: fibroblast, epicardial/endocardial cells; FBs/CM: fibroblast and cardiomyocyte cells; Mac: macrophage; RV-CM: right ventricular cardiomyocytes; SMCs: smooth muscle cells; aCM: atrial cardiomycotes; vCM: ventricular cardiomyocytes; vCM/endo: ventricular cardiomyocytes and endothelial cells. Color legend of cell type identities is shared between panel **a** and **b**. **b**). UMAP (Uniform Manifold Approximation and Projection for Dimension Reduction) plot of different cardiac cell types from the entire heart cell atlas. **c**). Dotplot of cell-type specific markers. **d**). RNA velocity streamline plot of cardiomyocyte (CM) maturation, with cells colored by the CM subpopulation identities on the **left** or expression of CM marker, *Ttn*, on the **right**. **e**). RNA Jacobian analyses of the cardiomyocyte maturation. From left to right: **i**). The Jacobian from cardiac TF *Tbx5* to another cardiac TF *Gata4* reveals broad activation pattern across various CM maturation states. Cells are shown in the UMAP space and only CM related cells are selected for this analysis; **ii**). The spatial geometry plot reveals this activation is restricted to CM cells but not the surrounding epicardial cells. All cells, excepted those didn’t made to RNA velocity vector field analyses, from the typical E16.5 Stereo-seq slice (also shown in panel **a**) are shown, same as for the spatial plots in the panel **f**; **iii**). Regulation from *Tbx5* to *Mef2c* has a reversal spatial pattern to that from *Tbx5* to *Gata4*. **f**). *In silico* perturbation prediction of *Tbx5* knockout. **Top**): Predicted RNA velocity streamline plot after knocking down *Tbx5*, a key CM maturation regulator. **Bottom left)**: Spatial geometry plot of the predicted response of *Gata4* after knocking out its upstream regulator *Tbx5*. **Bottom right)**: Spatial plot of the predicted response of *Ttn* after knocking out its indirect regulator *Tbx5*. **g**). Same as in panel **d** but for the maturation of fibroblast cells from epicardial cells. Expression of three known fibroblast marker *Postn, Fbln2*, and *Colfa1* in the UMAP space are shown, with RNA velocity streamline overlaid on the top. **h**) RNA velocity vector field analyses of the fibroblast maturation. From left to right: **i**). The Jacobian from epicardial TF *Tcf21* to fibroblast marker *Col1a1* reveals a broad activation pattern across various fibroblast maturation states. Cells are shown in the UMAP space and only fibroblast related cells are shown; **ii**). Scatterplot of the predicted response of *Col1a1* after knocking down the epicardial TF *Tcf21*. **iii**) Spatial geometry plot of the predicted response of *Col1a1* after knocking out *Tcf21*. All cells, excepted those didn’t made to RNA velocity vector field analyses, from the typical E15.5 Stereo-seq slice (also shown in panel **a**) are shown. Scale bars in panels **a, e, f**, and **h** indicates the physical scale of the spatial plots. Cell segmentations are used to draw the cell shape in each spatial plot.

On fibroblast cells, we revealed a linear maturation path from cells with low expression to cells with high expression of *Postn, Fbln2, Col1a1*, known markers of cardiac fibroblast cells (**Fig. 4g**) (Litviňuková et al., 2020). RNA Jacobian analysis of cardiac regulator *Tcf21* (Kanisicak et al.,2016) to *Col1a1* revealed a global activation (**Fig. 4h left**), which is restricted to the surrounding epicardial cells but not to the inner CMs (**Fig. 4h right**). Similarly, *in silico* perturbation also revealed that knockdown of *Tcf21* leads to partial dedifferentiation of mature fibroblasts in the middle fibroblast maturation stage and partial repression of *Col1a1* in relative inner region of the epicardium layer (**Fig. 4h middle, right**). This prediction is in line with other studies showing that *Tcf21* is a key regulator that controls the fate decision between fibroblast cells and smooth muscle cells involved in epicardial cell differentiation (Hu et al., 2020).

Collectively, these analyses showcase how the combination of Spateo with Dynamo’s RNA velocity vector field (Qiu et al., 2022) can reveal spatially-dependent gene regulation during heart maturation, providing important insights into the spatiotemporal control underlying complex cell state transitions.

### Spateo reconstructs 3D *in silico* models and performs morphometric analyses with sequential spatial transcriptomics measurements

The formation of an embryo and constituent organs is characterized by specific “organogenesis modes”, dynamic patterns of organ morphogenesis orchestrated by sequential gene regulatory programs. These gene programs also dictate hierarchical cell fate specification and organization into complex three-dimensional units of structure and function (Olson, 2006). However, technical challenges in penetrating the depth of a tissue or organ for *in situ* sequencing currently prohibit 3D spatial transcriptomics. As a result, existing spatial analyses are generally two-dimensional and static. To fulfill this unmet gap, in Spateo we computationally reconstruct 3D models of embryos and organs using multiple adjacent slices of embryos or organs. Furthermore, going beyond merely aliging and integrating spatial transcriptomics data, as demonstrated in (Zeira et al., 2022), we developed a general 3D modeling framework in Spateo that 1) align cells from multiple embryonic slices in 3D space, 2) build models of such aligned 3D point clouds (cells), 3) analyze the reconstructed 3D models and their constituent organs, 4) identify interesting continuous gene expression patterns in 3D space, and 5) unveil various “organogenesis modes’’ and morphometric kinetics during embryogenesis (see **STAR Methods** for details, **Fig. 5a**, and **Supplementary Animation 1**).

**Fig 5:**
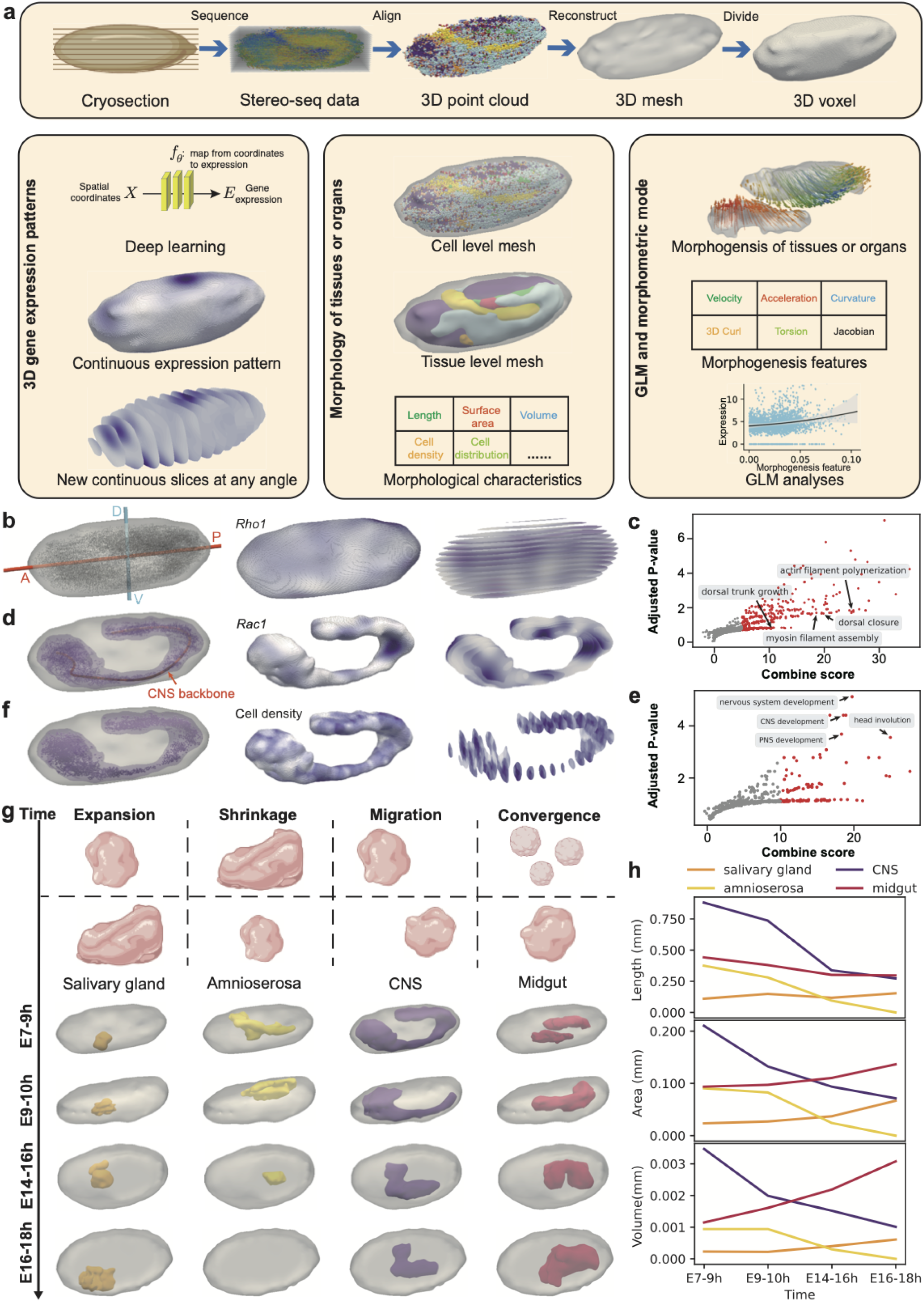
*In silico* 3D reconstruction of continuous Drosophila embryos and organs models reveal axis, organ-dependent genes, spatiotemporal organogenesis modes and morphometric kinetics. **a**). Workflow of 3D tissue/organ/embryo reconstruction and downstream analyses. **b**). A-P (anterior-posterior) or D-V (dorsal-ventral) axis of the embryo (**left**) and significant A-P dependent gene, *Rho1’s* expression pattern on the the surface (**middle**) or along the D-V axis (**right**), shown as the A-P plane, uniformly sliced from the *in silico* embryo model along the D-V axis. Gene expression in the middle and right subpanels of panel **b/d** are retrieved from the smoothed 3D gene expression model learned with the deep learning model (see **STAR Methods**) **c**). GO BP (biological process) enrichment analyses of significant D-V axis variable genes. Scatter plot of combine score vs. adjusted *p*-value of enriched GO terms of all the detected significant D-V axis variable genes. **d**). Same as in **b**, but for the principal curve of the central nervous system (CNS) and the associated significant CNS principal curve dependent gene, *Rac1*. **e**). Same as the top but for the significant principal curve of CNS backbone dependent genes. Key terms are highlighted. **f**) Similar to **d**, but for the cell density of CNS in the middle and right subpanels. For the left panel, the CNS tissue level mesh is embedded in the whole-body embryo mesh. **g**). Four typical organogenesis modes (migration: organs migrate during embryogenesis; shrinkage: organs shrink during embryogenesis; expansion: organs expand during embryogenesis; convergence: multiple disconnected organ primordia converge into a mature organ) and associated example organ during the *Drosophila* embryogenesis from E7-9h to E16-18h.**h**). Morphometric analyses of different organs over time during *Drosophila* embryogenesis. **Top**: Lengths of different organs (the length is based on the fitted principal curve of the organ) over time; **Middle**: same as the above but for the surface area; **Bottom**: same as above but for the volume.

We demonstrate the 3D reconstruction and modeling of Spateo using continuous slices of the whole Drosophila embryo generated for this study **(STAR Methods)** that leverage the large field-of-view of Stereo-seq (Wang et al., 2022). We reconstructed *in silico* 3D models of Drosophila embryos at four stages (E7-9h, E9-10h, E14-16h, and E16-18h), and focused on investigating the 3D gene expression pattern for the E7-9 embryo. We identified principal components of the reconstructed embryo as its A-P or D-V axis in 3D space (**Fig. S8, STAR Methods**). We next performed a GLM regression to identify genes that significantly change along the axis, for example *Rho1*, an actin cytoskeleton organization and morphogenesis related gene, along the D-V axis (**Fig. 5b**). GO enrichment analyses of all the D-V axis-dependent genes revealed strong enrichment of D-V patterning and specification-related pathways (**Fig. 5c**). Similarly, we identified *Rac1* as a gene significantly changing along the principal curve (a curve that passes through the middle of the point clouds of a particular organ in 3D space, see **STAR Methods**) of the central nervous system (CNS). This is in line with *Rac1’s* role in axon outgrowth and cell migration (Luo et al., 1994) (**Fig. 5d**), and the associated GO enrichment analyses revealed many neuronal processes, such as CNS development and head involution (**Fig. 5e**). 3D principal curve-based cell density analysis revealed CNS cells to be most dense at the two ends of the CNS (**Fig. 5f, Supplementary Table 2**).

Our 3D models enabled us to investigate the kinetics underlying “organogenesis modes”. We showcase four general “organogenesis modes”, or dynamic patterns of organ morphogenesis that occur during embryogenesis: 1. organ expansion, primarily driven by cell growth; 2. organ shrinkage, or volumetric shrinkage of an organ driven by cell death; 3. organ migration or movement; and 4. organ fusion by which multiple pieces fuse into a mature organ (**Fig. 5g top**). With Spateo, we were able to identify all these patterns: we found that the salivary gland increases in volume while the amnioserosa decreases and eventually disappears (**Fig. 5g bottom**). On the other hand, the CNS migrates while midgut converges from two separate pieces. Interestingly, morphometric analyses show that although the length of midgut doesn’t change significantly, the area and volume of midgut dramatically increase over time (**Fig. 5h**). Collectively, these fine-grained insights on organ development highlight Spateo’s capabilities to reconstruct 3D *in silico* models of embryos and organs for novel 3D differential gene expression and morphometric analyses, which cannot be properly characterized using static, 2D spatial data.

### Spateo learns morphometric vector fields, and dissects underlying molecular mechanisms of cell migrational curvature, torsion and others

Collecting multiple tissue slices in 3D across several time points allows linkage of dynamic phenotypes to dynamic gene expression patterns, enabling the prediction of cellular migration in an organ over time and the association of genes to movement patterns in events such as morphogenesis. Live imaging provides the opportunity to observe morphogenesis over time at high resolution (McDole et al., 2018), but it cannot associate complex regulatory programs to such morphometric changes because imaging can only measure a few genes within single cells over time. From Spateo’s reconstructed 3D models across different time points, cellular migration patterns can be approximated by constructing “morphometric vector fields” that map cells from an earlier time point to a later time point. Furthermore, potential regulatory mechanisms can be elucidated by leveraging differential geometry analyses of the reconstructed morphometric vector field and the accompanying transcriptomic data.

Here, we repurposed the RNA velocity vector field learning algorithm originally developed in Dynamo (Qiu et al., 2022) to learn a vector field of cell migration in 3D physical space. Instead of the gene expression space, we coupled the morphometric vector field with differential geometry analyses to identify morphogenesis-related genes in the *Drosophila* embryo. We first sampled ~2,000 cells from the E7-9h and E9-10h embryos respectively, then efficiently aligned the embryos based on the sampled cells to place cells from two time points in the same coordinate system. As midgut and CNS show the largest migration changes, we selected them as organs of interest to learn an optimal transport map (see **STAR Methods**) from the E7-9h time point to the E9-10h time point (**Fig. 6a**). We used coordinates from E7-9h cells and optimal transport couplings (visualized as quivers as in **Fig. 6b**) to learn a morphometric vector field using sparseVFC (Ma et al., 2013a). We then used the morphometric vector field to predict the migration trajectory of each cell, revealing convergence of two distinct midgut components into one mature organ (see this link: Atlas of Drosophila Development by Volker Hartenstein(sdbonline.org), **Fig. 6a, 6c**, and **Supplementary Animation 2**).

**Fig 6:**
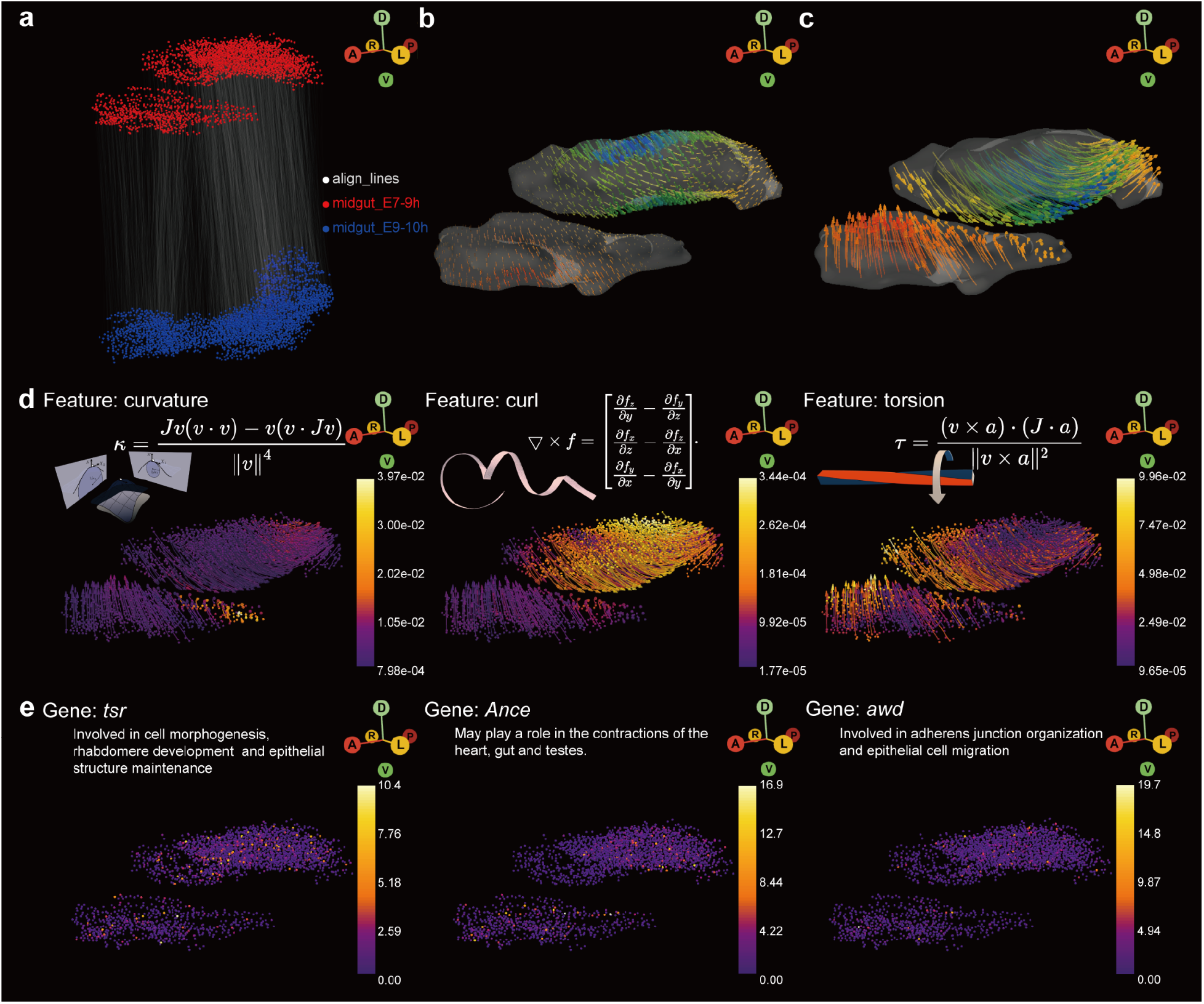
Learning morphometric vector field of Drosophila embryogenesis and unveiling molecular mechanism of morphogenesis via analytical differential geometry analyses. **a**). The alignment between cells of the *Drosophila* midgut from E7-9h to those from E9-10h. **b**). Quivers of the surface of the midgut predicted by the morphometric vector field learned with the cell alignment defined in panel **a**). Quivers are sampled uniformly across the surface. Color of the quiver corresponds to the z component of the velocity vector. **c**). Same as the **b**, but with the predicted trajectories of sample points, visualized as streamlines. **d**). Estimated morphogenic curvature, curl, torsion of the reconstructed morphometric vector field. The cell and streamline are colored by the corresponding norm of each quantity. Definition of each quantity is provided. Note that both curvature and curl are vectors of the *x,y,z* axis, while torsion is a 3 x 3 matrix for each sample point. **e**). Top example gene that is significantly associated with each differential geometry quantity. Short description of the function of each gene that is related to midgut development, or *Drosophila* morphogenesis in general is provided.

For RNA velocity vector fields, analytical differential geometry analyses enable discovery of regulatory mechanisms of cell fate conversion (Qiu et al., 2022). With a 3D morphometric vector field, differential geometry analyses can be analogously applied to the physical space to reveal cellular migration properties. Importantly, for 3D morphometric vector fields, 3D curl, curvature, torsion and others have real physical meanings, and crucial morphogenetic genes can be identified by significant dependence on these morphometric properties. We thus calculated velocity, acceleration and curvature vectors, 3D curl, torsion, divergence, and the Jacobian matrix (**STAR Methods**, **Fig. 6d**, **Fig. S9**) for each cell of the midgut at E7-9h with the learned morphometric vector field. Cells in the top midgut section move down while cells in the bottom section migrate upward, revealing a high degree of twists of the midgut on the boundary regions as revealed by the norm of torsion. We then calculated the norm of all quantities and performed a generalized linear regression, using the norm to identify several differential geometry quantity dependent genes **(Fig. 6e)**. For example, we find that *tsr*, a gene previously known to be involved in cell morphogenesis and rhabdomere development, is associated with curvature — that is the degree that a migration path deviates from a straight line, the curl-related gene *Ance* may play an important role in the construction of the midgut; and torsion-related gene *awd* is a known critical regulator of border cell migration. Applying similar analyses to study the migration of cells in the central nervous system also reveals key regulators of CNS migration, including *chic, Act5C* and *elav*, crucial for brain development, cell motility and muscle contraction and control of gene expression in the developing nervous system (Hilgers et al., 2012), respectively (**Fig. S10, Fig. S11, Supplementary Table 3**).

Collectively, these results highlight how we can go beyond descriptive 2D spatial analyses to more dynamical and predictive 3D spatiotemporal modeling with morphometric vector fields, leading to a paradigm shift in leveraging 3D spatial transcriptomics to reveal regulatory mechanisms of emergent properties. Lastly, we further demonstrated the general applicability of Spateo and applied it to analyze datasets from both *in-situ* profiling-based methods, such as seqFISH, MERFISH and STARmap, and *ex-situ* profiling-based methods such as Slide-seq and seqScope (**Supplementary Figure Fig. S12, Fig. S13**) to gain biological insights of spatial domains, niche effects, cell-cell communication and spatial polar-genes.

## Discussion

It has been over a decade since the advent of single-cell RNA sequencing transformed genomics, empowering the development of experimental and analytical methods capable of profiling and analyzing millions of cells at extraordinary breadth and depth. Now, we are at the dawn of another new era of spatial genomics which is providing an unprecedented opportunity to explore transcriptomic measurements with temporal, spatial and *in vivo* context. Here, we present Spateo, a powerful framework that bridges the gap between observational analysis of static samples and holistic biological understanding considerate of 3D space and time. Similar to other pioneering tools such as Giotto (Dries et al., 2021), stLearn (Pham et al., 2020) and Squidpy (Palla et al., 2022), Spateo is equipped with a complete suite of common preprocessing, visualization and spatial analysis modules, comprising functions for representation of spatial and imaging data, normalization and transformation, de-batching and integration, identification of cell types, associated markers and spatially-variable genes, and informative visualization through interactive plots.

Spateo introduces five major innovations, enabling it to serve as a foundational tool for advanced spatiotemporal modeling: it (1) identifies continuous spatial domains and expression archetypes, (2) Organizes spatial domains into discrete layers and columns to reveal spatially-polar genes, (3) dissects ligand-receptor interactions and predicts interaction-associated effects on gene expression with a spatially-aware regression model, (4) reconstructs 3D embryo and organ models coupled to whole-transcriptome measurements, and (5) learns “morphometric vector fields” that describe cellular migration patterns and facilitate identification of regulatory mechanisms underlying cell movement and morphogenesis. Spateo’s capabilities are made possible by its incorporation of diverse concepts from physics and mathematics, such as partial differential equations, vector fields and differential geometry.

As spatial technologies continue to mature and percolate through biological laboratories, we foresee an explosion of diverse optimization and application of spatial transcriptomics and for further development of Spateo. Many single-cell genomics approaches can be translated to spatial genomics approaches, including single cell multi-omics, RNA metabolic labeling, Perturb-seq and lineage tracing to enable multi-view, spatiotemporally resolved, lineage-resolved and perturbation-resolved cell state dynamics *in situ*. We additionally foresee opportunities to apply Spateo to understand biological systems in many circumstances, for example by generating a spatially-resolved cross-species cell atlases and comparing 3D models of organs between different species to understand the evolutionary emergence of tissue structures, such as the evolution of the four-chamber heart in mammals from the single-chamber heart in invertebrates (Olson, 2006). By further optimizing the 3D morphometric vector field approach, we expect many unique opportunities to directly link regulatory and functional genes with morphological changes at organ and embryo levels.

Spateo complies with the best practice of software engineering, including a modularized and extendable infrastructure that allows continuous optimization, future integration (including **dynamo** (Qiu et al., 2022), **dynast** (https://github.com/aristoteleo/dynast-release) and other frameworks from the Aristotle organization (https://github.com/aristoteleo) and active community contribution to draw from expertise across the field to build a uniform software ecosystem that enables dynamic, quantitative and predictive analyses of single cells and spatial transcriptomics.

### Limitations of the Study

First, we expect that Spateo could be further improved by extending 3D alignment to be compatible with cases where some cells in one slice have no correspondence in the other slice (a partial alignment) or to allow non-rigid alignment across slices because of tissue deformation during sample preparation, as well as to be able to jointly align multiple slices instead of sequentially aligning, which accumulates errors over each alignment. Second, to manage and explore the large and expanding datasets generated by spatial transcriptomics, an interesting prospect would be to draw from Google Earth’s infrastructure to house and interactively interface with spatially-resolved whole-organ 3D atlases. Third, the spatial RNA velocity and the morphometric vector field are currently done separately but could be uniformed into a single learning task to learn reaction-diffusion like spatiotemporal models, and linear regression models used to identify spatial polar genes could be extended to deal with non-linear interactions by repurposing optimal transport or graphical neural network based algorithms to derive communications between signal senders and receivers.

## Supporting information

Supplementary animation 1

Supplementary animation 2

Supplementary animation 3

## ACKNOWLEDGMENTS

We thank Jesse Engreitz (Stanford), members of Jonathan Weissman (Whitehead Institute), Xiao Wang (Broad Institute), Liangcai Gu (UW) and Xiaowei Zhuang’s (Harvard) labs, and Dylan Cable from Fei Chen’s lab (Broad Institute) for discussion and comments on this work. We also sincerely thank various technical support from China National Gene Bank.

## AUTHOR CONTRIBUTIONS

X. Q., Y. B., Shiping. L., J. S. W. and L. L conceived the project. X. Q., Y. B. and D. Y. Z. developed the models and theories. X. Q., D. Y. Z., J. Y., Z. J., L. Z. and J. M., H. P., S. W. developed Spateo, implemented the code, and performed the analyses. M. W., R. X., Z. T., Q. H., and Y. R. L. conducted the Drosophila Stereo-seq experiment, and Sha. L. conducted the mouse brain Stereo-seq experiment. S. H. design the logo for Spateo, under Shiping. L. and X. Q.’s supervision. X. Q., D. Y. Z., J. Y., Z. J., and L. Z. designed and generated the figures. X. Q., D. Y. Z., Y. B., Z. J., J.S.W wrote the manuscript with feedback from all other authors.

## Main figure titles and legends

**PDF version and high resolution png version of all main and supplementary figures can be downloaded from:** https://www.dropbox.com/sh/cvtibiy0yfkpws5/AAArbvcLUBa5vjfqjjgNovRGa?dl=0

**Supplementary Figure 1, related to Fig. 2.**
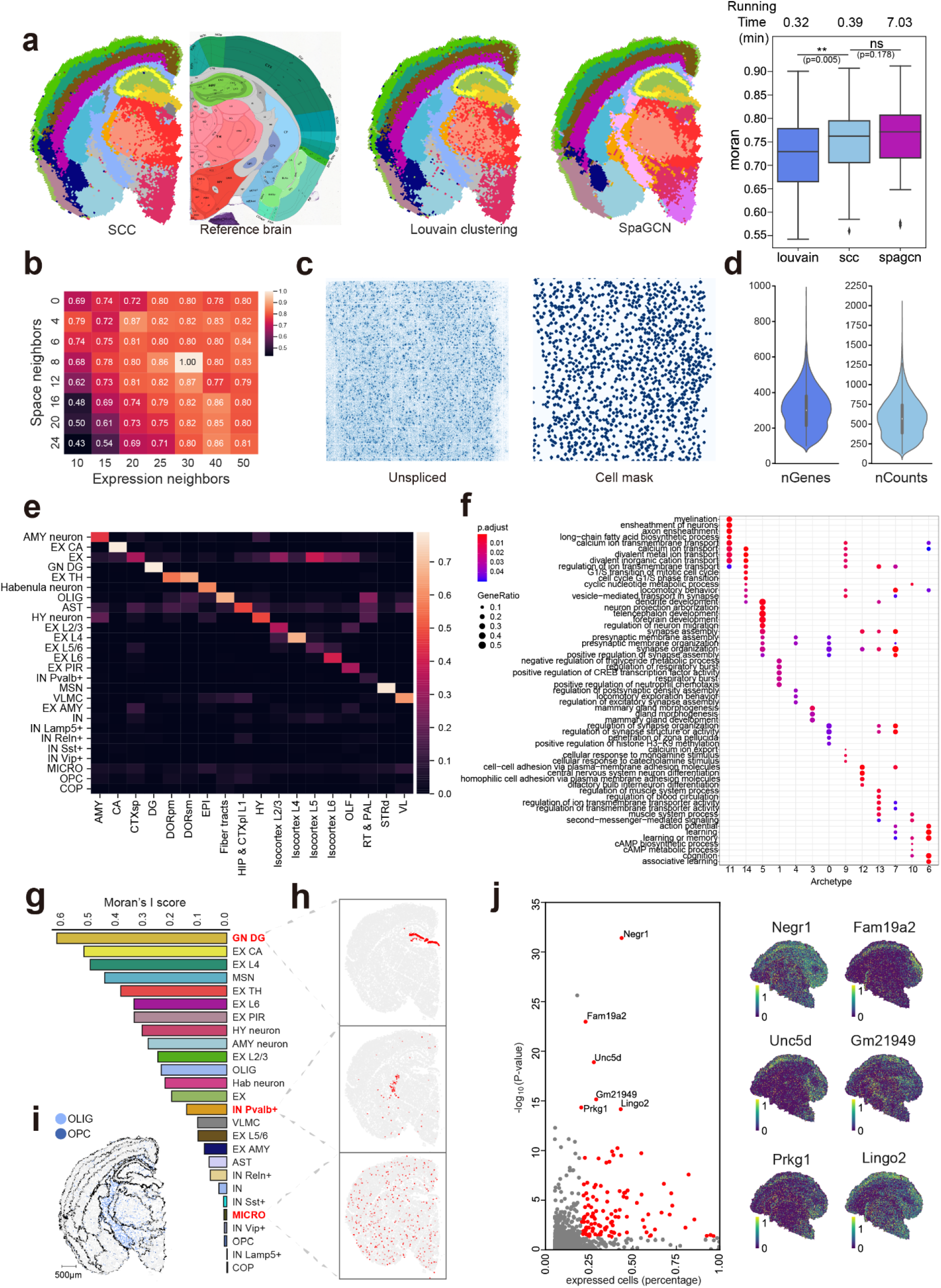
Spateo unravels the relationship between spatial domains and cell types, and identifies polarity genes and their enriched pathways. **a**). Various clustering algorithms reveal similar spatial domains of mouse adult coronal hemibrain section. **First**): Results of SCC (spatially constrained clustering) and Allen brain map reference; **Second**): Results of Louvain clustering; **Third**) Results of SpaGCN. All methods are spatially aware, except the first one. **Fourth**) Running time and distribution of spatial continuity of the resultant clusters for each method. Moran’s I (a measure of spatial autocorrelation) is used to quantify the spatial continuity. **b**). Robustness of SCC under a broad range of spatial or expression neighbors. Color of the heatmap corresponds to the adjusted rand index (ARI) between the clustering result from each setting and the default setting (30 expression neighbors and 8 spatial neighbors). **c**). Spateo leverages high spatial resolution of Stereo-seq to perform RNA signal based single cell segmentation. **Left**): Unspliced RNA intensity (total UMIs in each DNA nanoball); **Right**): Resultant single cell mask after processing unspliced RNA mask. **d**). Distribution of the number of genes and total UMI counts per segmented cell. **e**). The spatial distribution of each cell type in different spatial domains. Colors correspond to the proportion of a particular cell type that is distributed into a particular spatial domain. Columns are ordered based on the most enriched spatial domain of each cell type. **f**). Top enriched GO (gene ontology) BP (biological process) terms associated with gene expression archetypes (a group of genes that show consistent gene expression patterns across cells, see **STAR Methods**). Archetypes are ordered based on the specificity to the top enriched terms on the row. **g-h**). Moran’s *I* statistics stratifies the spatial distribution of distinct cell types. **g**): barplot of the Moran’s *I* score for each cell type. Moran’s *I* is calculated based on the binary vector (close to 1 if the cell comes from a particular cell type, close to 0 otherwise) of each cell type across the space. **h**): Example cell types with high (GN DG), middle (IN Pvalb+) and low (MICRO) Moran’s *I* score, representing cells with high spatial enrichment, adequate spatial enrichment and uniform spatial distribution. **i**) Spatial colocalization of OLIG and OPC. The black lines are the identified borders of the spatial domains. See all significant cell pairs in **Fig. 2f**. **j**). **Left**: Scatterplot of the fraction of expressed cells and *-log_10_(P-value*) based on the generalized linear regression analyses along spatial domain layers or columns (See **STAR Methods**). Highlighted are domain column specific genes, whose spatial expression is visualized in the **right**.

**Supplementary figure 2, related to Fig. 3.**
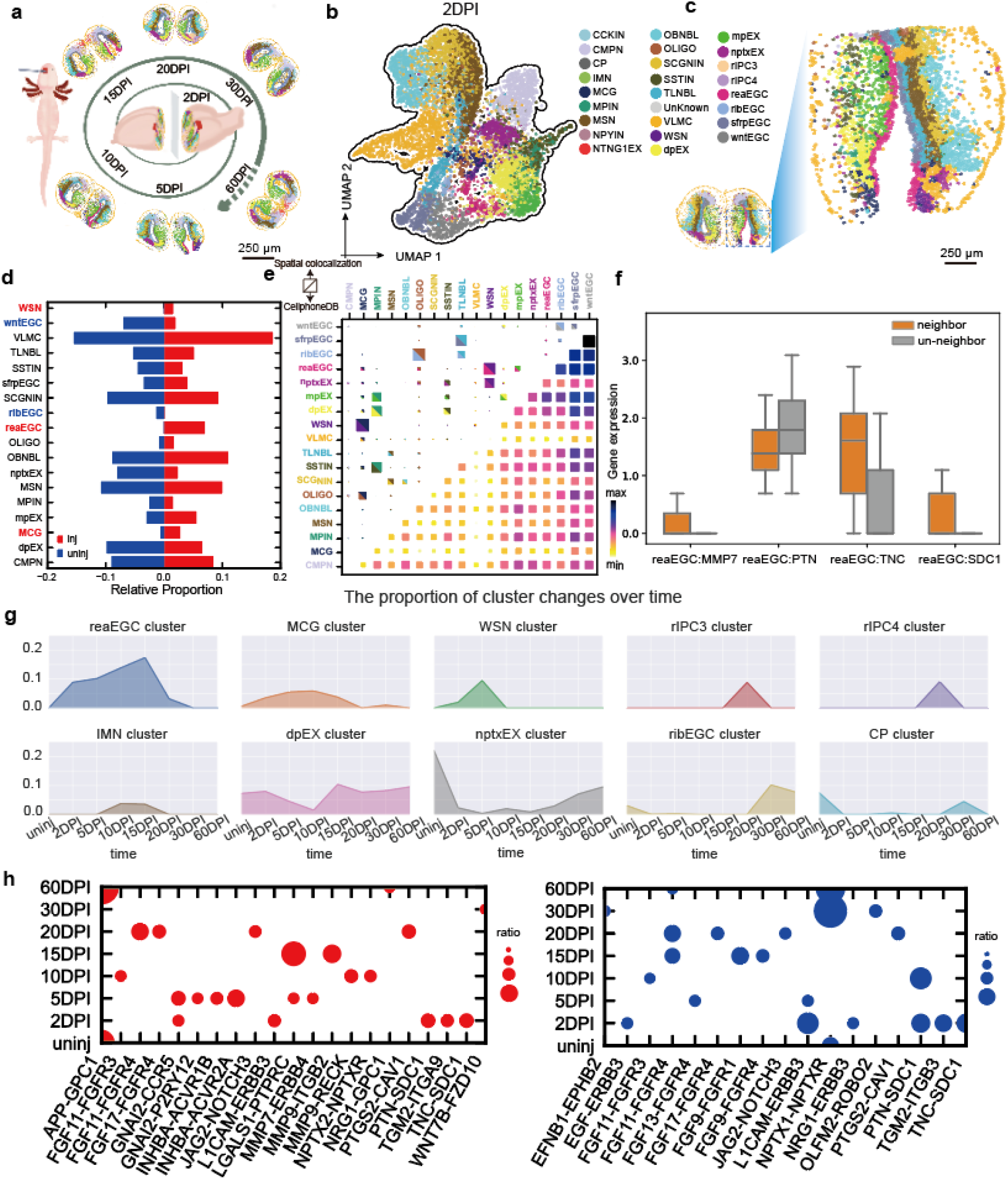
Temporal progression of the post-injury response in the axolotl telencephalon. **a**) Schematic diagram of sample collection and spatial visualization of cell types identified in the axolotl telencephalon sections by Stereo-seq at different stages of regeneration. Cells are colored using the same color legend in panel **b**. **b**) UMAP visualization of the segmented cells from 2 days post-injury (2DPI). **c)** Spatial visualization of the injured area on the 2 DPI section. **d**) Bar graph showing the fold change of the cell ratio in the injured hemisphere compared to the uninjured hemisphere at 2DPI. Significantly increased (WSN, reaEGC, and MCG) or decreased (wntEGC, ribEGC) cell types in the injured hemisphere are annotated with red and blue text, respectively. **e**) The use of spatial proximity reduces the number of false positive interactions and increases specificity on the axolotl brain regeneration dataset. Left and top axes display the identified cell types of axolotl brain at 2-DPI. Top-left triangle: colocalization inferred using actual spatial proximity. Bottom-right triangle: total CCI (cell-cell interaction) number inferred by CellPhoneDB (Efremova et al., 2020). Box sizes correspond to the number of interactions or strength of connectivity for the top-left and bottom-right triangles, respectively. The color scale in the bottom-right triangle has the same meaning as the box size. **f**) Boxplot shows ligand or receptor expression difference in spatial neighboring and not neighboring reaEGCs and WSNs. **g**) Line graphs showing the fold change of the cell ratio across time for the injured hemisphere, up to 60 DPI. **h**) Dotplot shows the interaction between the spatial regions corresponding to the two cell types occurred during the regeneration of nptxEX and dpEX, during the reprogramming process. **Upper:** The most significant interactions throughout the process of **nptxEX** regeneration. At 30 DPI and 60 DPI, the significant ligand-receptor pairs include *NPTX1-NPTXR* appeared also when cells were uninjured, but other times have different interaction signatures, e.g.: 2DPI-10DPI: *PTN-SDC1, L1CAM-ERBB3*, 15DPI-20DPI: *FGF11-FGFR4, JAG2-NOTCH3, PTGS2-CAV1*. **Lower**: Same as above but for **dpEX** regeneration. At 60 DPI, the significant ligand-receptor pairs include *APP-GPC1* appeared also when cells were uninjured, but other times have different interaction signatures, e.g.: 2DPI-5DPI: *GNAI2-P2PY12* and *INHBA-ACVR1A/B*, 15DPI: *LGALS1-PTPRC*, 20DPI: *PTGS1-CAV1*.

**Supplementary figure 3, related to Fig. 3.**
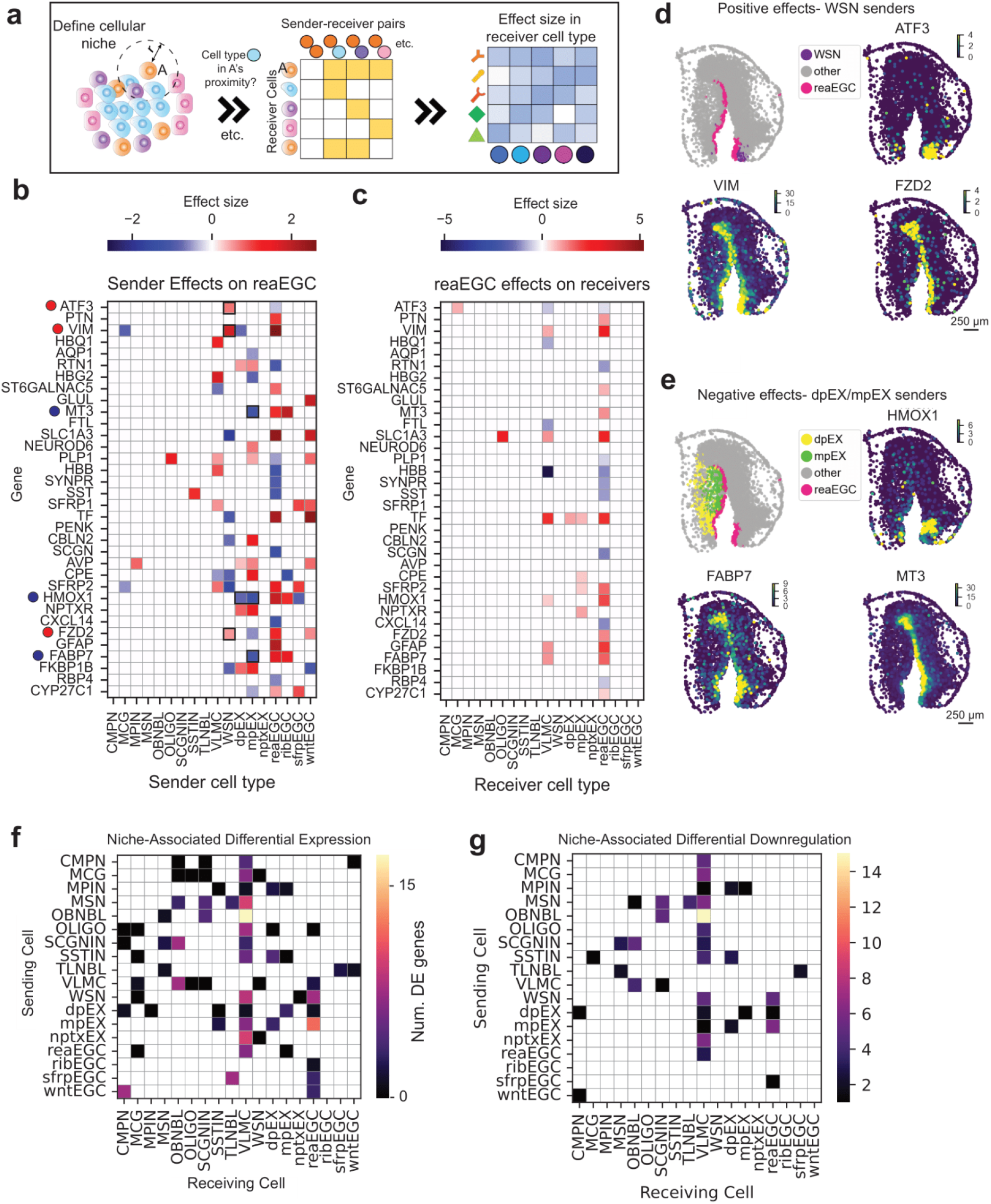
Estimated effect of niche composition on expression in specific receiving cell types. **a**) Construction of the design matrix for Spateo’s niche model, in which the cell type neighborhood of each cell is encoded into a feature array for regression on the corresponding gene expression vector. Parameters from the regression are used to estimate the relative strength of the association between a given cell type pair and gene expression. **b**) Heatmap of sender cell types’ effects on selected genes in reaEGC. For each gene, effect sizes correspond to a subset of coefficients of the spatial niche regression, each coefficient describing the case where reaEGC is the receiver cell type and the given cell type (bottom axis) is present in the local neighborhood of a reaEGC. Genes labeled with red markers are visualized in panel **c**, and genes labeled with blue markers are visualized in panel **d**. **c**) Heatmap of reaEGC’s effects on selected genes in different receiver cell types. For each gene, effect sizes correspond to a subset of coefficients of the spatial niche regression, each coefficient describing the case where a given cell type (bottom axis) is the receiver cell type and reaEGC is present in the local neighborhood of that cell as the sender cell type. **d**) Spatial distribution of reaEGCs and WSNs (**upper left**) and expression patterns for *ATF3, VIM* and *FZD2*, validating enrichment along the WSN-reaEGC interface corresponding to the estimated positive effect size. **e**) Spatial distribution of dpEX, mpEX and reaEGCs (**upper left**) and expression patterns for *HMOX1, FABP7* and *MT3*, validating relative depletion along the reaEGC-mpEX/dpEX interface corresponding to the estimated negative effect size. **f**) From outputs of the model labeled **I** in the right box of panel **a,**number of genes that are differentially expressed (either positively or negatively) in association with proximity of a given pair of cell types. **g**) From outputs of the model labeled **I** in the right box of panel **a,**number of genes that are both differentially expressed and negatively associated with proximity of a given pair of cell types (labeled as “differentially downregulated”).

**Supplementary figure 4, related to Fig. 3.**
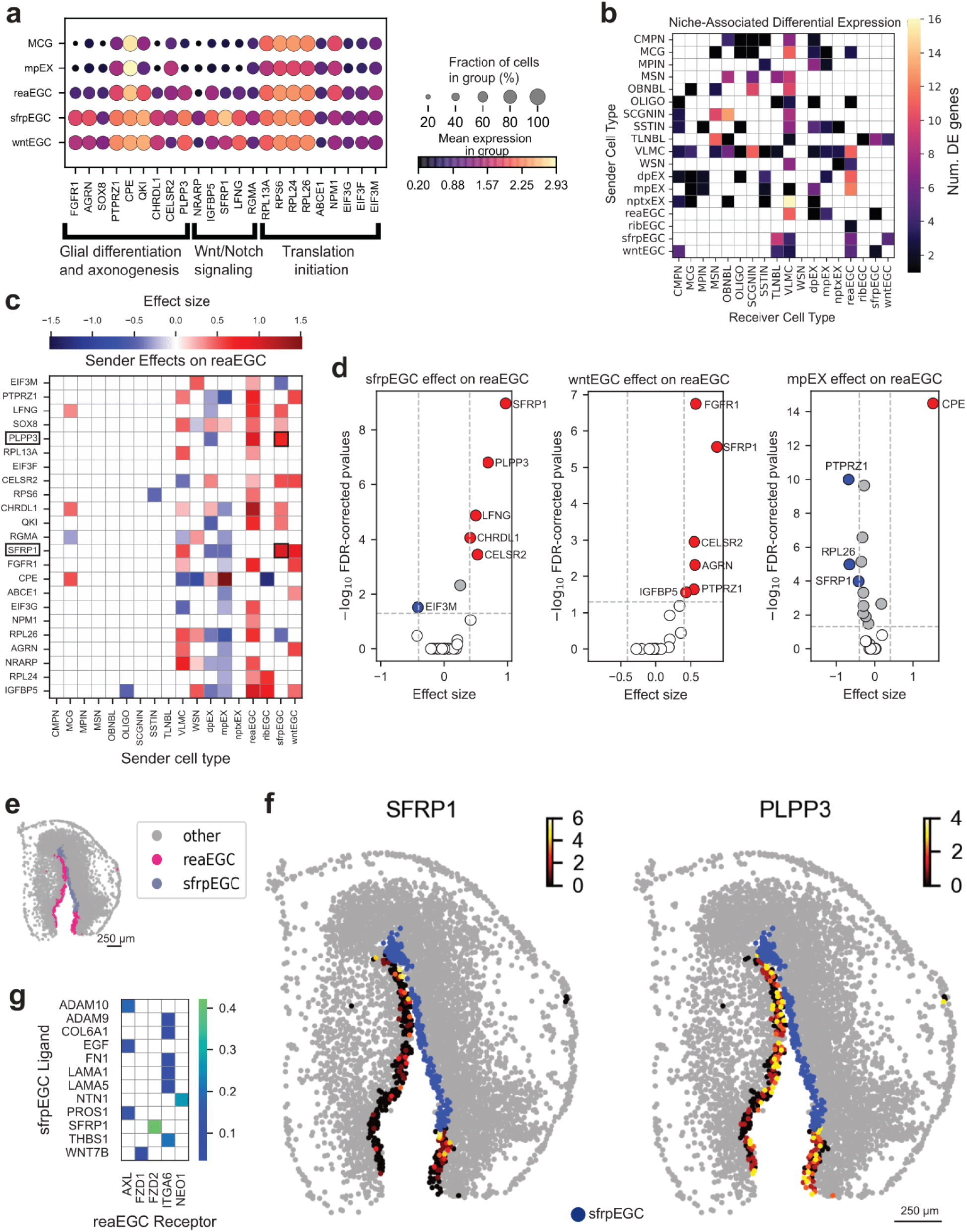
Characterization of niche effects on neuronal regeneration signatures. **a**) Expression of gene sets associated with glial differentiation and axonogenesis, Wnt/Notch signaling pathways and ribosomal translation in reaEGC, sfrpEGC, wntEGC, microglia (MCG) and mpEX, colored by mean expression and with dot sizes corresponding to the proportion of cells with nonzero expression. **b**) Number of differentially expressed genes (assessed by significance testing of coefficients) associated with proximity of a given pair of cell types, with neighboring (sender) cell types along the left axis and receiver (refer to annotation “A” in the “Microenvironment Modeling” panel in **Fig. 3a**) cells along the bottom axis. **c**) Heatmap of sender cell types’ effects on selected genes in reaEGC. For each gene, effect sizes correspond to a subset of coefficients of the spatial niche regression, each coefficient describing the case where reaEGC is the receiver cell type and the given cell type (bottom axis) is present in the local neighborhood of a reaEGC. Elements of interest for panel **f** are highlighted. **d**) From left to right, the estimated effect, with respect to both coefficient and -log_10_(*q*-values), of sfrpEGC, wntEGC and mpEX on expression of each gene in reaEGC. The patterns of differential expression are slightly different across sender cell types, with other radial glial cells having a greater number of positive associations compared to mature excitatory neurons. **e**) Spatial distribution of reaEGCs and sfrpEGC. **f**) reaEGC-specific expression pattern for *SFRP1* (**left**) and *PLPP3* (**right**) in conjunction with spatial distribution of sfrpEGCs, showing the tendency for sfrpEGCs to colocalize with expression hotspots in reaEGCs for two genes sfrpEGC is predicted to have positive effect on. **g**) Subset of identified nonzero interactions resulting from paired testing of sfrpEGC-produced ligands and reaEGC receptors as described in the **left box** of **Fig. 3a**, representing potential mechanisms that may result in the observed differential expression patterns.

**Supplementary figure 5, related to Fig. 3.**
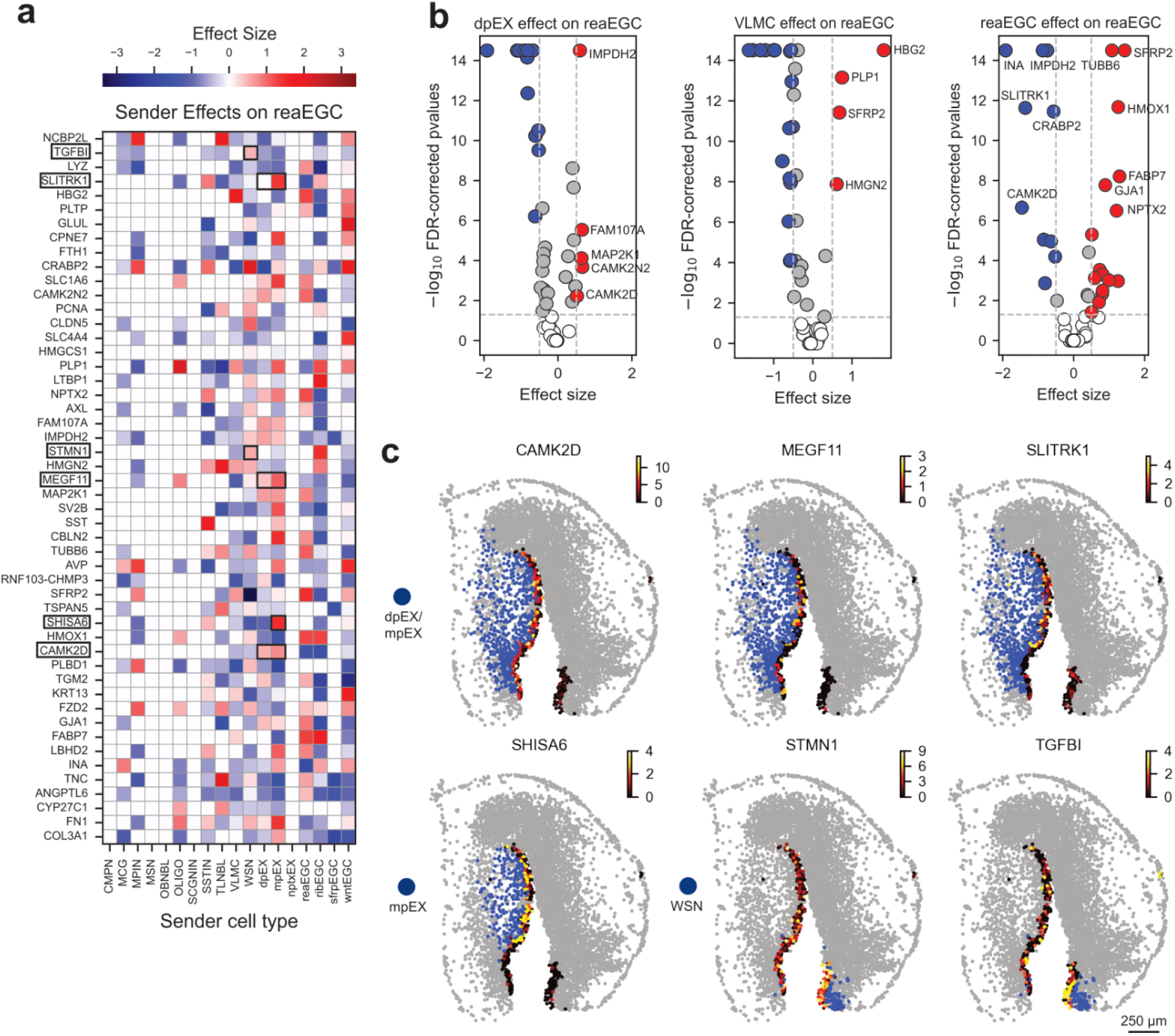
Characterization of niche effects on reaEGCs by regressing on many genes and selecting subset of interest. **a**) Heatmap of sender cell types’ effects on selected genes in reaEGC. For each gene, effect sizes correspond to a subset of coefficients of the spatial niche regression, each coefficient describing the case where reaEGC is the receiver cell type and the given cell type (bottom axis) is present in the local neighborhood of a reaEGC. **b**) From left to right, the estimated effect, with respect to both coefficient and -log_10_(*q*-values), of dpEX, VLMC and reaEGC (assessing homologous cell pairs) on expression of each gene in reaEGC. The patterns of differential expression are slightly different across sender cell types. **c**) reaEGC-specific expression pattern for *CAMK2D, MEGF11, SLITRK1, SHISA6, STMN1* and *TGFB1* in conjunction with spatial distribution of dpEX and mpEX (*CAMK2D, MEGF11, SLITRK1*), mpEX alone (*SHISA6, STMN1*) and WSN (*TGFB1*).

**Supplementary figure 6, related to Fig. 3.**
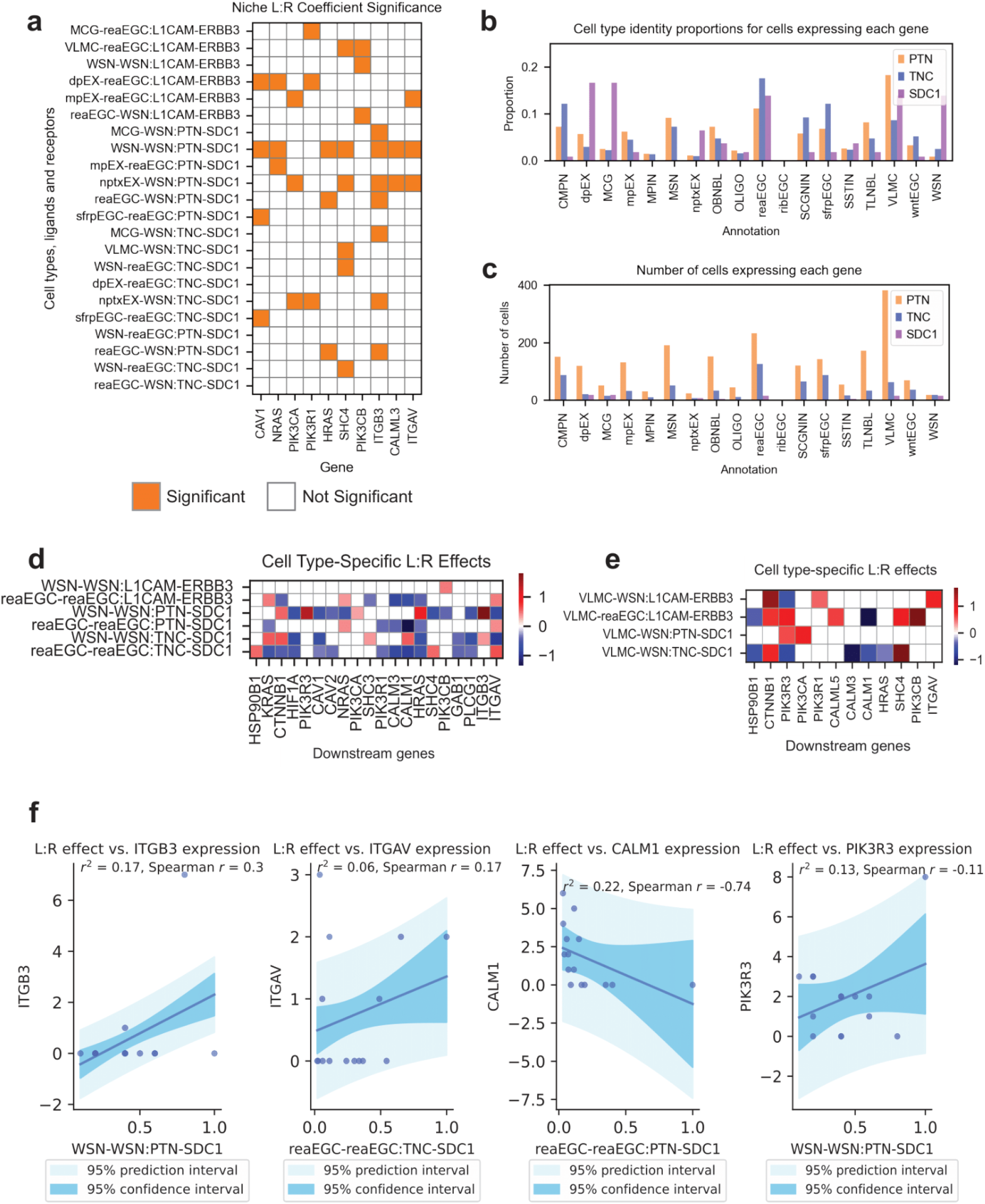
Characterizing the effect of niche-specific, cell type-specific ligand-receptor interactions on expression in specific receiving cell types. **a**) Significance of coefficients for the regression of each combination of cell type pairs, ligands and receptors on downstream targets, subsetted to features containing reaEGC or WSN. Elements are colored if significant, and uncolored if not. **b**) The proportion of the total number of cells with nonzero expression of *PTN, TNC* and *SDC1* for each labeled cell type. **c**) Barplot of the number of cells labeled for each specific cell type with nonzero expression of *PTN, TNC* and *SDC1*.**d**) Estimated effect sizes when cell type-specific patterns (specifically, reaEGC-reaEGC and WSN-WSN specific patterns) of ligand-receptor interaction are used to predict the expression patterns of select genes. Genes chosen by using a prior knowledge network database(Gillespie et al., 2022; Hu et al., 2019; Kanehisa and Goto, 2000; Shao et al.,2021) to choose genes downstream of *ERBB3* and *SDC1* (see **SLICE model** in **STAR Methods**). **e**) Same as in panel **d**, but for VLMC-reaEGC and VLMC-WSN specific patterns. **f**) Coefficient of determination and Spearman correlation for linear fits of cell type-specific ligand-receptor products and gene expression for select genes (*ITGB3, ITGAV, CALM1, PIK3R3* from left to right). Cell type-specific ligand-receptor products are computed by subsetting to receiver cells that both neighbor the indicated sender cell and express the indicated receptor, resulting in between 8-15 receiver cells. The cell type-specific ligand-receptor product is min-max scaled. 95% prediction intervals and 95% confidence intervals are superimposed in shades of blue, with 95% confidence intervals corresponding to the darker shade.

**Supplementary figure 7, related to Fig. 4.**
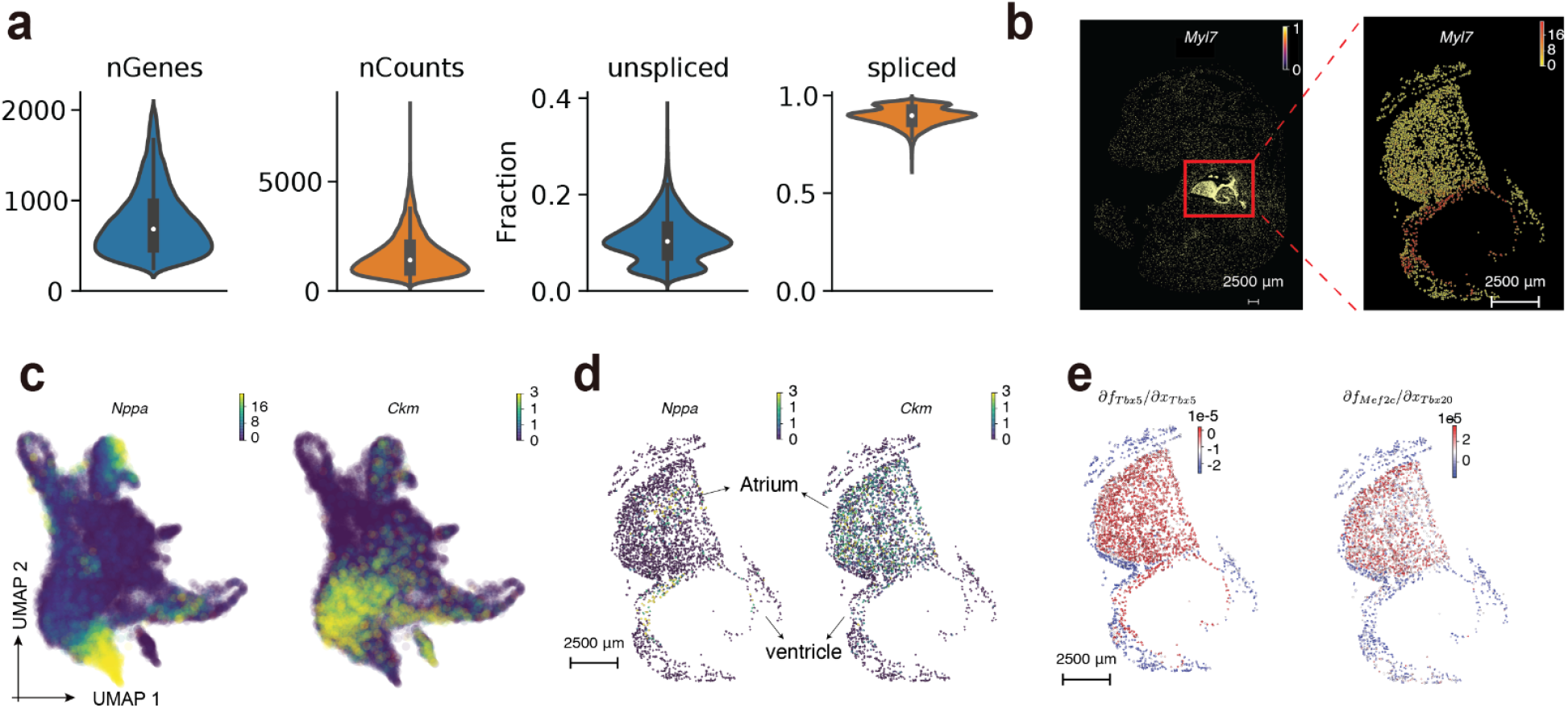
RNA velocity analyses of spatially resolved heart cell atlas during mouse organogenesis. **a**). Basic statistics (number of genes, *nGenes*, and number of total UMI counts, *nCounts*), and the fraction of unspliced and spliced RNAs of cardiac cells of the heart cell atlas. **b**). Spatial gene expression distribution of cardiac cell marker, *Myl7*, across the entire embryo and extracted heart region at E16.5. **c**). Gene expression of regional specific cardiomyocyte markers, *Nppa* and *Ckm* across all cardiac cells in the UMAP space. **d**). Spatial gene expression of *Nppa* and *Ckm* on the heart region of the typical E16.5 Stereo-seq slice (also shown in panel **Fig. 4a**), same as for the spatial plots in the panel **e**. **e**) Spatial RNA Jacobian of the regulation from *Tbx5* to itself or the regulation from *Tbx20* to *Mef2c*.

**Supplementary Figure 8, related to Figure 5:**
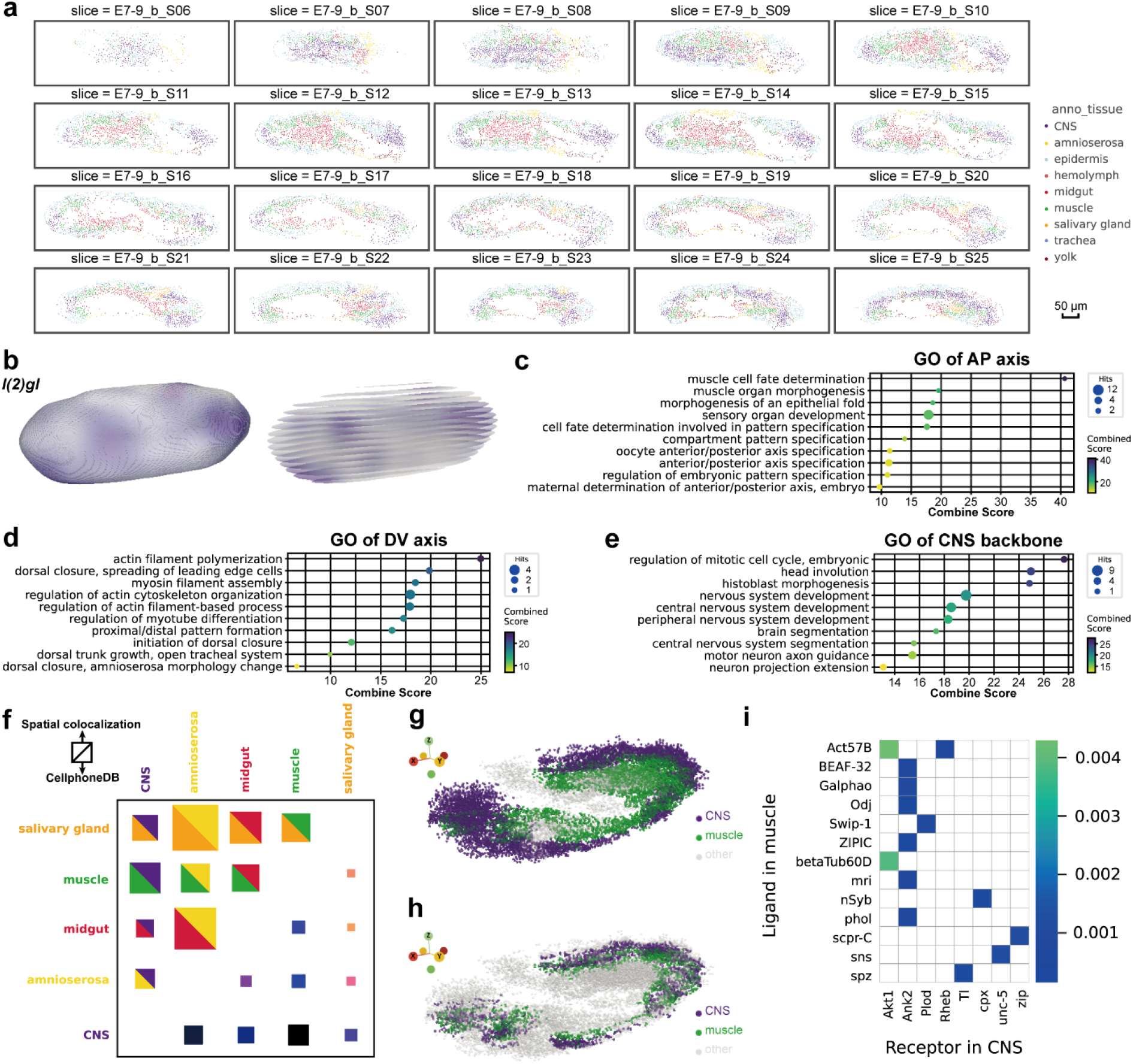
3D tissue/embryo reconstruction enables discovering spatial axis and tissue backbone dependent genes, and quantifying tissue-level cell-cell interactions. **a**). Scatterplots of aligned cells on each Stereo-seq slice. The color corresponds to the annotated tissue type. **b**). 3D surface (**left**) and computational slicing (**right**) visualization of the smoothed expression of A-P axis-dependent gene *ab* imputed with a neural net. **c-e**). Enriched GO biological pathway terms of significant A-P, D-V and CNS backbone dependent genes. **f**). Tissue interaction plot based on spatial proximity (**top-left triangle**) and inference by CellPhoneDB (**bottom-right triangle**). Box size corresponds to the strength of cell-cell interactions based on spatial colocalization or the number of CellPhoneDB predictions. **g**). 3D spatial distribution of cells from CNS and muscle cells. **h**). Same as in **g**, but pruned for CNS and muscle cells that are neighboring one another. **i**). Identified significant ligand and receptor pairs between sending muscle cells and receiver CNS cells.

**Supplementary Figure 9, related to Figure 6:**
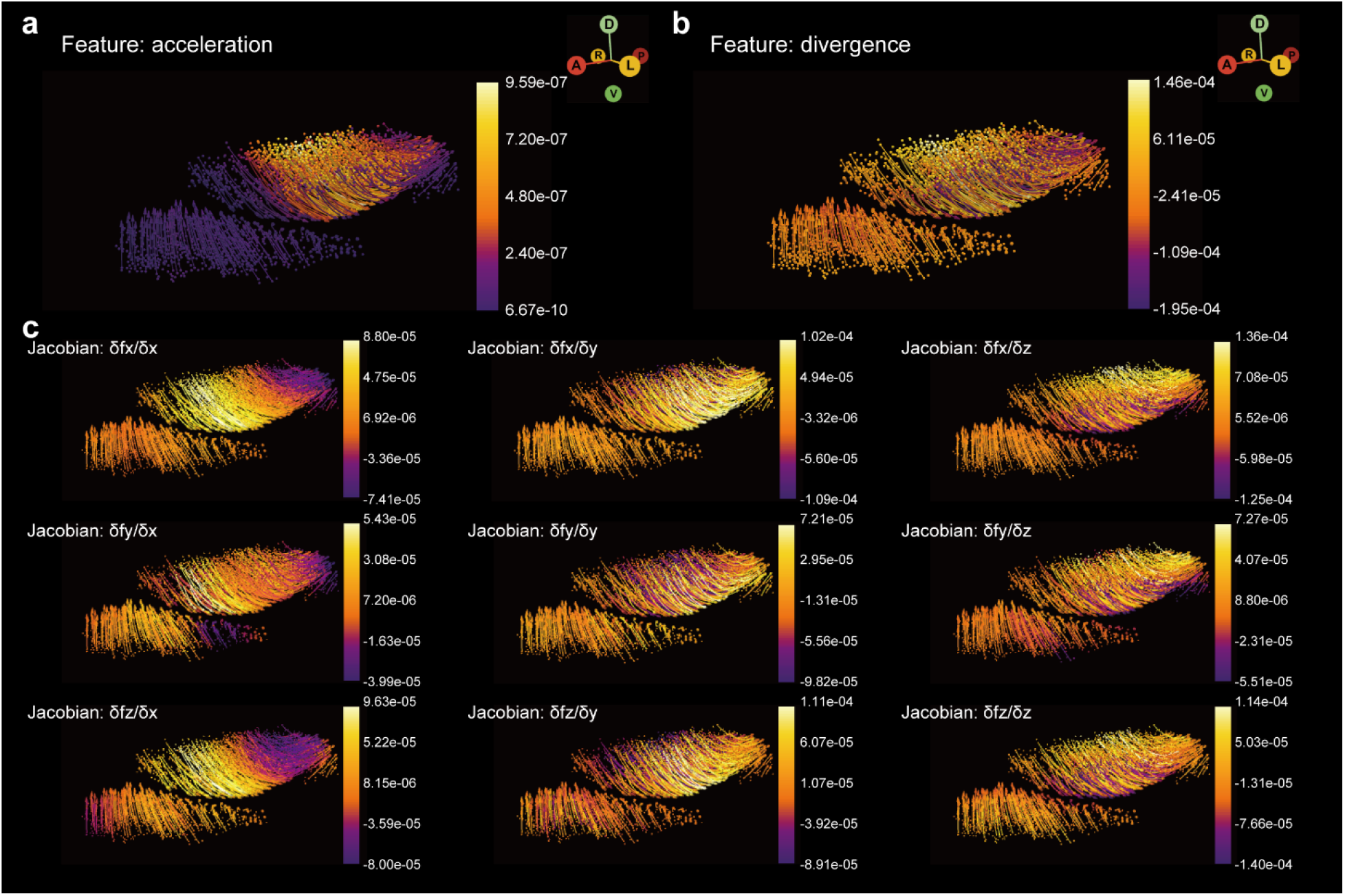
Spatial acceleration (a), divergence (b), and Jacobian (c) of the morphometric vector field of Drosophila midgut morphogenesis.

**Supplementary Figure 10, related to Figure 6:**
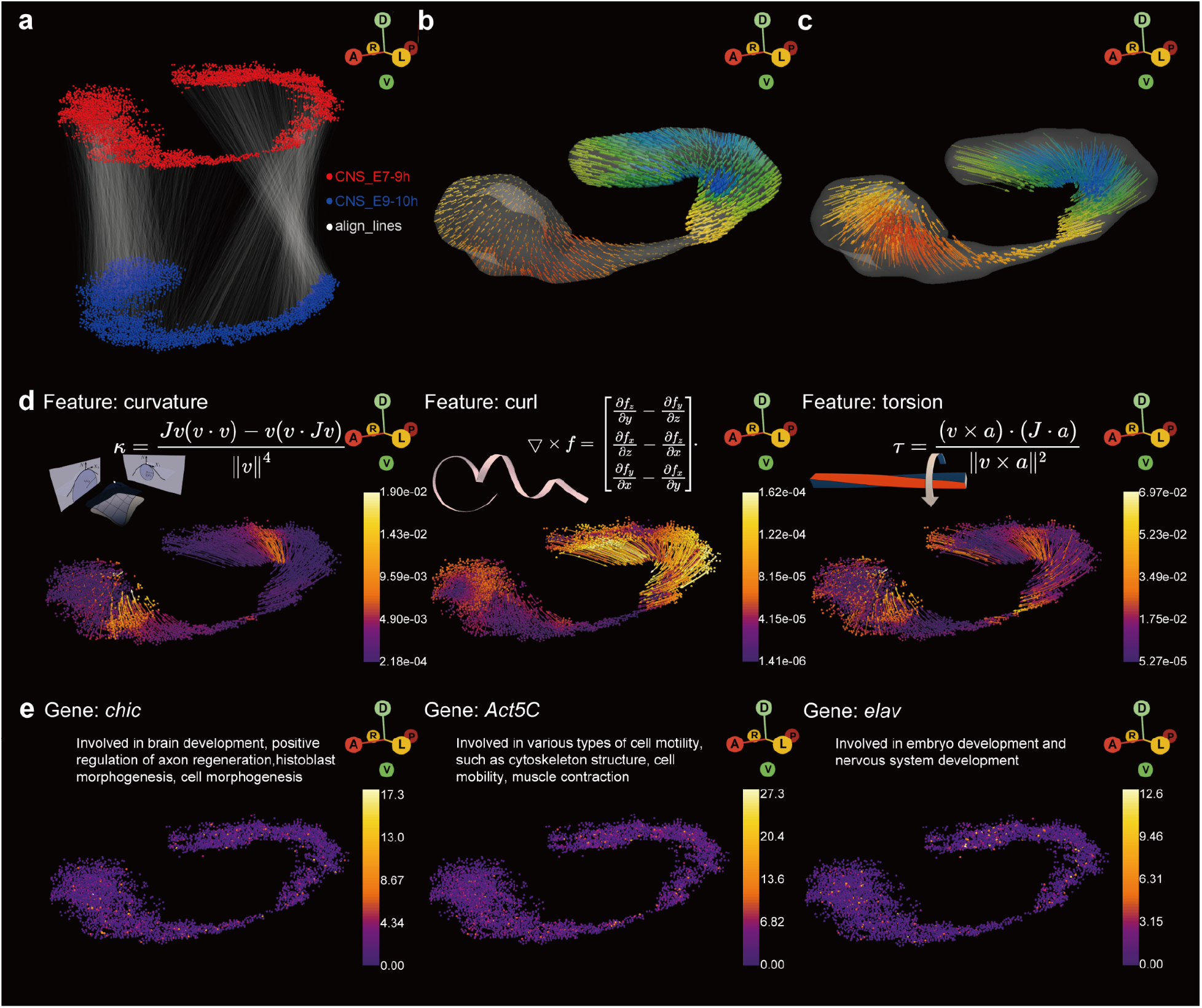
Morphometric vector field analysis of Drosophila central nervous system (CNS) morphogenesis. **a**). The alignment between cells of *Drosophila* CNS from E7-9h to those from E9-10h. **b**). Quivers of the surface of CNS predicted by the morphometric vector field learned with the cell alignment defined in panel **a**). Quivers are sampled uniformly across the surface. Color of the quiver corresponds to the z component of the velocity vector. **c**). Same as the **b**, but with the predicted trajectories of sample points, visualized as streamlines. **d**). Estimated morphogenic curvature, curl, torsion of the reconstructed morphometric vector field. The cell and streamline are colored by the corresponding norm of each quantity. Definition of each quantity is provided. Note that both curvature and curl are vectors of the *x,y,x* axis, while torsion is a 3 x 3 matrix for each sample point. **e**). Top example gene that is significantly associated with each differential geometry quantity. Short description of the function of each gene that is related to CNS development, or *Drosophila* morphogenesis in general is provided.

**Supplementary Figure 11, related to Figure 6:**
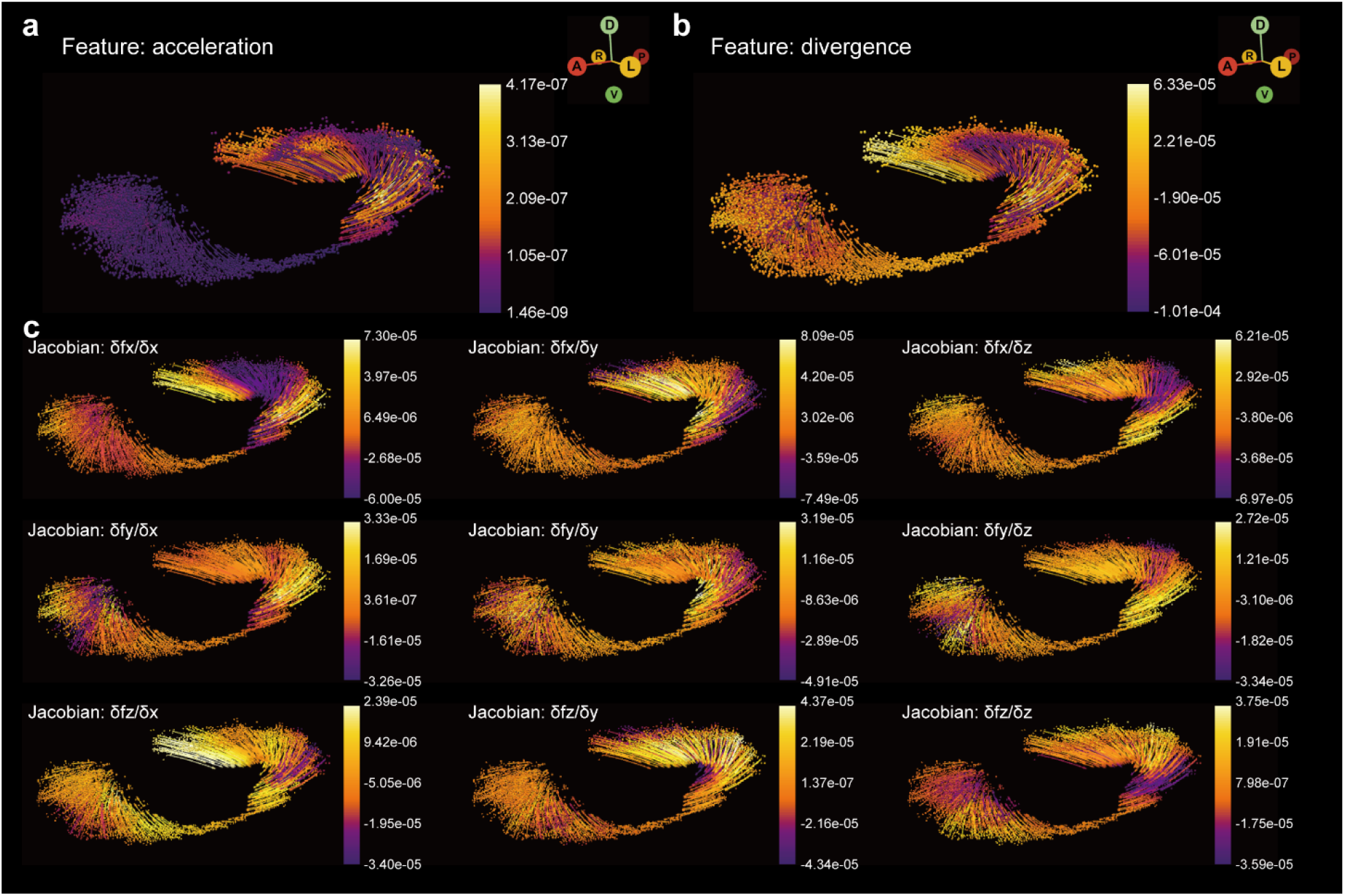
Spatial acceleration (a), divergence (b), and Jacobian (c) of the morphometric vector field of Drosophila central nervous system (CNS) morphogenesis.

**Supplementary Figure 12:**
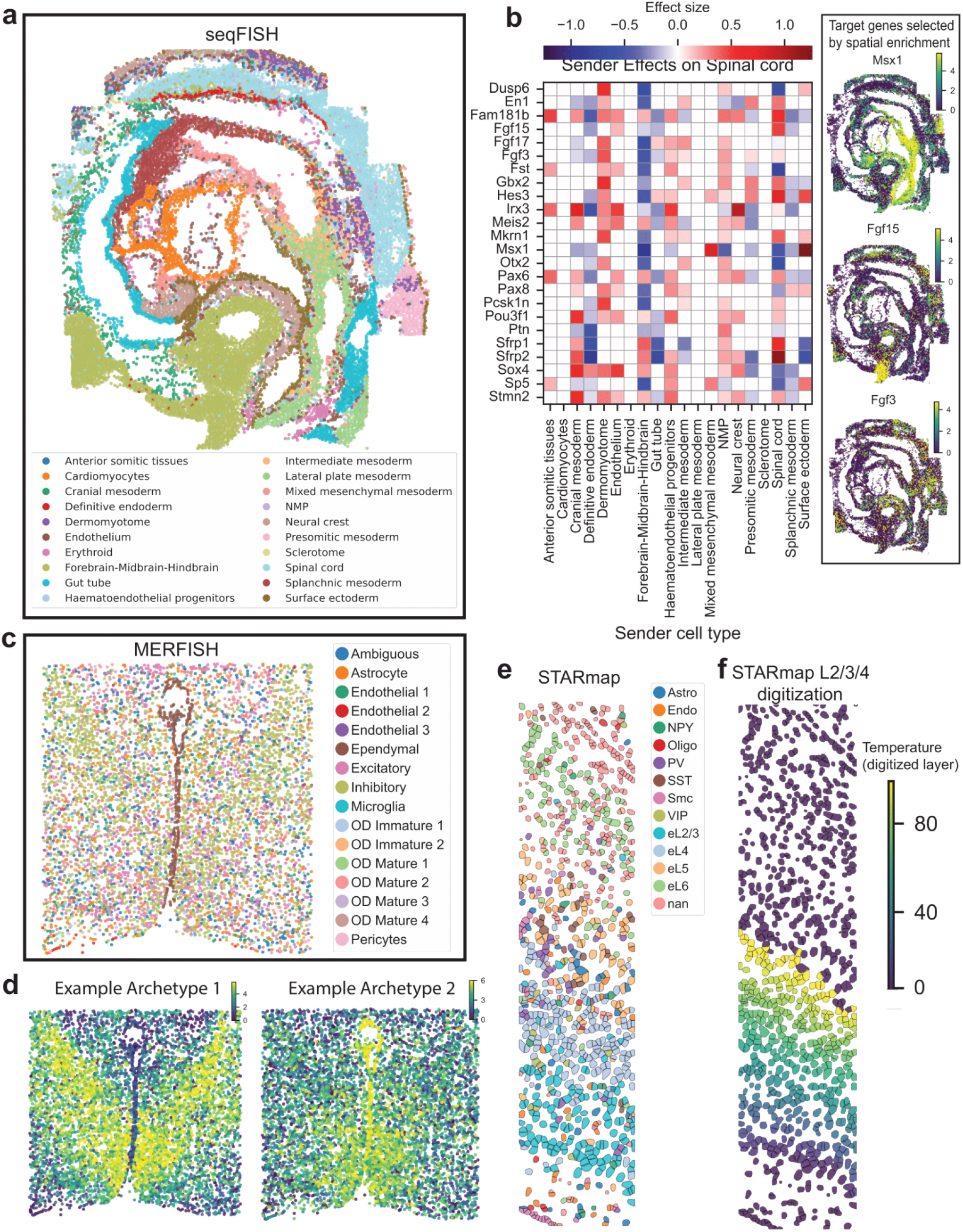
Spateo facilitates analysis of data collected with *in situ*-hybridization based methods. For *in situ* hybridization methods, cells can be identified from spot-based readouts using Spateo’s cell segmentation method, or other methods such as Ilastik(Berg et al., 2019) (as was used for the seqFISH and STARmap datasets here). **a**) seqFISH dataset collected from a mouse embryo in the somite stage. Cells are colored by tissue type, with progenitor cells that will become certain tissue types also included. **b**) Heatmap of sender cell types’ effects on selected genes in the developing spinal cord. For each gene, effect sizes correspond to a subset of coefficients of a spatial niche regression, each coefficient describing the case where spinal cord cells are the receiver cell and the given cell type (bottom axis) is present in the local neighborhood of a spinal cord cell. Gene set contains all genes with Moran’s *I* coefficient that is significant (with adjusted *p-*value threshold set to 0.05) and >0.1, with expression patterns for example genes *Msx1, Fgf15* and *Fgf3* in the right subpanel. **c**) MERFISH dataset collected from the posterior preoptic region of the mouse hypothalamus. Cells are colored by neuronal cell type. **d**) Expression of select gene expression archetypes (representing hierarchical clusterings of genes) in space. **e**) STARmap dataset collected from the mouse medial prefrontal cortex. Cells are colored by neuronal cell type. **f**) Digitization of cortical layers (L2-L4) by solving the layer-wise potential equation. Cells are colored by the value returned from this solution (electric potential from the equation), representative of their digitized layer number.

**Supplementary Figure 13:**
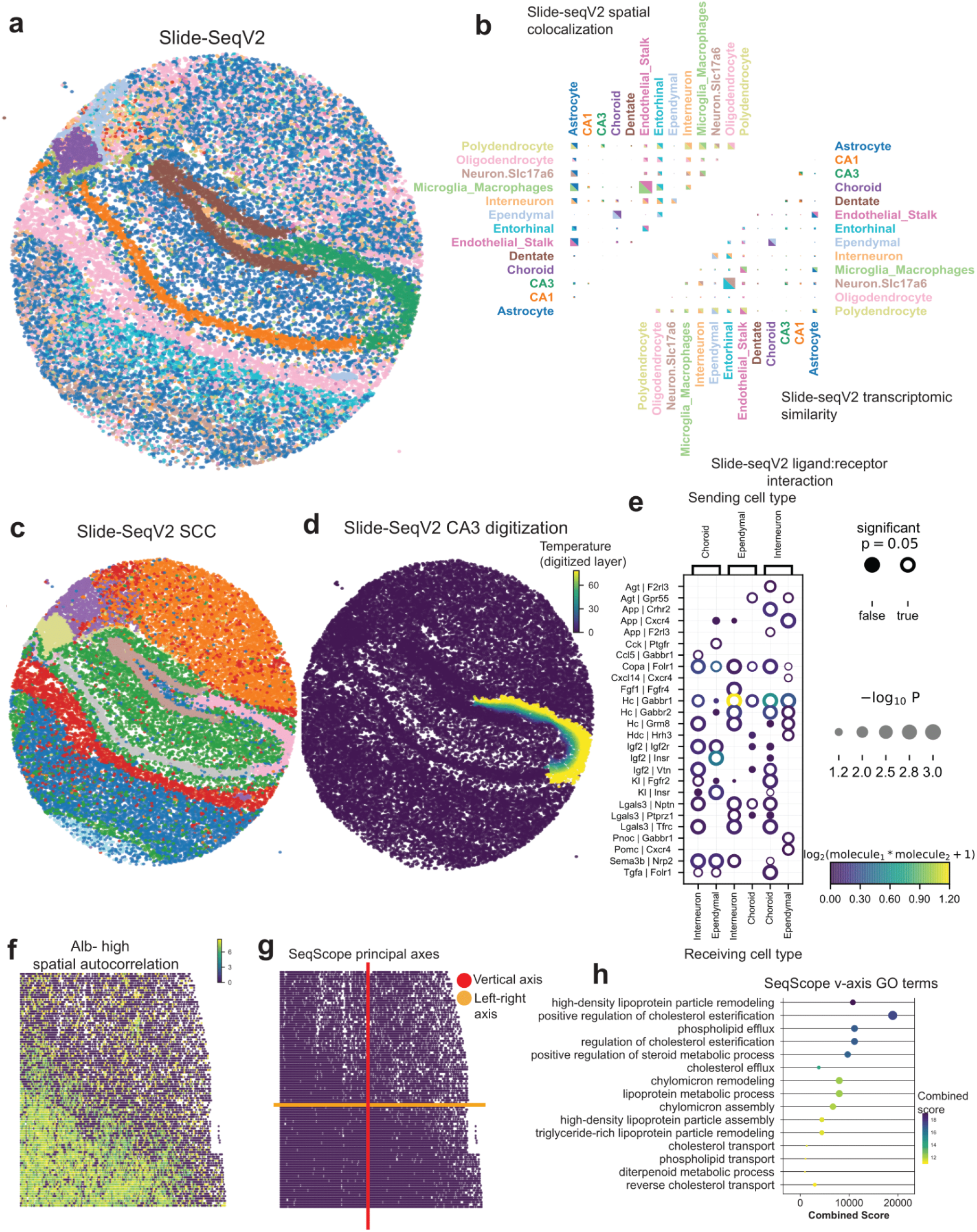
Spateo facilitates analysis of data collected with *ex situ*-sequencing based methods. **a**) Slide-seqV2 dataset collected from the mouse hippocampus and surrounding section of brain. Cells colored by the primary cell type inferred by RCTD(Cable et al., 2022a). Refer to panel **b**) for cell type labels and corresponding colors. **b**) Quantification of spatial colocalization between different cell types. *Top triangle:* spatial connectivity; *bottom triangle:* expression connectivity. Spatial connectivity is computed based on weighted spatial distances and cell type labels and normalized by number of cells, and expression connectivity is computed based on weighted gene expression space distances. Box size corresponds to the strength of connectivity. Some cell types are pre-filtered on the basis of low total cell count. **c**) Spatial domains identified by the unsupervised SCC method at bin 10 resolution. **d**) Digitization of the CA3 region of the hippocampus by solving the layer-wise potential equation. Cells are colored by the value returned from this solution (electric potential from the equation), representative of their digitized layer numbers. **e**) Most significant ligand-receptor interactions between different sending and receiving cell types in the Slide-seqV2 data. All interactions involve combinations of interneurons, ependymal cells and choroid plexus cells. Ligand-receptor pair on left axis, sender and receiver cell types on bottom and top axes, respectively, corrected *p*-value denoted by circle size, ligand-receptor product by circle color, and significance by an open or filled circle. **f**) From the Seq-Scope dataset collected from the mouse liver, the gene expression distribution of *Alb* over space, representing an example of a gene with high spatial autocorrelation, determined by Moran’s *I* score >0.1 and *q*-value < 0.05. **g**) Learned principal axes of the 2D Seq-Scope dataset. **h**) Enriched GO biological pathway terms of significant vertical axis-dependent genes, determined using a linear regression. Combined score computed by the product of the log-transformed *p*-value and the z-score for the GO term.

## Supplementary Tables and Animations

**Supplementary table 1**: The table that compares Spateo with existing major spatial transcriptomics dataset analyses toolkits, including Squidpy, stLearn, Giotto, Seurat and StUtility in terms of package focus, infrastructure, spatial statistics, image analysis, visualization, integration, advanced spatial and temporal analysis and others.

**Supplementary table 2**: The table of the results of A-P (anterior-posterior) / D-V (dorsal-ventral) axis dependent genes, associated GO (gene ontology) enrichment and 3D cell-cell interactions analyses. There are seven sheets in total, namely: 1). Results of A-P axis dependent differential expression analysis; 2). Results of GO enrichment analysis of the A-P axis dependent genes; 3). Same as 1) but for D-V axis; 4). Same as 2) but D-V axis dependent genes; 5). Same as 1) but for CNS (central nervous systems) backbone (see **STAR Methhods**); 6). Same as 2) but for CNS backbone dependent genes; 7). Top identified ligand-receptor pairs between muscle and CNS cells.

**Supplementary table 3**: The table of the results of morphometric vector field differential geometry analyses (see **STAR Methods**). There are six sheets in total, namely: 1). Significant curvature dependent genes in Drosophila midgut; 2). Significant curl associated genes in Drosophila midgut; 3). Significant torsion associated genes in Drosophila midgut; 4). Significant curvature associated genes in Drosophila CNS; 5). Significant curl dependent genes in Drosophila CNS; 6); Significant torsion dependent genes in Drosophila CNS.

**Supplementary animation 1**: Animation showing the reconstructed 3D models of embryo and various organs, including CNS, amnioserosa, embryo shell, epidemis, midgut, muscle and salivary gland in 3D space, rotating firstly along the D-V axis and then the A-P axis.

**Supplementary animation 2**: Animation of the the predicted cell migration paths of E7-9h Drosophila midgut cells. Animation is repeated with cells colored with the norm of acceleration, divergence, curvature, curl, torsion sequentially at each repeat.

**Supplementary animation 3**: Same as in supplementary animation 2 but for the CNS cells.

## STAR+METHODS

### KEY RESOURCES TABLE

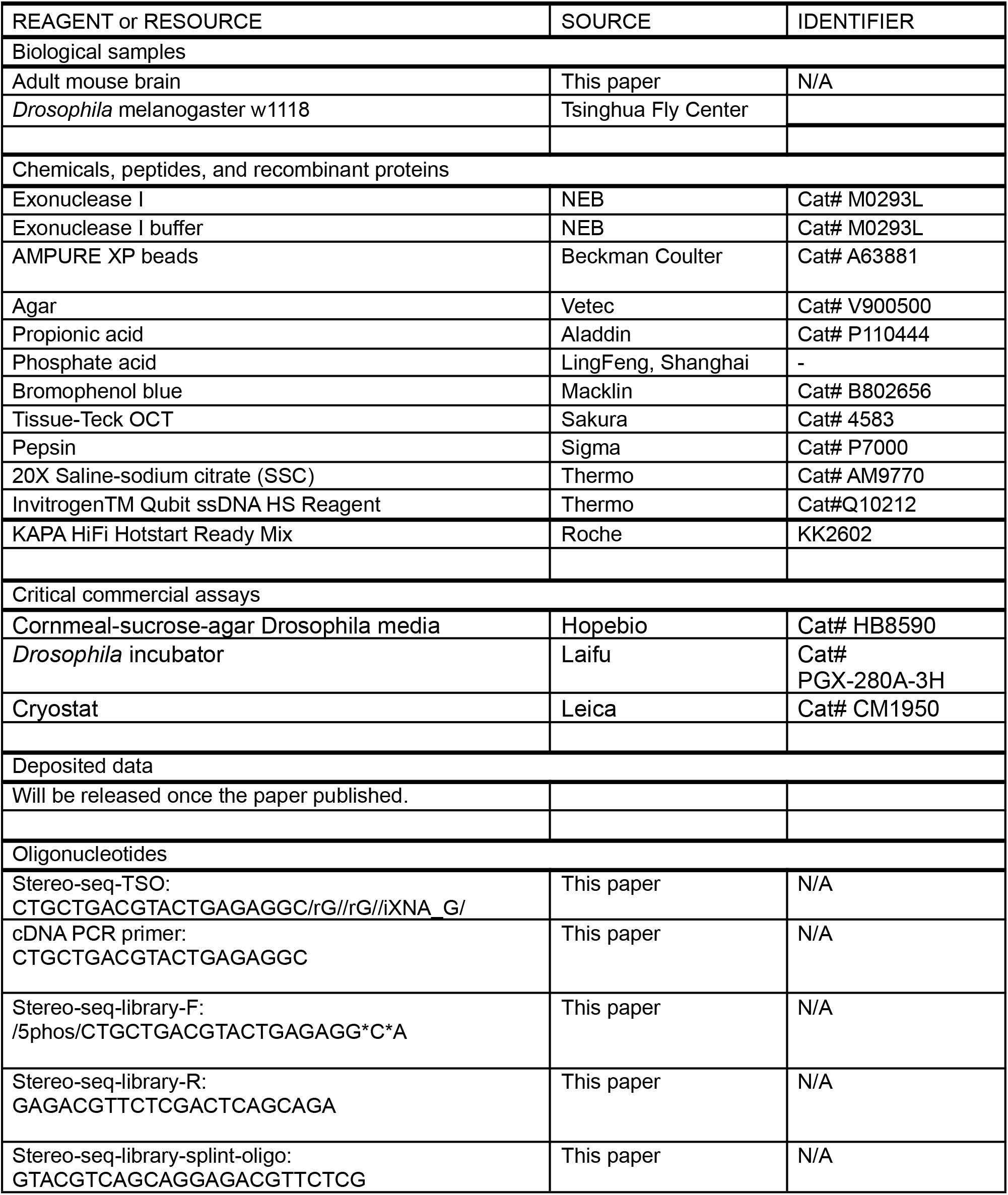
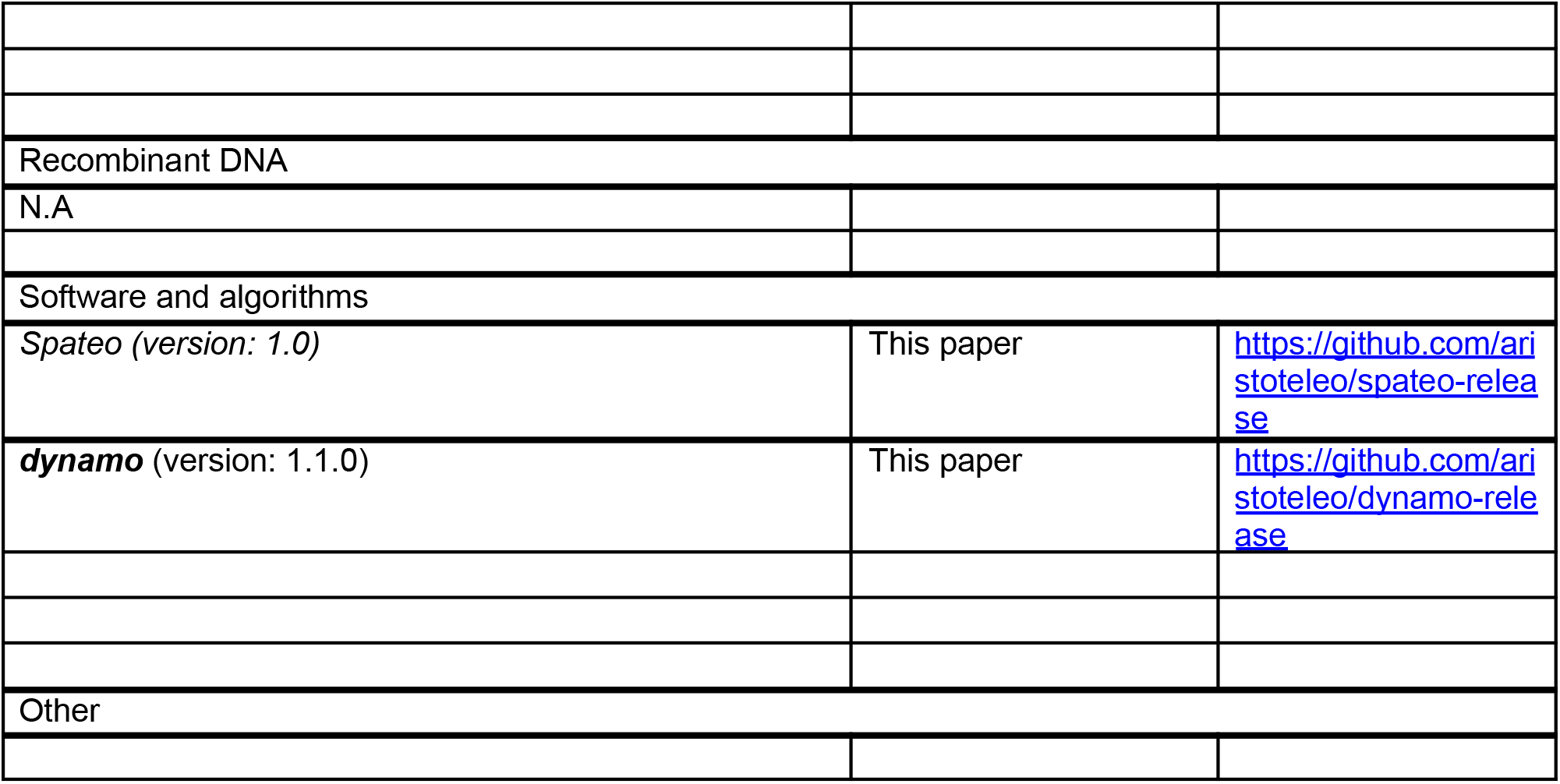

### RESOURCE AVAILABILITY

#### Lead contact

Further information and requests for resources and reagents should be directed to and will be fulfilled by the Lead Contact Xiaojie Qiu.

#### Materials availability

This study did not generate new unique reagents.

#### Data and code availability

The following public Stereo-seq datasets are used in this study: the axolotl brain regeneration dataset (Wei et al., 2022), MOSTA datasets (Chen et al., 2022), mouse hippocampus Slide-seq V2 dataset (Stickels et al., 2021) (https://singlecell.broadinstitute.org/single_cell/study/SCP815), mouse embryo seqFISH dataset (Lohoff et al., 2022) (https://crukci.shinyapps.io/SpatialMouseAtlas/), mouse hypothalamus dataset (Moffitt et al.,2018) (https://datadryad.org/stash/dataset/doi:10.5061/dryad.8t8s248), mouse cortex STARmap dataset (Wang et al., 2018) and mouse liver seqScope dataset (Xi et al., 2022)). E14-16h and E16-18h Drosophila 3D Stereo-seq datasets are downloaded from the Flysta3D database https://db.cngb.org/stomics/flysta3d/download.html(Wang et al., 2022).

Mouse adult coronal hemibrain and continuous slicing profiling of Drosophila embryogenesis datasets are newly generated for this study. Processed datasets are provided within Spateo and the raw data will be released upon the publication of this study.

***Spateo*** (version: 1.0) is implemented as a Python package and is available through GitHub (https://github.com/aristoteleo/spateo-release). Notebooks, tutorials for reproducing all figures in this study, and tutorials of ***Spateo*** usage cases are also available through GitHub (https://github.com/aristoteleo/Spateo-notebooks, https://github.com/aristoteleo/Spateo-tutorials).

### METHOD DETAILS

#### Spatially-constrained clustering (SCC)

While conventional clustering analyses of transcriptomic data identify cell type identities and reveal the primary sources of cellular heterogeneity, they are unable to identify regions of cells within a tissue that share transcriptomic properties and correspond to anatomical domains. Spatially-constrained clustering (SCC) jointly models the transcriptomic and spatial information by firstly constructing two k-nearest neighbor graphs, the expression graph 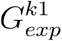 based on distance in transcriptomic space and the spatial graph 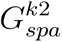 based on distance in the physical space. In the example in **Figure 2**, to construct, 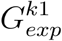, the Stereo-seq data is gridded into 50 * 50 DNB or DNA Nanoballs square lattices (the grids formed by the DNA Nanoballs on the Stereo-seq chip), termed bin 50, or lower resolutions to merge multiple cells (or part of cells) into distinct spatial units. Assuming that physically adjacent cells tend to share similar expression profiles(Nitzan et al., 2019), this binning strategy smooths gene expression which is directly beneficial for identifying spatially continuous tissue domains. The binned data is then normalized for sequencing depth, followed by log-transformation for variance stabilization, ensuring the genes with high expression after log-normalization weigh more in downstream PCA performance and eventually in the identification of spatial domains (Thorrez et al., 2011). By default, the top 30 PCs and 30-nearest neighbors (*k*_1_ = 30) are selected to build the expression graph 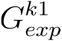, although these parameters can be modified in the function call to *st.tl.pca_spateo()* and *st.tl.neighbors()*. For the spatial graph 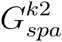, the number of physical neighbors *k*_2_ is set to be 8 by default. Both *k*_1_ and *k*_2_ can be adjusted to balance the weight given to transcriptomic and physical neighbors, and we have demonstrated the robustness of SCC overall performance under a broad range of values for both alongside a series of the coupled parameters (**Fig. S1b**). We take the union of both graphs and obtain 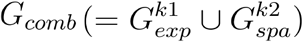, which is then used as input for downstream Louvain or Leiden clustering in the *st.tl.scc()* function. In the end, SCC reveals well separated tissue domains that have distinct intra-tissue transcriptional profiles, which can be further used for downstream analysis, such as spatial column or layer digitization. SCC can be implemented by calling the following:

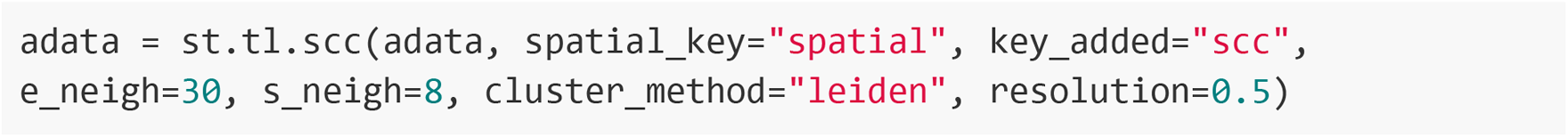

where *e_neigh* and *s_neigh* specify the number of nearest neighbors to consider in gene expression space (number of neighbors in the PCA space) and the number of nearest neighbors in physical space, respectively, and the resolution is the Louvain or Leiden method’s internal resolution parameter, controlling the number of clusters in the final result. For further information of each argument, please refer to the documentation in the Spateo package: https://spateo-release.readthedocs.io/en/latest/autoapi/spateo/index.html, same as for all the following code blocks.

#### Identify gene expression archetypes and archetype associated genes

We adapted the implementation from Novosparc (Nitzan et al., 2019) to identify gene expression archetypes and archetype associated genes. Size factor normalized and log(1+x) transformed gene expression data are used. Pearson correlation matrix of gene expression between detected highly variable genes or all genes passed basic filterings obtained via preprocessing module in Spateo are then calculated. Hierarchical clustering based on the Pearson correlation matrix are then calculated and the resultant cluster number can be determined by the users. Once gene cluster number is determined, we calculate the average gene expression of all the genes belonging to each cluster and define the average gene expression pattern as the gene expression archetypes. To identify archetype associated genes, we calculate the Pearson correlation coefficient between each gene within the corresponding cluster to the archetype expression pattern of the cluster of interests. Genes with highest correlations are regarded as the archetype associated genes.

#### Spatial domain digitalization

Spatial domain digitalization describes the process of constructing a spatial coordinate reference system in accordance with any arbitrary axis, enabling identification of genes with graded or periodic distributions along the directions defined by this coordinate system. Mathematically, for an arbitrarily-shaped spatial domain, a digitalization result is determined by the shape of domain boundaries, boundary-derived isolines (analogous to equipotential lines in physics or cartography) and perpendicular streamlines that define regions within the reconstructed coordinates system. In detail, digitalization of a spatial domain Ω is adapted by mapping the potential field in physics, i.e. a scalar field such as the electrical field, given boundary conditions. To be specific, a continuous Ω is digitized according to the gradient of a scalar variable ▽ψ (e.g. spatial layer/column values), which depends inversely on the distance of a given bucket **r** (a bin or a cell) to the target boundary Γ (as *∂*Ω). Such a scheme can be modeled by Poisson’s Equation:

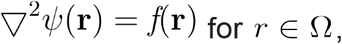

where *f* is the known function of **r** in the domain Ω. Importantly, the potential field *ψ* can only be solved when boundary conditions (BCs) are defined. The conditions are either specified by the bucket-wise values (called Dirichlet or class I BC) under certain known function *g*_1_,

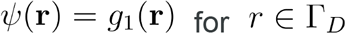

or by the normal derivative of the solution on the boundary (called Neumann or class II BC)

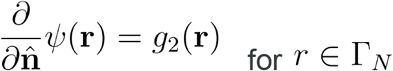

These BCs are crucial to determine *ψ* via iteration methods in numerical analyses, especially when the geometries of boundaries are complex and irregular.

In this study, we first digitalize the spatial domain with a steady-state field and then infer the effect of the true internal sources by highly variable genes along the potential gradients that violate the null hypothesis, thus it simplifies Poisson’s equation to its homogenous case (*f* = 0): Laplace’s equation. In the three-dimensional (3D) space, Laplace’s equation is written as:

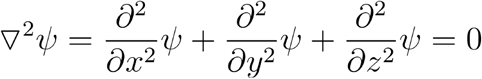

For visualization purposes, we first use the basic two-dimensional form to demonstrate the numerical solution with the Jacobi method under the supplied boundary condition,

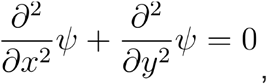

and further extend it to the 3D space.

Numerically, the equation is discretized as

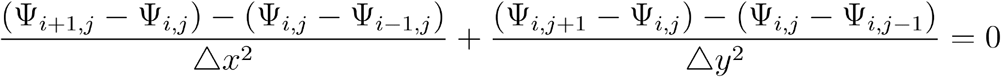

When we choose Δ*x* =Δ*y* for homogeneity in both *x,y* dimensions, the equation can be written as

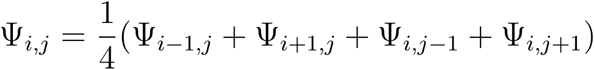

Now, the discrete form of Laplace’s equation shows that the potential of the central grid point is the equal-weighted average of four neighborhoods.

Based on the Jacobi method (Saad, 2003), for all interior grids at the (k+1)-th iteration, we have:

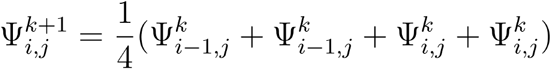

And for an enclosed domain, any neighbor of an interior grid would not fall out of the boundary, no matter how irregular the domain shape is **(Fig. 2g)**. The iteration process ends either with convergence when the normalized L2 loss

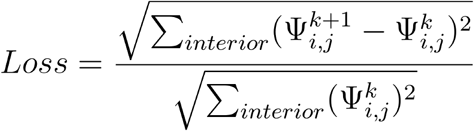

between two iterations is below 1e-5 by default, or when iterations surpass the maximum number of iterations (default 100000).

We then solve the real-world boundary problem. For any shape of a bounded domain, the boundary Γ can be arbitrarily divided into four connected boundaries with specified breakpoints **(Fig. 2h)**. With two sides, Γ_*D*_2__ and Γ_*D*_4__, respectively hold the fixed potentials as the Dirichlet BCs, the equipotential lines (the dotted lines with arrows) are naturally perpendicular to the Neumann BCs Γ_*N*_1__ and Γ_*N*_3__. Note that the normal derivative for an irregular boundary can not satisfy the numerical estimation, we project the original Neumann boundaries to two custom line segments, thus approximate the Neumann BC to Dirichlet BC with uniform distribution along these segments (or original boundaries, with slight loss of precision) for downstream analysis.

It is derived as

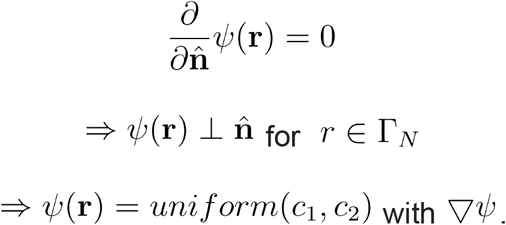

And

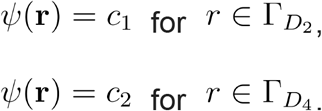

For numerical convenience, the grids that lay outside the potential field are set to value zero and *c*1 and *c*2 are symmetric about zero. By default, we set *c*_1_ = –1 and *c*_2_ = +1,

The final task becomes to solve *ψ*^1^ with the equation:

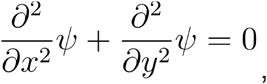

with the boundary conditions

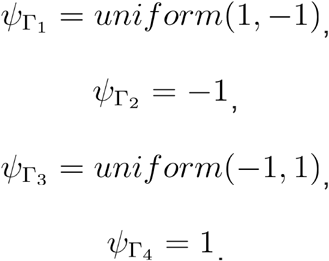

To hold these Dirichlet BCs during the iterative updating, the values of the boundary grids should be re-initialized during each iteration. This process is crucial to insulate the influence of any exterior grid when only the four closest neighbor grids are applied to calculate the value of the target grid in the Jacobi algorithm.

Once the solution of *ψ* is obtained (numerically, every grid has its potential value Ψ_*i,j*_), the domain is theoretically digitalized and can be partitioned into meshes of any equal-area size. To dissect the “layers”, the equipotential lines are generated by ligation of the grids with the same value from boundary Γ_*N*_1__ to Γ_*N*_3__ and evenly spaced with a constant interval. For the “columns”, we propose two approaches to handle the boundaries Γ_*D*_, with varying degrees of complexity. The first approach is the “one-step” approach.

When the boundaries are relatively regular or smooth (**Fig. 2i**), the concept of calculating streamlines that are perpendicular to the equipotential lines is workable. It is equivalent to calculating the tangent vector field,

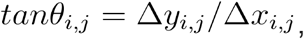

for each grid, and it is numerically solved by the Runge Kutta fourth order method (RK4) for approximating the solution of ordinary differential equations.

After the tangent field is constructed, the streamlines are computed by starting at any point of Γ_*D*_2__ perpendicularly crossing each equipotential line under the direction of the tangent field, and finally reaches the boundary Γ_*D*_4__. However, in practice, the RK4 method decreases its precision along with increment of the boundary irregularity.

The second approach is the “two-round” approach. In the first round of the digitalization, only equipotential lines are recorded as the result of domain layering. And the method then can be interpreted as rotating the domain 90 degrees, and repeating the whole process but orthogonally columning the domain in the second round. This approach is a rough but effective approximation that can be adapted in most real-world cases without any extra parameter settings.

The three-dimensional form of potential for each central point becomes:

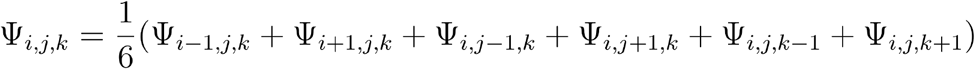

and the Jacobi method still works as long as the 3D domain is enclosed. However, the rate of convergence usually slows down when comparing the solution in two-dimensional space.

In Spateo we can use st.dd.digitize to digitize the spatial domain of interests into different layers or columns:

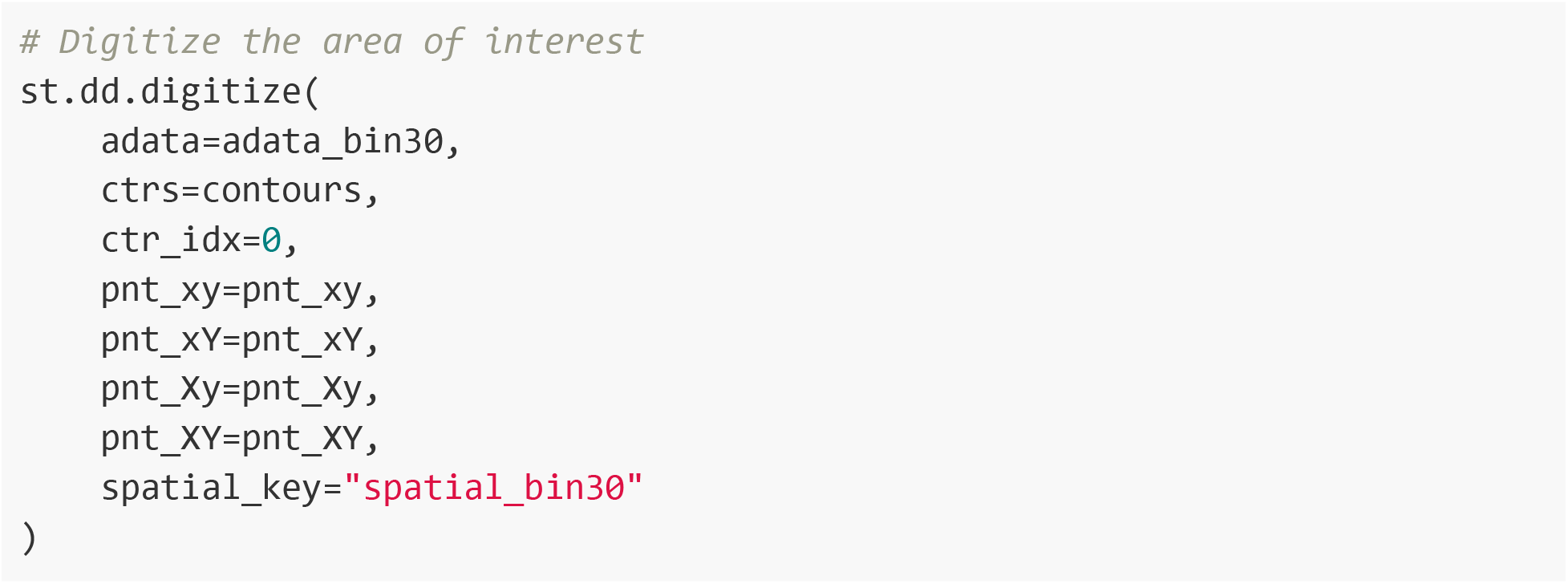

#### Construct cell type colocalization matrix

We use the ball tree algorithm to find the 10 nearest neighbors in Euclidean space for each spot, along with the distances to each nearest neighbor. The indices are used to construct a weighted adjacency matrix, where the magnitude of the entries are spot-to-spot distances. The sample indices are matched with the categorical labels of each sample, allowing for generation of a normalized one-hot matrix where each row corresponds to a categorical label (i.e. cell type) and is normalized by the count of the label in that row. Using the weighted adjacency matrix and the normalized one-hot matrix, we perform a congruence transformation to aggregate spatial weights for each cell type pair, returning an array describing cell type-by-cell type colocalization.

In Spateo, we use the following code to create a heatmap of cell-type colocalization:

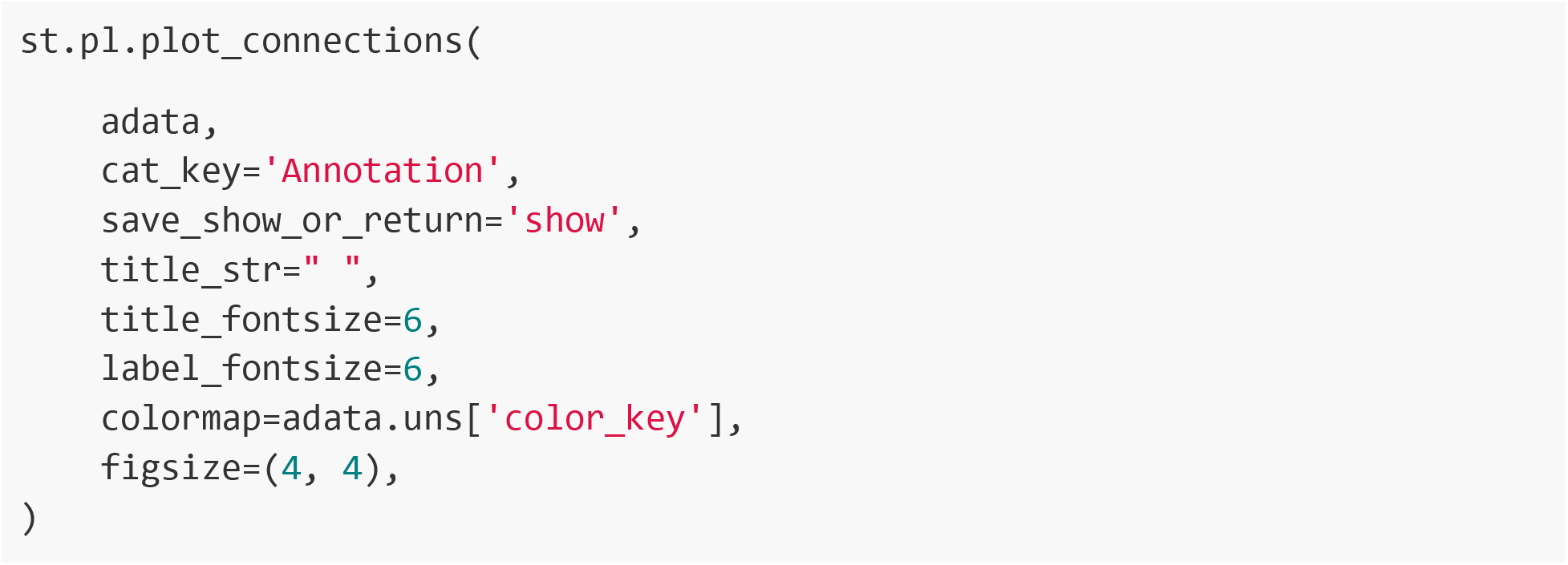

#### Investigate cell-type colocalization across different spatial scale

To investigate the spatial niches of a particular cell across different spatial scale, we first calculate the spatial neighbor cells of all the cells belonging to the cell type of interest. Then, we calculate the percentage of each cell type within the neighbors of each cell, which defines the niche composition of the cell. We can next calculate the mean and variance of the percentages across all cells. By increasing the neighbor size, we can inspect how the niche composition changes over different spatial scale.

#### Detection of spatially autocorrelated genes

To detect spatially autocorrelated genes, we posit the null hypothesis that the gene expression is spatially randomly distributed without any significant spatial enrichment. We use global Moran’s *I* to measure the spatial autocorrelation, which evaluates whether the gene expression is clustered (close to +1), disperse (close to −1), or random (close to 0). For each gene, the global Moran’s *I* index is defined as:

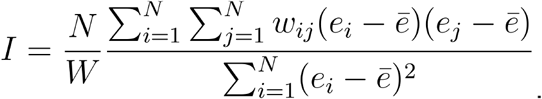

The formula assesses the relative concordance (simultaneous over- or under-expression to the mean *ē*) of expression of any two pairs of neighboring locations *i,j*. *w_ij_* is the spatial weight, set to 1 for pairs of neighbors and 0 otherwise. *W* is the sum of all spatial weights, and *N* is the total number of the spatial spots/bins of interests. The index is proven to follow the asymptotic normal distribution (Sen, 1977, 2010) and can be standardized to obtain the z-score and corresponding p-value.

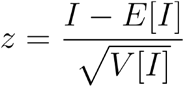

where:

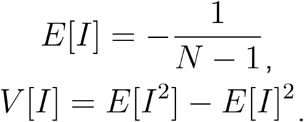

For the genes with statistically significant p-values (0.05 by default) and positive z-scores, the Benjamini-Hochberg (BH) correction is applied to control the overall false discovery rate (FDR, 0.05 by default). We also provide a permutation-based test(Fang et al., 2022) to estimate the *p*-value. We perform the permutation 200 times to generate a null distribution, and then determine the fold change between the measured Moran’s *I* score and the mean expected score from the permutations. By calculating the z-score for the observed Moran’s *I* score based on the null distribution, we obtain the *p*-value based on the permutations.

To evaluate the local spatial autocorrelation for each gene, we leverage the local Moran’s *I* index

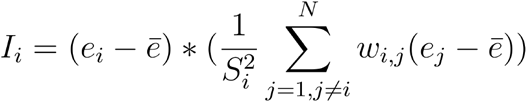

with the scaling factor

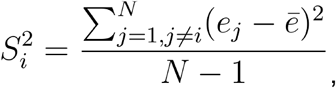

a local spatial autocorrelation metric that can identify cases in which the value of an observation and the average of its surroundings is either more similar (positive value) or dissimilar (negative value) than we would expect from pure chance. We bring additional information to improve the interpretability of local Moran’s I index. Firstly, we create a Moran plot(Anselin, 2019) by visualizing the expression of each gene in each bucket vs. its local average used in calculating the index. Next, we perform the same permutation test procedure to reveal the statistically significant buckets, followed by dividing those significant buckets into four quadrants, where the first quadrant (hotspot) indicates the gene is highly expressed in target bucket *i* and its neighbors with *w_i,j_* > 0(*w_i,j_* = 1 by default), the second quadrant (diamond) indicates the gene is highly expressed in the target bucket but surrounded primarily by low values, the third quadrant (cold spot) indicates the gene is lowly or not expressed in both the target bucket and surrounding buckets, and the fourth quadrant (doughnuts) indicates the gene is lowly expressed in the target spot but highly expressed in the surrounding.

To dissect each spatial domain with respect to each of the four quadrants, we first calculate, among all spatial domains, the maximal number, maximal fraction and maximal specificity of buckets belonging to any particular domain for any particular quadrant category. We then identify the spatial domain with highest fraction and similarly the spatial domain with highest specificity of buckets across all domains and for each quadrant. The identified spatial domain and associated quadrant are then used for downstream analysis and interpretation. With Spateo, spatially autocorrelated genes can be identified using

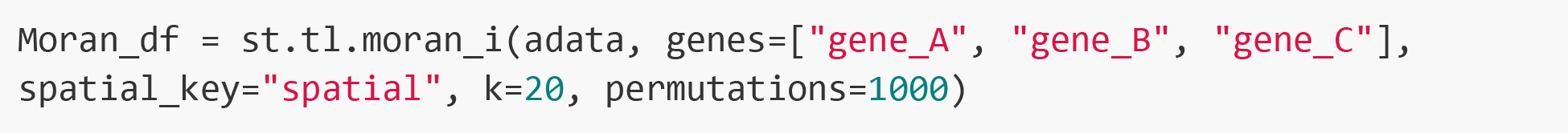

#### Calculate cell type spatial enrichment score via Moran’s *I*

We use the distribution of numbers of cells belonging to a particular cell type across all bins (bin size 60 for the hemibrain data) for spatial enrichment analyses. Specifically, we first assign each cell to a particular bin based on the identity of the bin that contains the centroid of the cell calculated during the segmentation procedure. By doing so, each cell will be uniquely assigned to a bin, but each bin may have more than one cell. Then, for the cell types of interest, we set the value for the centroids of all cells belonging to this cell type to be 1 and else 0. We next calculate the sum of the values of all centroids for each bin. The resultant values can then be used to calculate Moran’s I score for each cell type, similar to what is described in *Detection of spatially autocorrelated genes*.

#### Apply generalized linear models (GLM) to detect differentially expressed genes in different contexts

In this study, we used GLM as a general approach to identify genes significantly changing as a function of some continuous variables, such as the digitized layers / columns (Fig. 2G), the pseudospace defined by A-P axis or the principal curve learned for a particular 3D reconstruct organ, or the differential geometry quantities computed after learning the morphological vector field. In general, the full model of the GLM regression is:

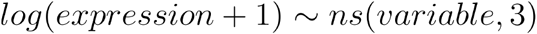

And the reduced model is:

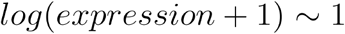

A likelihood ratio test is then used to compare these two models and to compute p-value. We BH adjust the P-value and define significant genes as genes with q-val < 0.05.

#### Ligand-receptor interaction prediction in spatial transcriptomics

A critical advantage of spatial transcriptome (ST) data over conventional scRNA-seq data is that ST preserves spatial information, allowing us to characterize the cells and their surrounding microenvironment and infer the complex interplay. A cell-type proximity matrix is firstly calculated to pinpoint spatially co-localized cell types, and prioritize cell-cell interactions between the cell types of interest. For the given couple of cell types, users should manually define the sending cell type (that expresses the ligands) and the receiving cell type (that expresses the receptors). The potential interacting cell pairs of these two cell types are then extracted from the physical *k*-nearest neighbors (KNN) graph (N=10 by default). Candidate ligands or receptors from the CellPhoneDB(Efremova et al., 2020) v3.1.0 database are selected when their expression levels are high (i.e. compared to all possible ligands/receptors, the ligand or receptor is among the top 20 highest expressed ligands/receptors, where 20 is used as the default cutoff) in the appropriate sub-cluster (short for spatial-proximal subset of ligand/receptor cell cluster). Subsequently, for a ligand-receptor pair, the interaction score is either calculated as the averaged product of ligand and receptor expression for all extracted cell pairs, or as the positive rate, the proportion of cell pairs that simultaneously express the ligand and receptor. To test the significance of the interaction score, a random sampling test is performed. In each sampling process, the same number of spatially-neighboring cell pairs are randomly selected from across the entire sample to calculate the interaction score for the given ligand-receptor complex. The random sampling process is repeated 1000 times by default, and the *p*-value is defined as the proportion of the interaction scores that are higher than the actual scores. The interactions with *p*-value <0.05 are considered statistically significant. Furthermore, the discovery of interactions can be additionally conditioned on the specificity of the ligands and receptors to the regions of sending and receiving cells that are close to one another. After computing the score for each ligand-receptor pair, the sender or receiver cell type of interest can be categorized as either being spatially proximal to or spatially distant from cells of the receiving or sending cell type, respectively (henceforth referred to as the “spatially proximal subset” and the “spatially distant subset”)- this information is available from the *k*-nearest neighbors graph (see above). The “specific” interactions can be identified from significant differential expression of the ligand (for sending cell types) and the corresponding receiver (for receiving cell types) in the spatially proximal subset compared to the spatially distant subset, where this identifier is assigned using a Wilcoxon rank-sum test. Thus interactions can be deemed to be “significant but not specific”, “significant and specific”, etc. Spateo contains reference databases for human, mouse, *Drosophila*, zebrafish and axolotl ligand-receptor interactions, which can be accessed for ligand-receptor interaction analysis by calling:

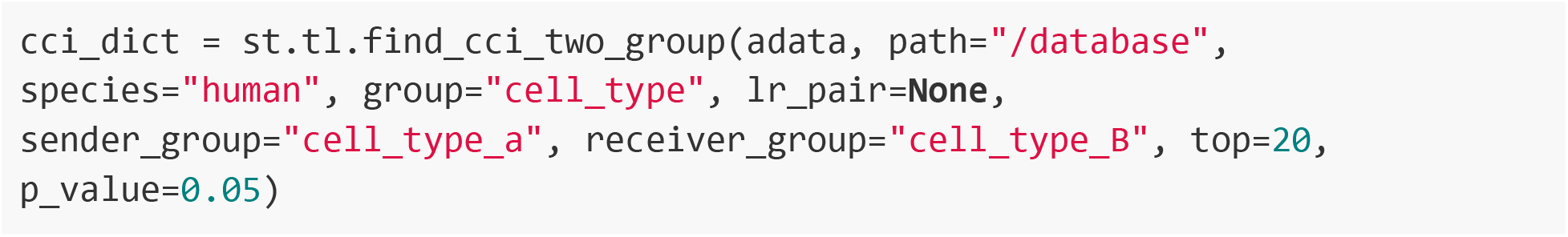

where *path* is the path to the folder containing ligand-receptor database(s); in the Spateo repository, these can be found in the “database” folder. *Group* is the key in the AnnData object that stores cell type labels, *Ir_pair* can be used to provide a list of ligand-receptor pairs of interest (where each takes the form “{ligand}-{receptor}”), and *top* is used to filter the list of ligands and receptors to consider down to the *top* genes based on expression in the cell types specified by *sender_group* and *receiver_group*.

#### Spatially-aware modeling of cell-cell interaction with spatial transcriptomics

Briefly, on the rationale behind the implementation of spatially-aware models: nonspatial methods assume that random errors of the regression are independent and identically distributed; i.e. the error for one cell is completely uncorrelated from the errors of its neighbors. As can be seen with our methods to measure the significance of spatial distributions of cell types, many genes are spatially dependent, such that spatially proximal pairs of cells will be more similar in many respects compared to spatially-distant pairs, violating the key assumption of spatially-unaware regressions. Significant spatial autocorrelation and cross-correlation are capable of driving spurious correlations(Ploton et al., 2020); to mitigate potential overestimations of model predictive power, for each cell we combine information from each of its neighbors to constitute the final feature set, with the architectures for the two classes of methods for doing so described below.

Spateo’s modeling suite comprises spatial lag models that consider neighboring values for endogenous and exogenous values as well as error terms, and two modifications to a generalized linear formulation for which each exogenous variable aggregates information from its neighbors, in which each endogenous (dependent) term is modeled by a log-linear distribution and each exogenous (independent) variable represents a factor for which the endogenous variable is assumed to be dependent on to some degree. This dependency is parameterized via a maximum likelihood estimation procedure in which the model takes the following general form for each gene ***P***:

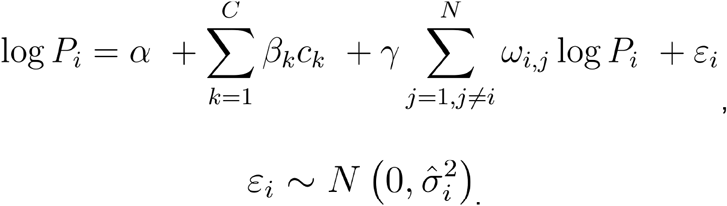

Where the generic *c_k_* represents an array of any spatial variates or covariates, for example representing the *k* – *th* spatial domain in the form of a binary one-hot dummy variable, a probabilities vector obtained using a cell type deconvolution algorithm, or the expression of a gene of interest. *c_k_* can represent cell-specific molecular attributes, as is the case with the spatially lagged model, or can incorporate information from the local neighbors through matrix operations, as is the case with both the niche composition and niche interaction models introduced in the following sections. Optionally, the γ term describes the contribution of spatial lag of the dependent variable, where *w_i,j_* is the spatial weight between cell ***i*** and cell ***j*** in its local neighborhood. Here *ε_i_* is the error term, assumed to be a white noise represented by a Gaussian distribution with zero mean and unit variance.

##### Infrastructure

###### Nearest neighbor graph

We represent the spatial neighborhood of each cell via a weighted undirected graph *G*(*V, E*), where each vertex *v* ∈ *V* represents a cell and each other vertex is connected to *v* by an edge with weight dependent on the distance between the two vertices, computed using the spatial coordinates of each cell. Programmatically, this is encoded as a weighted adjacency matrix. From this graph, we define the neighborhood of each cell in terms of its *k* nearest neighbors, where *k* is either a fixed user input or can be computed by finding all neighbors within a radius defined by user-specified parameters *p* and *sigma*, where *p* is a proportion of the maximum radius and *sigma* is the standard deviation of the Gaussian distribution that spatial weights are assumed to decay following. The fixed-neighbor and fixed-radius methods for nearest neighbor definition can be implemented as follows, respectively:

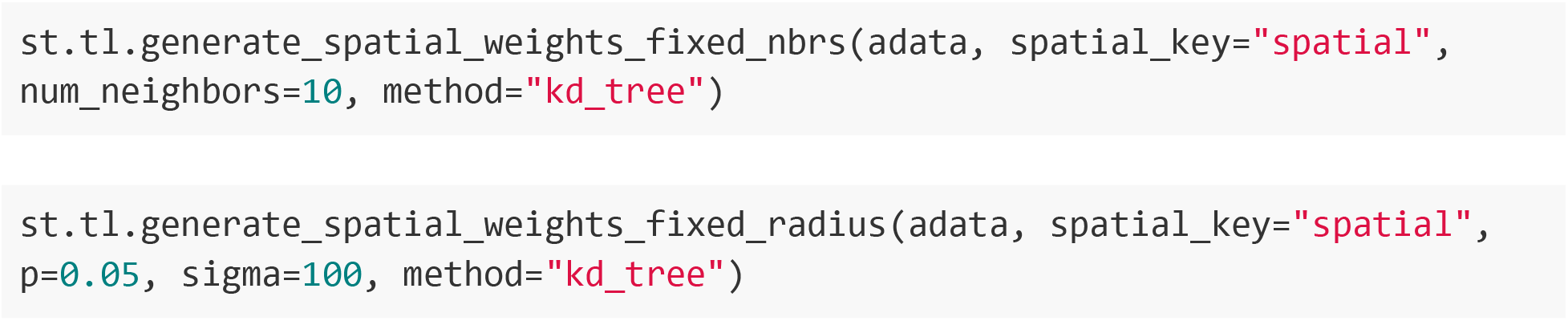

The spatial graphs and spatial distances are saved to adata.obps[‘spatial_connectivities’] and adata.obsp[‘spatial_distances’], respectively. By default, the 10 nearest neighbors are selected to build the graph.

##### Spatially-aware models

Multiple potential sources of technical variation lead to overdispersion and high dropout rates in single-cell data, producing characteristic right-skewed count distributions that result in inferential underpowering of general linear models due to an inability to reasonably assume normality of the raw or log-transformed counts. Probabilistic approaches have typically been used instead, fitting data at the level of reads and UMIs using a negative binomial(Grün et al., 2014), zero-inflated negative binomial(Eraslan et al., 2019; Lopez et al., 2018; Risso et al., 2018), or Poisson distribution(Cable et al., 2022a, 2022b; Kharchenko et al., 2014). Note that Spateo’s modeling core is capable of generalized linear regression following a variety of distributional assumptions, including Poisson, Gaussian, and negative binomial, however a model following the Poisson assumption will be detailed here. For each gene of interest, we aim to model *Y_i,j_*, the observed RNA counts for cell ***i*** and gene ***j***. We assume Poisson sampling such that

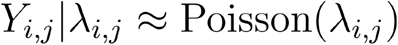

with expected transcript count *λ_i,j_*. Here *λ_i,j_* is a mixture of ***k*** explanatory variables *x_k_* ∈ ℝ^*k,j*^ for each gene ***j***, defined generally by

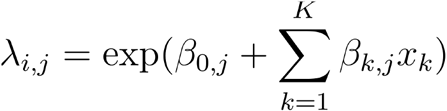

where each predictor vector *x_k_* for variable ***k*** is parameterized by linear coefficient *β_k_,k* = {1,2,…, *K*}. In general, each variable ***k*** can take many forms, for example:

1. Indicator variable, as is the case for the **Niche model**.In this case, *x_k_* is always either 0 or 1, representing e.g. a classification into a particular cell type among a set of possible cell types (among any other possibility involving categorical labels).
2. Discrete variable, as is the case for the category regression model included in Spateo. In this case, *x_k_*, can be 0 or any other integer, representing e.g. the number of cells of a particular category within the local neighborhood of each cell (among any other possibility involving categorical labels).
3. Continuous variable, as is the case with the cell-type specific ligand-receptor regression model described in the **SLICE** section. In this case, *x_k_* can take on continuous values representing e.g. gene expression.

In the context of our Poisson regression, each coefficient *β_k,j_* can be interpreted as the log fold-change of gene expression per unit change in *x_k,j_* for variate ***k*** and gene ***j***.

We additionally introduce an optional stabilizing nonlinear rectification(CIRANO. and Dugas, 2002) to prevent the explosion of λ when parameters are large, selectable by choosing “softplus” as the argument for the “distr” parameter. With this modification, *λ_i,j_* is instead defined as

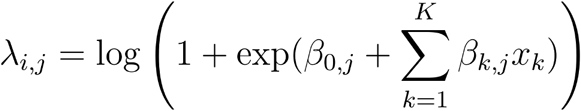

where the constant term 1 ensures numerical stability. For model fitting, we aim to parameterize the Poisson distribution such that the probability of observing given transcript count *Y_i,j_* is maximized; this can be achieved by minimizing the negative log-likelihood function for a Poisson distribution

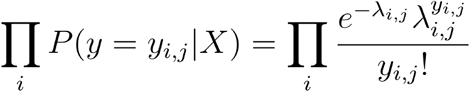

with respect to parameters *β*^0^ and *β^j^*, given by

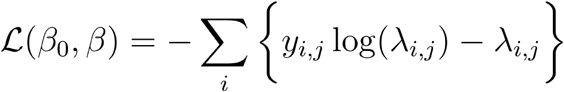

for each gene *j*.

Additionally, as niche-based models are prone to fitting large numbers of features due to the consideration of all possible combinations of categorical labels within a given cellular niche, for interpretability and to eliminate redundant information from super-collinear futures we constrain these models to be sparse, such that a relatively small number of relevant features are identified under the assumption that not all features provide substantive contributions to each component. Model parameterization is guided by a penalization of the log-likelihood, where the penalty term depends on the size of each parameter. Specifically, we introduce an elastic net penalty capable of interpolating between the Lasso(Tibshirani, 1996) and Ridge(Hoerl and Kennard, 1970) penalties with hyperparameter *α*:

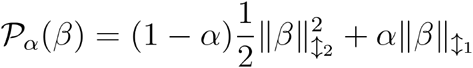

where *β* is the parameter vector except the intercept term *β*_0_. When *α* is zero, the model will use the Ridge penalty, and when *α* is one the model will use the Lasso penalty. The elastic net penalty and log likelihood function are additively combined to result in the objective function, which is minimized following a batch gradient descent procedure,

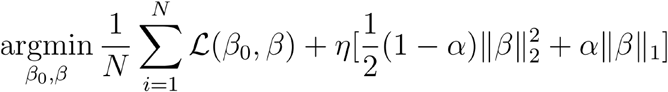

where *η* is the L1 regularization parameter (referred to as “lambda” in code but changed here as lambda is used for the Poisson conditional intensity function) and is discovered at runtime by choosing the optimal value following a *k*-fold cross validation procedure. *k* = 3 was used for the analysis shown in **Figure 3.** For each test set, any number of provided values of *η* along a descending regularization path *hmax* ≥ … ≥ *η*_0_ are tested, initializing successive models with the solution of the previous model for rapid convergence. The *η* resulting in the highest pseudo-R^2^ value is chosen, where the pseudo-R^2^ value is given by

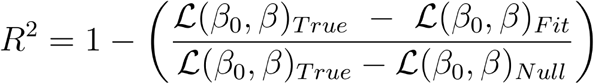

where 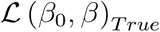 is the log-likelihood of observing the true transcript counts given a theoretical parameterization that would perfectly fit the target distribution, 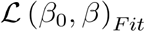 is the log-likelihood of observing the true counts given the parameterization that we obtain from our fitting process, and 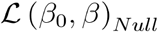 is the log-likelihood of observing the true counts given a null model parameterization that maps each set of data to the mean of the target distribution.

Additionally, a list of values can be given for any regression parameter, and a grid search will identify the optimal value for each parameter given in this way.

###### Niche model

Cell-cell interactions are indirectly connected to phenotypic changes, which are mediated by subsequently coordinated changes in gene expression in receiving cells. To elucidate these effects, an understanding of differential expression conditioned on cell-cell communication is required. However, it is difficult to attribute gene expression patterns to any singular ligand-receptor interaction, difficult to disentangle the contributions of multiple ligand-receptor interactions and difficult to adequately explain the variance in any gene expression pattern given only what we already know about ligand-receptor interactions, necessitating a method capable of incorporating both known ligand-receptor interactions and any unknown interaction mechanisms. To do so, we note that as a result of the quantifiable change in receiver gene expression directly resulting from ligand-receptor interaction, a partial statistical dependency is induced, of a component of the receiving cell’s gene expression profile on all aspects of the sending cell’s gene expression profile. From this partial dependency, patterns may emerge in which the marker genes of the sending cell tend to be more closely associated with gene expression patterns induced in the receiving cell, an association that can be measured and modeled. With the niche model, we propose to leverage these dependencies, hypothesizing that the identity of a cell’s closest neighbors can be used to predict downstream signaling effects. The design matrix of this model and the cell type-specific interaction model contain variables parameterized by coefficient *β_g,j_* and can be interpreted as the gene expression change across cells per unit change of each variate ***g*** for gene ***j***. The inputs are a gene expression matrix *Y* ∈ ℝ^*N*×*G*^, where ***N*** is the number of cells and ***G*** is the number of genes, a matrix of observed cell types *X_l_*, ∈ ℝ^*N x L*^ where ***L*** is the number of distinct cell type labels and an adjacency matrix *A* ∈ ℝ^*N*×*N*^, which is calculated based on the spatial proximity of cells and furthermore binarized such that *a_i,j_* = 1 if *d*(*x_i_,x_j_*) > *δ_max_*, where *d* describes the Euclidean distance between any two nodes *i,j* ∈ *N* and *δ_max_* is the user-defined maximum allowable distance between neighboring nodes, and *a^i,j^* = 0 otherwise.

To generate the design matrix, we compute discrete target cell interactions as the matrix product of the adjacency matrix and the cell type identity array *X_l_* similarly to the process used for NCEM(Fischer et al., 2021), resulting in

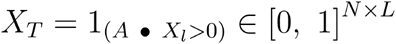

where ^1^(•) represents an indicator function. *X_T_* can be interpreted as a specification of the presence of cell type ***L*** within the local neighborhood of each sample. To generate a matrix representation of source-target cell interactions, we compute the interaction between each row of *X_l_* and each row of *X_T_* via the matrix product of the transposition of *X_l_* and *X_T_* followed by an optional thresholding step such that the features then become indicator functions representing the presence of at least one interacting pair of cell types within the niche of any given cell. The resulting interaction matrix is then taken as the complete design matrix, given by

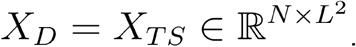

Overall, a particular gene’s expression ***Y***_*i, j*_ for cell ***i*** and gene ***j*** can thus be modeled as

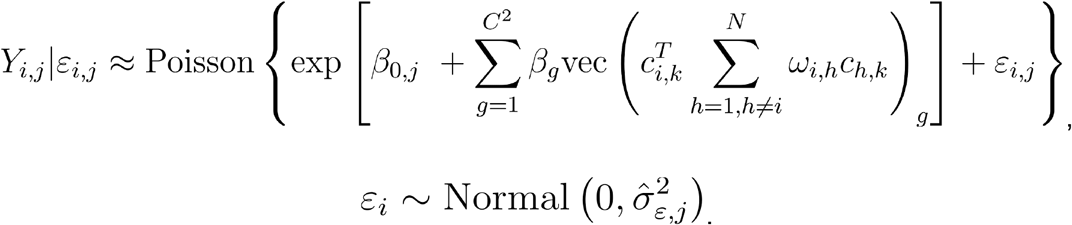

where *c_i,k_* is the one-hot cell type (or other category) vector for the ***i***^*th*^ cell with ***k*** possible categorical classifications,is the one-hot cell type vector of the ***h***^*th*^ neighboring cell with ***k*** possible categorical classifications, *w_i,h_* is the spatial weight between cells ***i*** and ***h*** (the weight of the appropriate edge in the spatial graph), “vec” is a flattening operation and *β_g_* is the ***g***^*th*^ variate, here representing the ***g***^*th*^ cell type-cell type pair as well as the ***g***^*th*^ column of the product in the innermost parenthetical term. This product can also take the form of an indicator array, where values are 1 if a given cell type-cell type pair can be observed and 0 if it cannot. We are thereby able to predict RNA count values using a spatially-aware generalized linear model, considering the suboptimality of the normal distribution assumption for scRNA-seq data(Hafemeister and Satija, 2019; Townes et al., 2019).

#### Spatially-aware Ligand:receptor-based Inference of Cell type-specific Effects (SLICE) model

Although useful to infer the correlates and potential consequences of interaction, the niche regression model cannot inform which ligand(s) and receptor(s) likely caused the observed expression. Prediction of precise downstream effects is confounded by the ability of promiscuous receptors to bind multiple ligands and *vice versa*, necessitating analysis methods capable of unraveling the contributions of each individual ligand-receptor pair. For this purpose, we introduce a parametric framework combining spatial, categorical and ligand-receptor expression information to estimate downstream effects of specific interactions. This model utilizes the categorical identities of ligand-expressing sender cells and receptor-expressing receiver cells as covariates of the expression of specified ligands and receptors, such that the predicted effects are additionally conditioned on qualities of interest, e.g. cell type.

Users can supply either a list of ligands of interest and a list of additional genes of interest to serve as the predictors and targets of the regression model, respectively, or can supply only the list of ligands, in which case a knowledge graph is used to automate the selection of targets from prior knowledge signaling networks. Ligand-receptor interactions from CellTalkDB (Shao et al., 2021), pathways from the Kyoto Encyclopedia of Genes and Genomes (KEGG) (Kanehisa and Goto, 2000) and Reactome (Gillespie et al., 2022), and transcription factors from AnimalTFDB (Hu et al., 2019) were integrated to construct this signaling network.

Similarly to the **Niche model**, the inputs used to construct the design matrix are a gene expression matrix *Y* ∈ ℝ^*N x G*^, a matrix of observed cell types *X_l_* ∈ ℝ^*N x L*^ and a binarized adjacency matrix *A* ∈ ℝ^*N x N*^. Given ***m*** ligands and optionally ***r*** receptors, all possible interacting pairs are identified using the CellPhoneDB(Efremova et al., 2020) database (v3.1). We amend the cell type identity array *X_l_* with the expression of each ligand and the expression of each receptor, resulting in

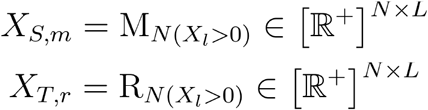

respectively, where ***m*** and ***r*** refer to the ***m**^th^* ligand and ***r**^th^* receptor and *M_N_* and *R_N_* are ligand and receptor expressions in the ***N***^*th*^ sample. Optionally, we use the spatial lag of *R_N_* to compute *X_T,r_*, where each element in *R_N_* is instead a weighted average of the expression of receptor ***r*** computed over its local neighborhood. Similarly to the procedure for the **Niche model**, we compute the interaction between each row of ***X***_S,m_ and each row of ***X***_*T,r*_ via the matrix product of the transposition of ***X***_*S,m*_ and *X_T,r_* This process is repeated for all combinations of ligands and receptors identified, and the resulting arrays are combined to form the complete design matrix, given by

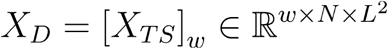

where ***w*** denotes the combinations identified. Overall, a particular gene’s expression *Y_i,j_* for cell ***i*** and gene ***j*** can thus be modeled as

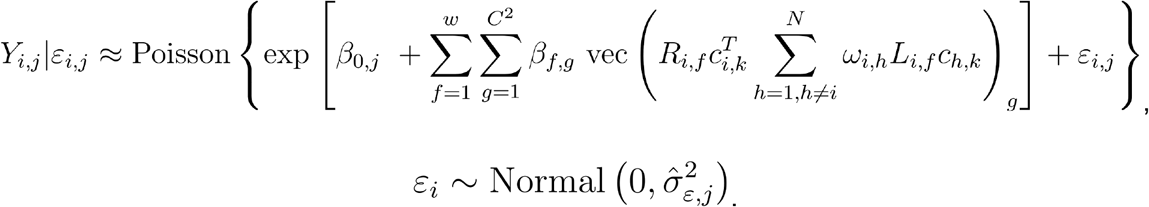

where *β_f,g_* is the ***g***^*th*^ variate for the ***f***^*th*^ ligand-receptor pair, here representing the ***g***^*th*^ cell type-cell type pair as well as the ***g***^*th*^ column of the product in the innermost parenthetical term, *L_i,f_* is the expression of ligand ***f*** in cell ***i***, *R_i,f_* is the expression of receptor ***f*** in cell ***i*** and all other quantities are as described for the **Niche model**.

For this model as well as the niche model, the regression can be constrained on the basis of any preselected subset of categorical groupings. An example of class instantiation and gene expression prediction is included below:

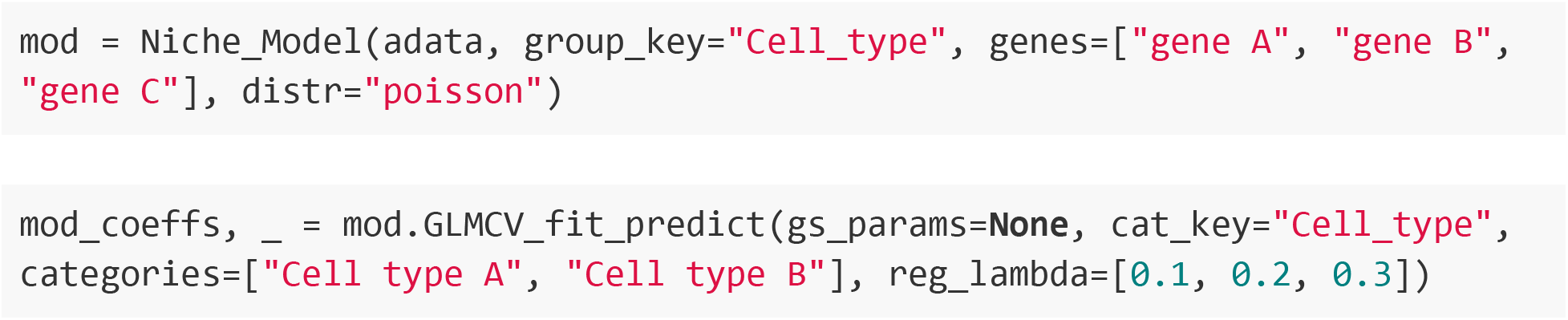

where “categories” sets the aforementioned subset of categorical groupings, and “cat_key” is used to specify the key in adata.obs where the categorization is defined. Note also that “reg_lambda” is used to provide a list of possible regularization coefficients and “gs_params” is an optional dictionary with keys corresponding to arguments for the “GLM” class, used to pass lists of values to designated arguments to initialize a grid search that will find the optimal combination of these hyperparameters. The optimal lambda and optimal combination of hyperparameters is decided by selecting the fitted model with the highest pseudo-R^2^ value. For the analyses in **figure 3**, we allowed lambda to vary between three values on a log scale between 0.001 and 10^-5^.

#### Spatially-lagged models

Single-cell RNA sequencing data is frequently transformed before analysis (e.g. through a logarithmic transformation) such that the distribution of values more closely approaches a Gaussian and so that other methods that assume normality can be used. For this case, general linear regression can be used also in estimation of gene-level or niche-level effects, with the consideration that the model should also consider the spatial dependencies between samples and variables. For this purpose, we are able to consider the exogenous lag (the values of the independent variables in the neighborhood of each sample) as instruments for the regression, and can introduce a spatially lagged linear regression model(Rey and Anselin, 2010), where the expression *Y_i,j_* for cell ***i*** and gene ***j*** can be modeled generally using **equation 1**, where the spatial lag term *γ* is nonzero and determined by model fitting. For receptor genes of interest (typically chosen by hypothesizing that these genes may be associated with particular ligands of interest), Spateo can determine the relationship between expression of these receptors, expression of these receptors in the neighboring cells and expression of ligands in the neighboring cells by fitting the following model:

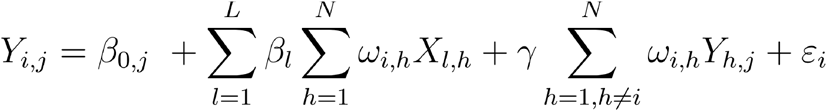

where *Y_i,j_* is the log-transformed (or any other equivalent normalizing transformation, e.g. variance scaling) variant of the gene expression, *X_l,h_* is the identically-transformed expression of ***ℓ*** genes of interest (stylized as ***ℓ*** here to refer to multiple ligands as per the inputs to the model), and the exogenous lag (the expression of gene ***j*** in the local neighborhood of the site ***i***, here represented as the weighted summation of all neighboring cells ***h***) is a component of the estimation, with its dependency quantified by coefficient *γ*. Similarly to the **SLICE model,** if target genes are not provided, Spateo’s database is used to automate the selection of likely-linked targets for the given ligands.

We additionally extend this idea to the **Niche model**, noting that simultaneous information about the cell type colocalization patterns and gene expression patterns of neighboring cells may be useful in eliminating coincidental patterns that may not be biologically meaningful. This takes the form:

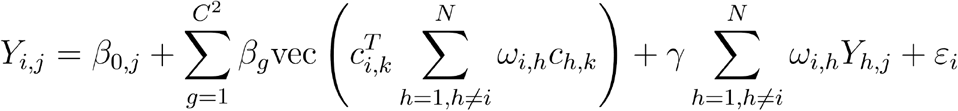

where *Y_i,j_* is again the log-transformed (or any other equivalent normalizing transformation, e.g. variance scaling) variant of the gene expression, “vec” is a flattening operation and *β_g_* is the ***g***^*th*^ variate, here representing the ***g***^*th*^ cell type-cell type pair as well as the ***g***^*th*^ column of the product in the innermost parenthetical term.

For both models, the coefficients *β_g_, γ* and the intercept are estimated by the two-stage least square method implemented in Pysal’s *GM_lag()* function. Each of these models can be instantiated from the same class:

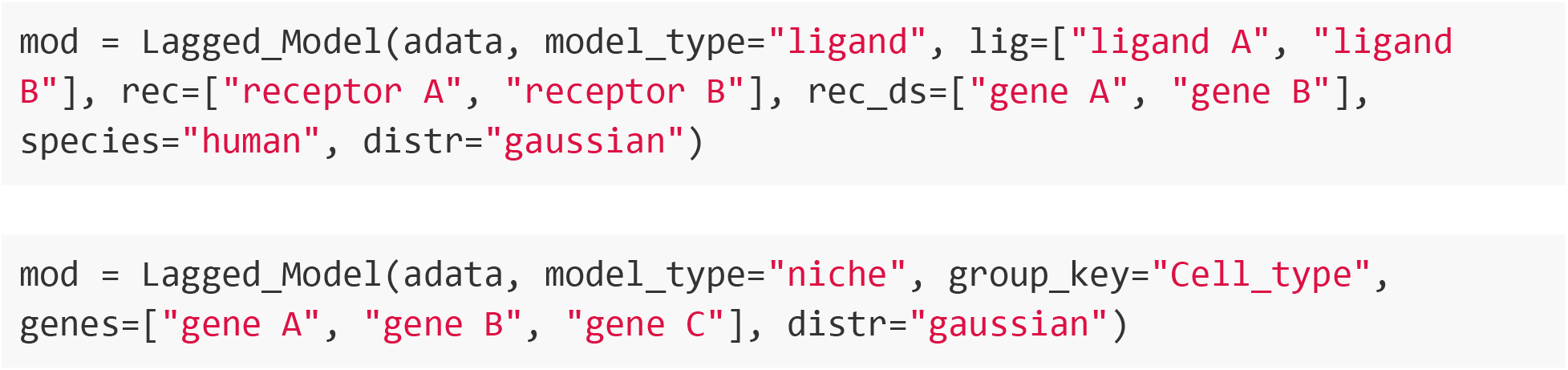

From here, the model can be fit using the same function call:

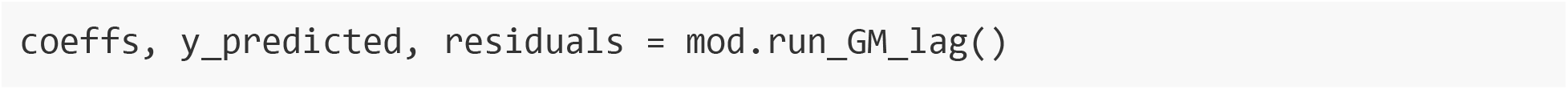

#### Computation of sender-receiver effects

For the all spatially-aware and spatially-lagged models, we use hypothesis testing to determine significance for each interaction parameter following estimation of the parameter vector *β_k,g_* for variate ***k*** for gene ***g***. By the principle of asymptotic normality(van der Vaart, 2000), we can approximate the deviation of the estimated value from the unknown true parameterization by

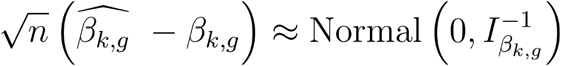

where ***n*** is the total number of cells and 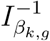 is the inverse Fisher information matrix that measures the amount of information an observable random variable ***X*** carries about parameter *β* upon which the probability of ***X*** is dependent. The standard error of *β_k,g_*, denoted *S_k,g_*, is

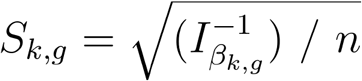

Knowing the value of the parameter as well as its standard error, we can compute the *z*-statistic:

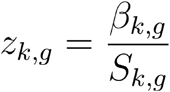

Using a two-tailed *z*-test, we compute a *p*-value for the null hypothesis that as

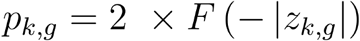

where *F* is the standard normal distribution function. To assign “significant” labels, *q* values are calculated across all features to control the false discovery rate (FDR) using the Benjamini-Hochberg procedure(Benjamini and Hochberg, 1995), using a default FDR of 0.05. Following computation of sender-receiver effects, we can set cutoffs for effect size and FDR-corrected *p*-value to identify genes for which the predicted effect indicates a significant change in the presence of the niche factors used as regressors, with 0.4 and 0.05 as the respective default values.

#### Selection of regression targets

For the axolotl brain data, several gene sets were used as regression targets. The gene set in **Fig. 3i** and **Fig. S3** comprises all genes with nonzero expression in >5% of cells in addition to Moran’s *I* coefficient that is significant and >0.4. The gene set in **Fig. S4** consists of genes with function in glial differentiation and axonogenesis, Wnt/Notch signaling, and translation initiation, as determined from gene ontology (GO) annotations. The gene set displayed in **Fig. S5** consists of all genes expressed in >5% of cells and which had a significant coefficient with reaEGC cells as the receiver cell population.

#### Spatial RNA velocity vector field analyses of mouse heart cell atlas

For the heart cell atlas dataset, we identified cell types across batches and timepoints by computing Pearson residuals and performing Harmony (Korsunsky et al., 2019) de-batching, Louvain clustering and finally marker gene detection with differential gene expression analyses. RNA velocity estimation and RNA velocity vector field analysis were performed using Dynamo with default parameters.

#### The Stereo-seq experiment

##### Tissue collection

The methods related to laboratory animal experiments presented were approved by the Institutional Review Board of the Ethics Committee of BGI (Permit No. BGI-IR20210903001). Mouse brain was dissected from adult C57BL/6J mice. First, the mice were sacrificed via carbon dioxide asphyxiation. Then the whole brain was harvested and immediately immersed in embedding molds with pre-cooled Tissue-Tek OCT (Sakura, 4583). Next the mold with tissue was snap-frozen in liquid nitrogen prechilled isopentane until the OCT was completely solid and then transferred to a −80 °C freezer for storage before the experiment. The embedded whole mouse brain was cut coronally with a thickness of 10 μm using a Leika CM1950 cryostat.

*Drosophila* samples used in this study were from *Drosophila* strain w1118. Flies were maintained on the cornmeal-sucrose-agar media at a constant temperature of 25°C in an incubator with 12 h/12 h light/dark cycle. Embryos of desired ages were collected from the grape juice plate (2.15% w/v agar, 49% v/v grape juice, 0.2% v/v propionic acid, 0.02% phosphate acid). Designated embryos were picked and observed under a microscope to verify their developmental stages. Before embedding, samples were stained and washed following methods in the previously published paper(Wang et al., 2022). We immobilized samples with double-sided tapes and adjusted the orientations of each sample to cut them along the left-right axis for the sagittal section. *Drosophila* embryos used in this study were embedded and sectioned together. Cryosection was performed with 7-μm slice thickness.

##### Stereo-seq experiment and spatial transcriptomic sequencing

Tissue sections were applied to Stereo-seq chips immediately after cryosection. To reduce batch effect, multiple cryosection slices of *Drosophila* embryos were subjected to one 1 x 1-cm Stereo-seq chip. The mouse brain section was subjected to one 1 x 1-cm Stereo-seq chip.Stereo-seq library was constructed and sequenced following the Stereo-seq protocol (https://db.cngb.org/stomics/flysta3d/stereo.seq.html). Briefly, sections on Stereo-seq chips were fixed in methanol at −20 °C for 40 min. After removing the methanol, chips were incubated in the tissue fluorescence staining solution (1/200 InvitrogenTM Qubit ssDNA HS Reagent and 2 U/μl RNase Inhibitor in 5× SSC) for 5 min. Tissue fluorescence staining solution was then removed and the chip was washed with 100 μl 0.1× saline-sodium citrate (SSC) buffer. After taking fluorescence images from the chip, sections were permeabilized on chip with 100 mL 0.1% pepsin in 0.01 M HCl at 37 °C for 5 min. Permeabilization solution was then removed and the chip was washed twice with 100 μl 0.1× SSC buffer. For cDNA generation, 100 μl RT buffer (10 U/mL reverse transcriptase, 1 mM dNTPs, 1 M betaine solution PCR reagent, 7.5 mM MgCl2, 5 mM DTT, 2 U/mL RNase inhibitor, 2.5 mM Stereo-seq-TSO and 1x First-strand buffer) was added on the chip and then the chip was maintained in a 42 °C incubator for 1 h. RT buffer was then removed and the chip was washed twice with 100 μl 0.1× SSC buffer. Tissue removal was performed by incubating the chip in tissue removal buffer (10 mM Tris-HCl, 25 mM EDTA, 100 mM NaCl, 0.5% SDS) at 37 °C for 30 min. Tissue removal buffer was then removed and the chip was washed twice with 400 μl 0.1× SSC buffer. cDNA was released from the chip by incubating the chip in 400 μl release buffer (5% Exonuclease I and 1× Exo I Reaction Buffer in H2O) at 55 °C for 3 h. Released cDNA was purified with 0.8× XP clean beads (Beckman Coulter) and then amplified with KAPA HiFi Hotstart Ready Mix with 0.8 mM cDNA PCR primer. The PCR product was purified using 0.7X XP beads. For each chip, 20ng cDNA was sheared with in-house Tn5 transposase and then amplified and purified. Final libraries were sequenced on a MGI DNBSEQ-Tx sequencer.

##### Reconstruction and analyses of whole-body 3D models of tissues and organs

Most existing spatial transcriptomics methods offer 2D spatially resolved transcriptomes but tissue, organ and embryo are 3D identities that have unique spatial structure. Leveraging the ultra-high field of view of Stereo-seq, we can perform continuous slicing of the same tissue or organ (such as *Drosophila(Wang et al., 2022*)or even mouse embryos(Chen et al., 2022)). However, after the slicing and sequencing, the original coordinates of the cells are lost. Spateo builds upon the PASTE algorithm(Zeira et al., 2022) to align different slices to create aligned 3D point clouds, from which we can build 3D models and perform various downstream morphometric analyses.

##### Align serial slices by fused-Gromov–Wasserstein (FGW) optimal transport and solving generalized weighted Procrustes, based on the PASTE algorithm

Assume we have spatially-resolved, serial continuous slices profiled with Stereo-seq and others, represented as 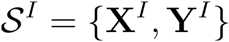, where *I* ∈ {1,2,…, *t*} denotes *t* profiled slices, *Y* ∈ ℝ^*N_I_* x *D*^ denotes the spatial coordinates in two dimensional space (*D* = 2) of spots in slice *I*, and *X^I^* ∈ ∝^*N_I_* x *G*^ is the measured readout features (a vector of gene expression) at these buckets (representing either cells, binned grids, or spots). In general, instead of using the spatial coordinates, we use the distance matrix **D**^*I*^, where *D_ij_* ||*y,y_.j_*||, to represent the spatial information with the advantage of being invariant to translation or rotation of the slice(Zeira et al., 2022). Our goal is to align these slices sequentially to reconstruct a 3D model of a tissue, organ or an entire embryo based on the matching of the gene expression pattern and spatial localization between buckets across slices. Spateo builds on the PASTE algorithm(Zeira et al., 2022) to align the slices with fused-Gromov–Wasserstein (FGW) distance, which extends the Wasserstein and Gromov–Wasserstein distances to encode the gene expression and spatial information in learning the optimal transport. Specifically, for every two consecutive slices, *i,i*’ containing *n,n*’ buckets, over a common set of *m* genes expressed across all slices, we want to find a optimal mapping Π = *τ*(*b,b*’), following the transport cost (*c*) and spatial preservation(Zeira et al., 2022):

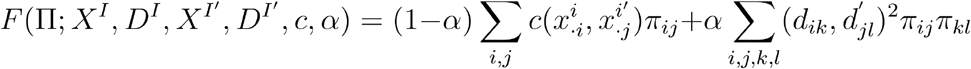

Intuitively, the above equation implies that the optimal mapping will ensure most similar cells across slices will be aligned while two cells nearby in one slide will be mapped to two other cells also nearby in another slice. With the optimal transport matrices Π learned, we calculate the rotation and translation matrices (*R,t*) sequentially, starting for the first two slices, to project to all slices to the same coordinate system by solving a generalized weighted Procrustes problem(Zeira et al., 2022):

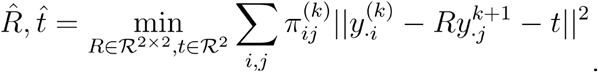

Setting the coordinates of the first slice as the reference, Spateo next sequentially applies the rotation (*R*) and translation (*t*) matrix to project all slides into the same 2D coordinate system. After appending the coordinates from the third dimension for all buckets of the same slice with the physical depth along the slicing axis, Spateo returns the aligned 3D coordinates of all buckets in 3D space.

In practice, we find that using the binning grids to learn the optimal transport and the rotation and translation matrices often improves the accuracy and robustness of the alignment probably because single cell segmentation is imperfect and noisy, while binned grids enable us to focus on the global mapping, as demonstrated below:

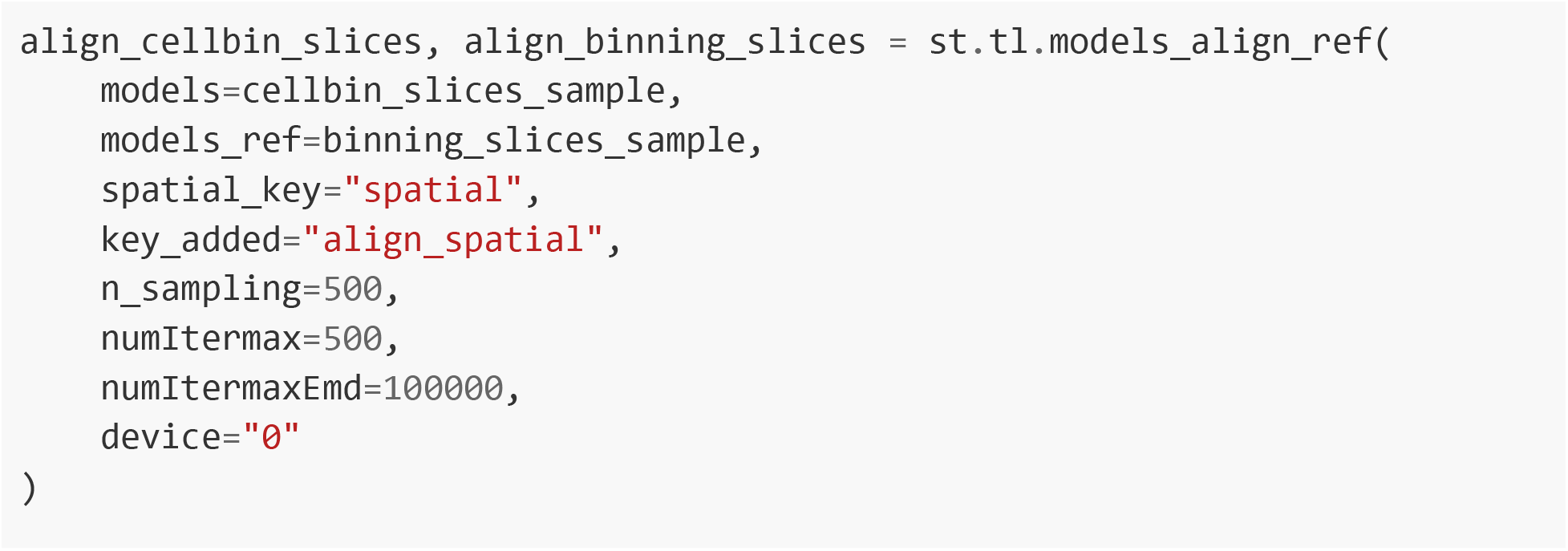

##### Create surface polygon mesh, volumetric mesh and voxel models of a whole embryo, or individual organs from 3D point clouds with PyVista

The key innovation of Spateo over PASTE is its ability to perform various downstream 3D modeling and morphometric analyses with the 3D aligned points clouds. In particular, we leverage the PolyData data structure from PyVista to represent the 3D point cloud, where each cell is annotated with the tissue identity, using the following function:

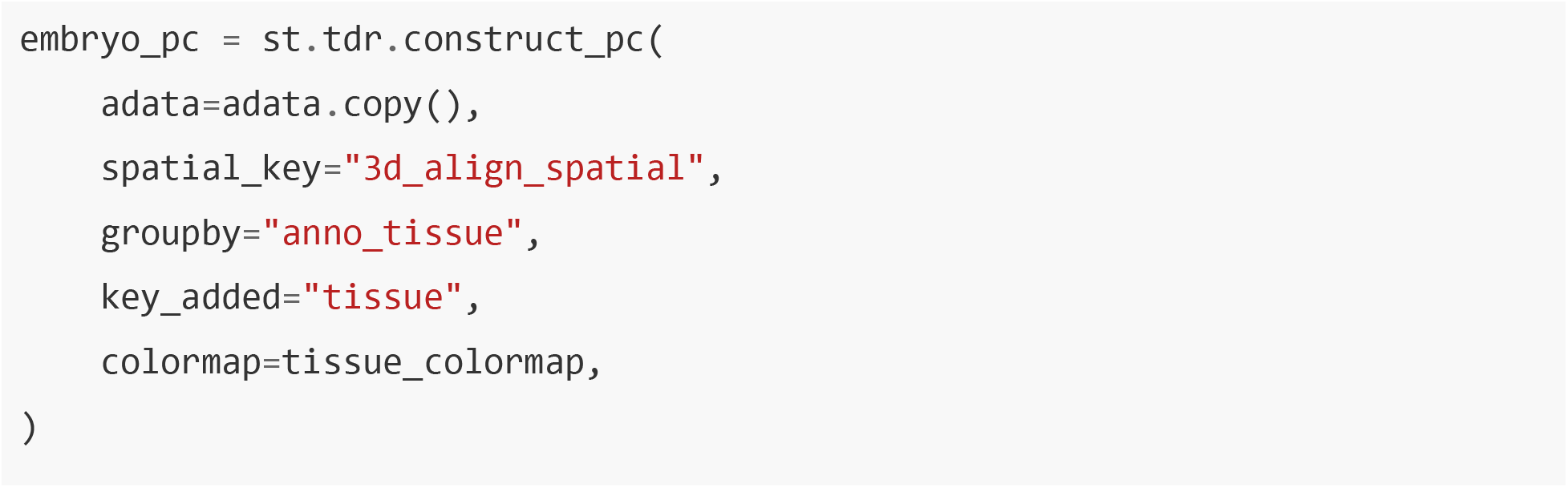

With the point cloud data, we can build upon several established algorithms to create surface polygon meshes for the entire embryo or different organs, as shown below:

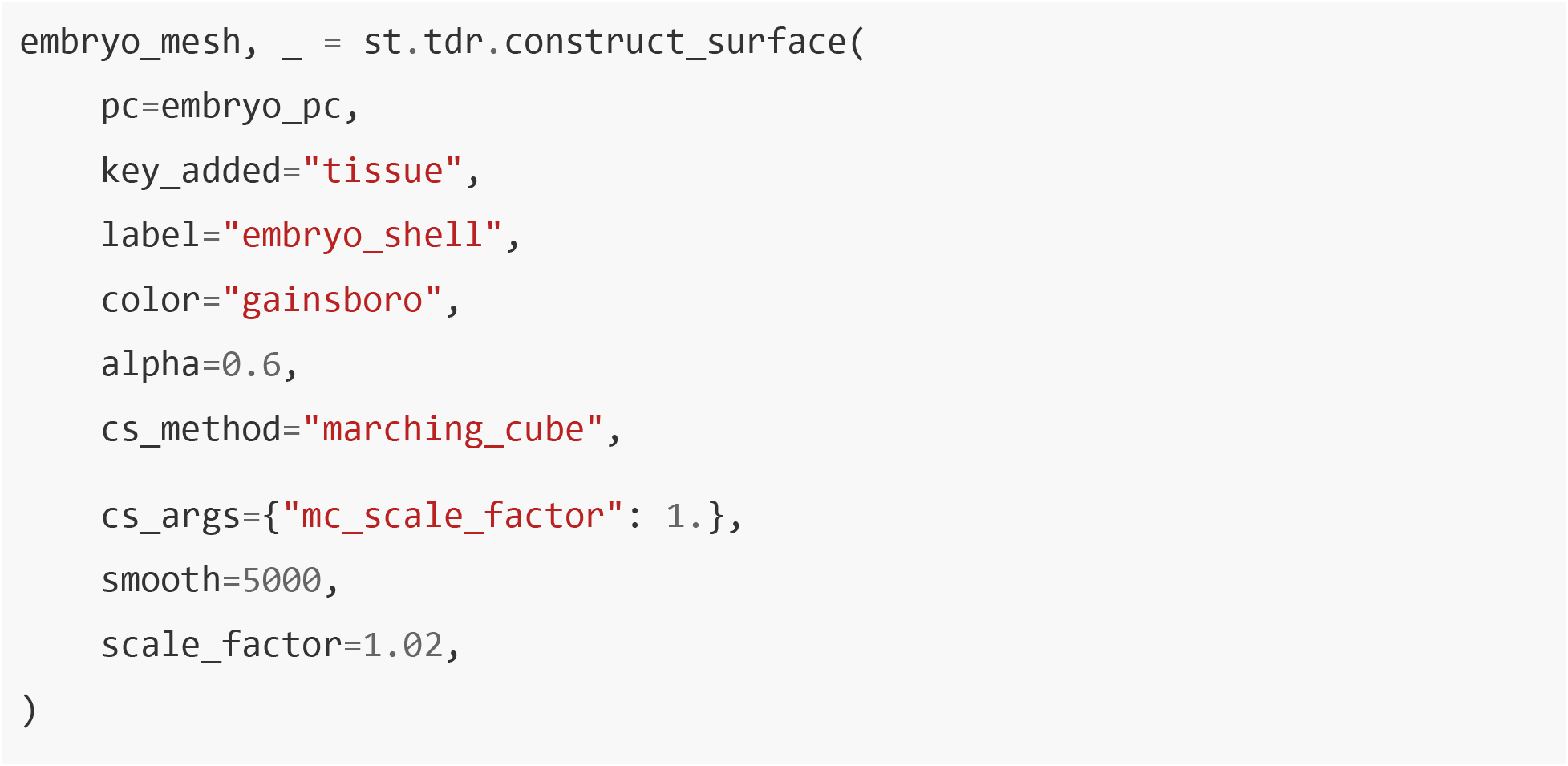

The methods for constructing surface meshes include:

- ‘pyvista’: Generate a 3D tetrahedral mesh based on pyvista.
- ‘alpha_shape’: Computes a triangle mesh based on the alpha shape algorithm.
- ‘ball_pivoting’: Computes a triangle mesh based on the Ball Pivoting algorithm.
- ‘poisson’: Computes a triangle mesh based on the Screened Poisson Reconstruction.
- ‘marching_cube’: Computes a triangle mesh based on the marching cube algorithm.

We next set the diameter of the each cell’s 3D geometry as that from the segmentation to obtain 3D volumetric meshes of each cell. In Spateo, three geometries, including cube, sphere, and ellipsoid, are supported:

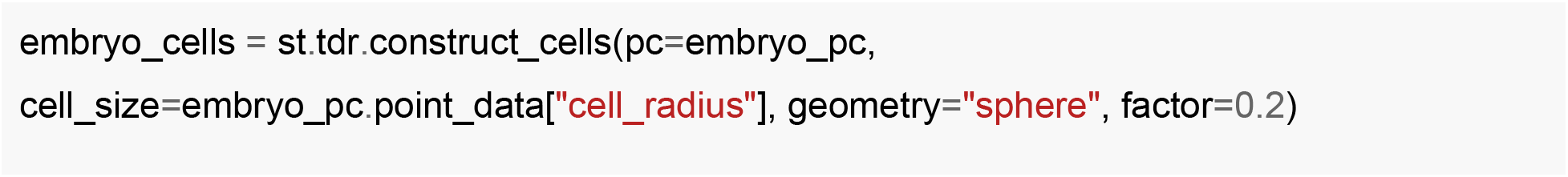

We can also voxelize the closed surface mesh into 3D voxels, where the size of voxels is determined by the density parameter:

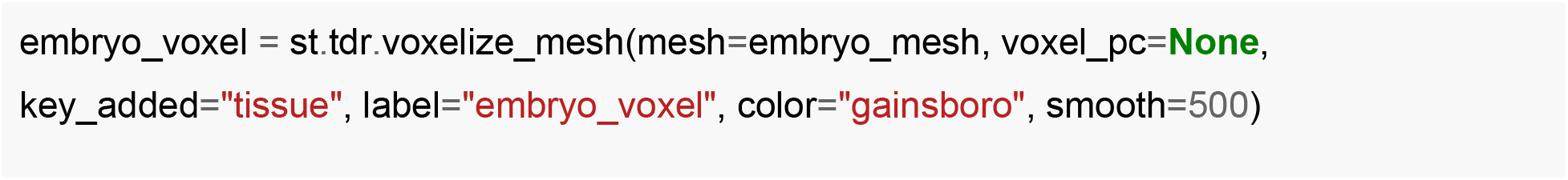

The implementation of building point cloud, surface mesh, and 3D cell geometry or voxel models in Spateo is a highly flexible strategy that can be generally applied to a single organ or the entire embryo.

##### Automatically identifying anterior-posterior (A-P) and dorsal-ventral (D-V) axes

To identify the A-P and D-V axes of the reconstructed 3D model of an embryo, we apply principal component analysis (PCA) to the 3D coordinates of aligned buckets. We then manually select the principal components to define the axes. For the Drosophila embryo used in this paper, the first and third principal components correspond to the A-P and D-V axes respectively, as shown below:

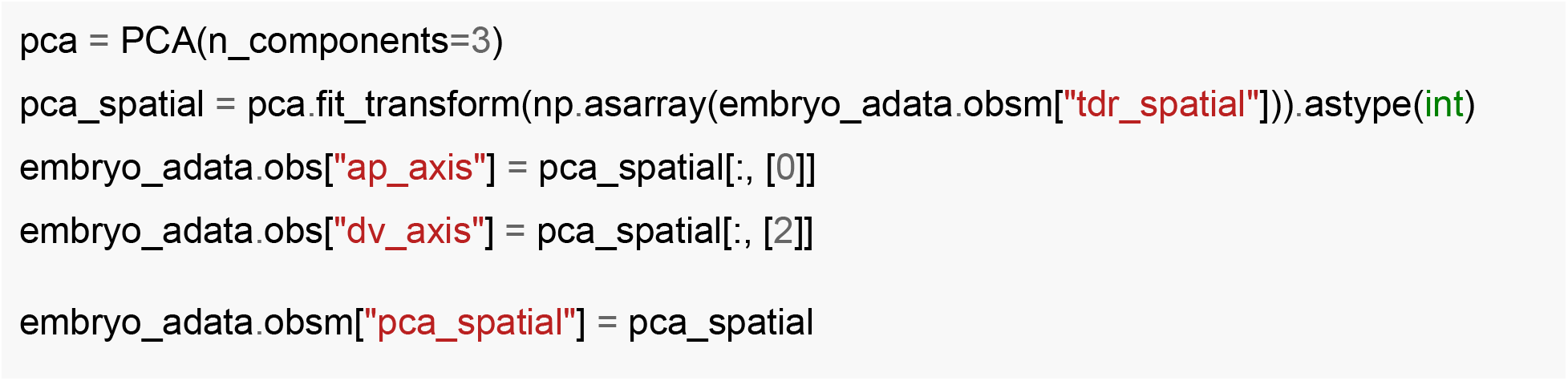

To identify axis-dependent genes, we next projected each cell to the axis to define the “pseudospace” for each cell based on the distance from the projection point to the start projection point along the axis. We next performed a GLM regression to identify genes that significantly change along the axis.

##### Volumetric and morphometric analyses

With the reconstructed 3D voxel model of the organ or embryo, we can categorize morphogenesis modes for each organ, including organ expansion, shrinkage, migration and convergence. We can further calculate a series of morphometric properties, including length, surface area, volume, and cell density:

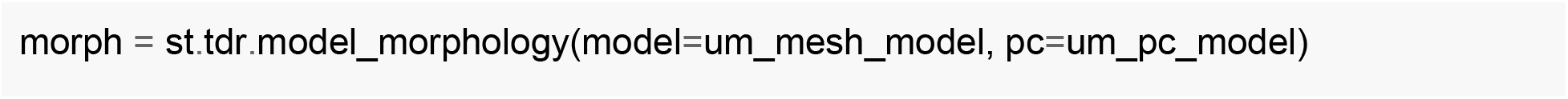

By comparing each morphometric quantity across different time points, we can reveal the temporal morphometric kinetics at organ or embryo level.

##### Principal curve and principal graph analyses of organs

A principal curve or graph is a *p*-dimensional curve or graph that passes through the middle of a data cloud. Previously, principal curves or graphs have been used to infer the pseudotemporal trajectories of linear, bifurcated, circular or other complex biological processes from single cell dataset, e.g. scRNA-seq. Here, we extend their application to reveal the structure of an organ based on the reconstructed 3D models. In Spateo, we incorporated three powerful approaches to learn the principal curve or graph that represents the organ skeleton:**NLPCA** (Nonlinear principal component analyses), two **RGE** (reversed graph embedding) algorithms, including **SimplePPT** (Simple principal tree algorithm) and **DDRTree** (Discriminative dimensionality reduction via learning a tree), and **ElPiGraph** (elastic principal graphs).

###### NLPCA

NLPCA is a global learning algorithm, implemented in prinPy (https://github.com/artusoma/prinPy) that we adapted in Spateo, to compute the principal curve via nonlinear principal component analysis. This algorithm starts with making an initial guess of a principal curve and iteratively refine the curve by creating an autoassociative neural network with a “bottle-neck” layer which forces the network to learn the most important features of the data (https://github.com/artusoma/prinPy).

###### RGE

Reversed graph embedding (RGE) is a general and powerful framework of graph learning that was championed in accurately and robustly inferring complex pseudotemporal trajectories from scRNA-seq or scATAC-seq datasets. The key novelty of the RGE is that it simultaneously learns a principal graph of the cell trajectory and often a low dimensional representation of the single cell dataset that can be mapped back to the original high-dimensional space. Various RGE algorithms have been developed, including SimplePPT, DDRTree, and others, each is developed for certain learning tasks.

The first RGE technique proposed is the SimplePPT algorithm which is tailored for learning a tree structure in the original space, or in some lower dimension retrieved by dimensionality reduction methods such as PCA. DDRTree is a novel extension of simplePPT from the RGE family, and is used by Monocle 2 (Qiu et al., 2017) as the default RGE technique. Compared to the SimplePPT algorithm, it provides two key features: First, DDRTree explicitly learns the principal graph while simultaneously reducing its dimensionality and learning the trajectory. Second, DDRTree also dramatically reduces the computational cost by clustering cells into different groups while learning the principal graph and performing dimension reduction.

###### ElPiGraph

ElPiGraph is a scalable and robust method for approximation of datasets with complex structures, via approximating complex topologies with principal graph ensembles that can be combined into a consensus principal graph which does not require computing the complete data distance matrix or the data point neighbourhood graph (Albergante et al., 2020).

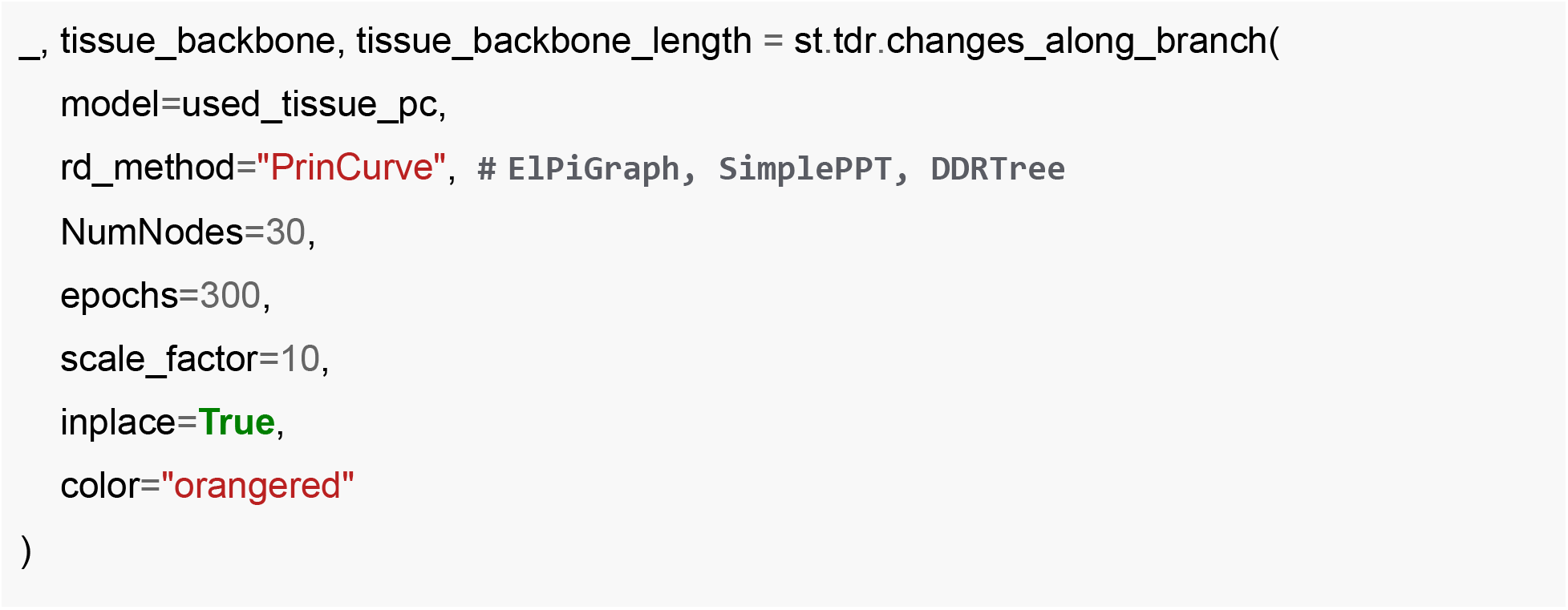

Once a principal curve or graph is constructed, similar to pseudotime algorithms that are implemented in Monocle ⅔, we can project each cell in the physical 3D space to the nearest points on the principal curve or graph. Then we can define a root principal point, such as the head point of the principal graph, and calculate the geodesic distance along the curve or graph to define a measure of ***pseudo-space*.** We use *“**pseudo-space”*** to identify principal-curve/graph dependent genes similar to pseudotime-dependent genes, as previously implemented in Monocle 2/3.

##### Build 3D continuous expression model

To reveal the continuous gene expression pattern across 3D space, we developed a deep learning framework to map the spatial coordinates to the gene expression with three consecutive fully-connected networks which use common activation functions such as ReLU or Leaky ReLU. Then we can use the trained neural network model to perform arbitrary slicing to reveal gene expression gradients within the 3D space.

##### Morphometric vector field and morphometric differential geometry analyses

The concept of vector field, a *vector-valued function* that assigns a vector for any point in a space, was originally introduced in physics and used to model the speed and direction of moving fluid or the strength and direction of magnetic or gravitational forces through the physical space. A vector field is often visualized as assigning vectors on a grid of points in a *n*-dimensional space (a grid quiver plot) or a streamline plot. The introduction of RNA velocity, defined as the time-derivative of the spliced RNA 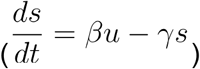, resulted in low dimensional representation of local cell fate predictions, visualized with grid quiver plots. Although such visualizations have been treated as the “vector field”, the development of *dynamo* finally enabled the reconstruction of vector fields in functional form. It is worth noting that once we have an analytical vector field, the differential geometry quantities that we can calculate will have direct physical meanings.

##### Preprocessing for morphometric vector field reconstruction

A series of preprocessing steps are required for obtaining pairs of current 3D spatial coordinates and migration velocity vectors, required for morphometric vector field reconstruction of organ morphogenesis. Specifically, in this study, after we reconstructed 3D models of whole Drosophila embryos, we set the embryo from the first time point as the reference by transforming the coordinates of the embryo such that the centroid of the embryo is located at origin, where the A-P axis and D-V axis correspond to the *x* and *y* dimension in the coordinate system respectively. To place embryo models from later time points to the same coordinate system of the initial reference embryo model, we align 3D embryos across time points. To overcome the computational burden of aligning whole embryos, we downsample 2,000 cells from each embryo and use these cells to perform 3D alignment with PASTE. Next, we rotate and translate the embryos from later time points based on GPA (generalized weighted Procrustes analysis) to place embryos from later time points to the same coordinate system. When there are multiple time points, we perform sequential 3D embryo alignment, similar to that for 2D serial slices. After aligning and transforming the coordinates of embryos, we are ready to calculate pairs of current 3D spatial coordinates and migration velocity vectors from the aligned embryo across time. We focus on analyzing individual organs instead of the entire embryo, which is more practical given the complexity of the whole-embryo migration pattern and the imperfect data quality. In particular, we focus on midgut and CNS from E7-9h and E9-10h. We first align all the cells of midgut or CNS between two time points, again using PASTE, but this time reduce the weight for the spatial preservation by setting a small *αα* = 1*e* — 3 to allow significant cell migration and give more weights on identifying highly similar cells across time points. We next iterate through the optimal transport matrix Π and assign each cell from E7-9h to another mostly like cell in E9-10. Because the cells from two time points are aligned in the coordinate system, we can take the coordinates *X_t_* from the early time point and the difference between the future time point to the current time point as velocity vectors or *V_t_* = *X*_*t*+1_–*X_t_*. The pairs of *X_t_*, *V_t_* for all cells at a prior time point can then be used to learn the vector field.

The *st.tdr.cell_directions* function from Spateo enables us to learn the mapping from cells from the early time point to the later time point, as shown below:

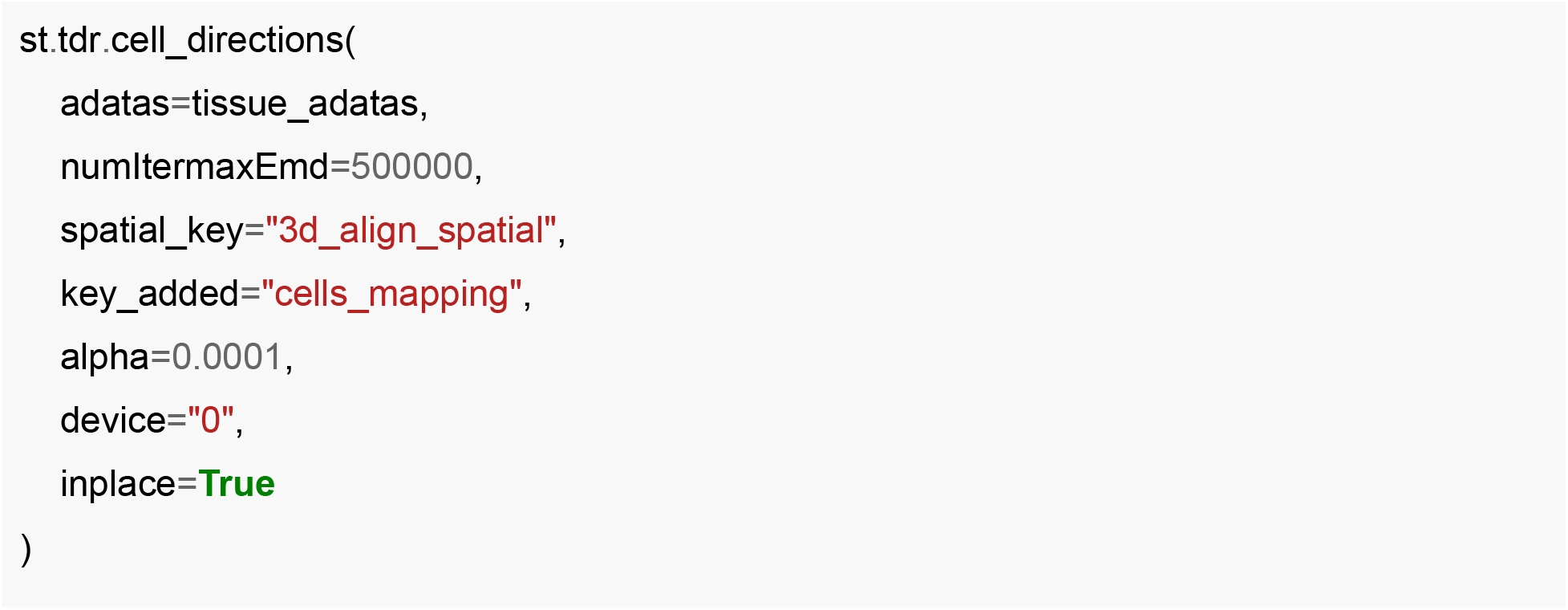

##### Reconstruct morphometric vector field with the sparseVFC algorithm

In order to learn morphometric vector fields, we consider a set of pairs of 3D physical coordinates of cell 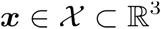 and migration velocities computed from cell alignments based on the optimal transports 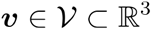, i.e. 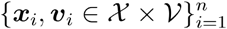, where *n* is the number of cells from a prior time point. Although the migration velocity vector based on the optimal transport alignment is noisy and discrete, we suppose that the morphogenesis of an organ follows a smooth, differentiable vector field in 3D space that assigns each cell’s current physical position ***x*** with a migration velocity vector ***ν***, as previously conceptualized (Levin, 2012). In this study, we apply sparseVFC(Ma et al., 2013b) to reconstruct a morphometric vector field of organ morphogenesis to robustly learn a vector-valued function ***f***, which outputs an migration velocity vector ***ν*** given any physical coordinate ***x*** of the cell, based on the observed noisy and discrete pairs of data 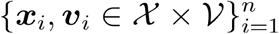.

The final loss function of vector field learning with sparseVFC is as following:

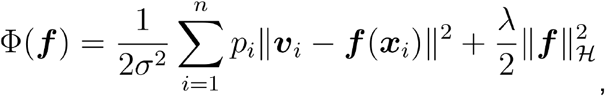

where *σ* is a parameter accounts for inlier noise, *p_i_* is a weight deciding the importance of the *i*-th data point in the loss function, λ is the regularization coefficient, 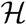 indicates the sparse reproducing kernel Hilbert space, and the second term corresponds to a vector-valued *L*_2_ regularization term. The vector field function can be evaluated at any point in 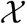, as a summation of Gaussian kernels centered on the so-called “control points”:

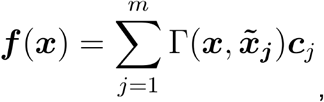

where *m* is the number of control points and 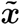 is the coordinate of the control point. ***c***’s are coefficient vectors in ℝ^3^. And the Gaussian kernel is defined as(Ma et al., 2013b):

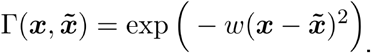

Overall, the sparseVFC algorithm (Ma et al., 2013b) consists of an **E-step** and an **M-step** to allow modeling of noise velocity (inliers) from the data. See more details at (Ma et al., 2013b;Qiu et al., 2022).

To learn the morphometric vector field in Spateo, we will use the *st.tdr.morphofiled* function as shown below:

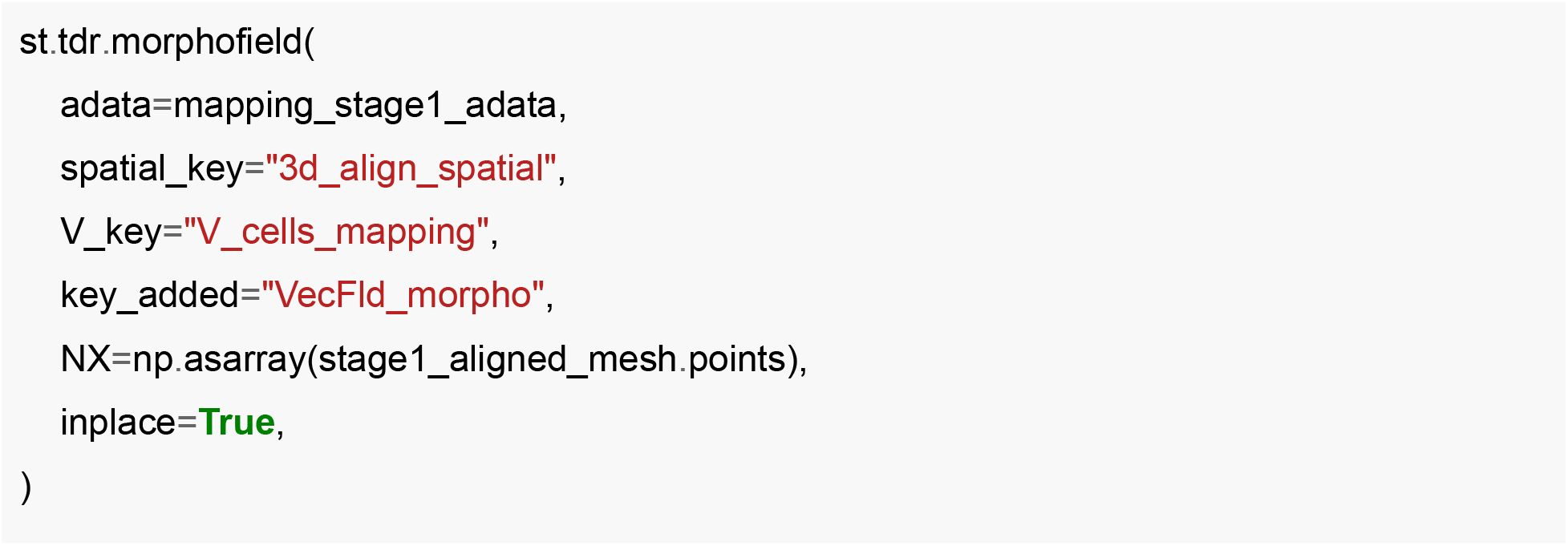

##### Differential geometry analyses of morphometric vector field: Jacobian, divergence, acceleration, curvature, curl, torsion

Once the analytical morphometric vector field is reconstructed, we can move beyond velocity to calculate higher-order differentials, including morphometric Jacobian, divergence, acceleration, curvature, curl, torsion, etc. We start with introducing *Jacobian*, a 3 × 3 matrix:

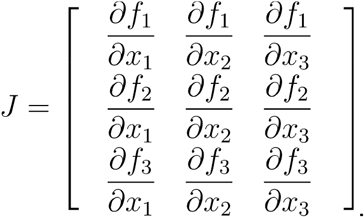

A Jacobian element *∂f_i_*/*∂x_j_* will tell you whether the migration velocity of one dimension *i* will be affected by another dimension *j*. The sum of the diagonal of the Jacobian is *divergence*:

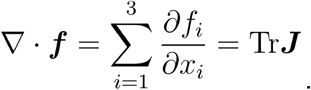

Divergence can be used to reveal whether the tissue is expanding (positive divergence) or shrinking (negative divergence). *Curl* is a quantity measuring the degree of rotation at a given point in the morphometric vector field and is defined as:

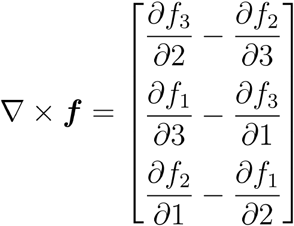

Although the input to the morphometric vector field only includes velocity vectors, we can leverage 3D manifold to calculate higher-order differentials, such as the acceleration and curvature, once a vector field is learned. The *acceleration* is the time derivative of the velocity and is defined as:

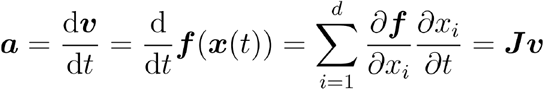

Similarly, the curvature vector of a curve is defined as the derivative of the unit tangent vector 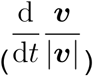, divided by the length of the tangent (|***v***|):

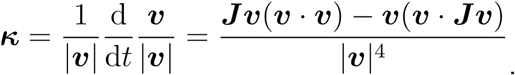

Another very interesting differential geometry quantities for 3D vector fields, that has discussed in (Qiu et al., 2022), is torsion, defined as:

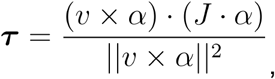

which can be used to quantify the degree of twisting of a 3D object.

In Spateo, various differential geometry quantities can be calculated as the following:

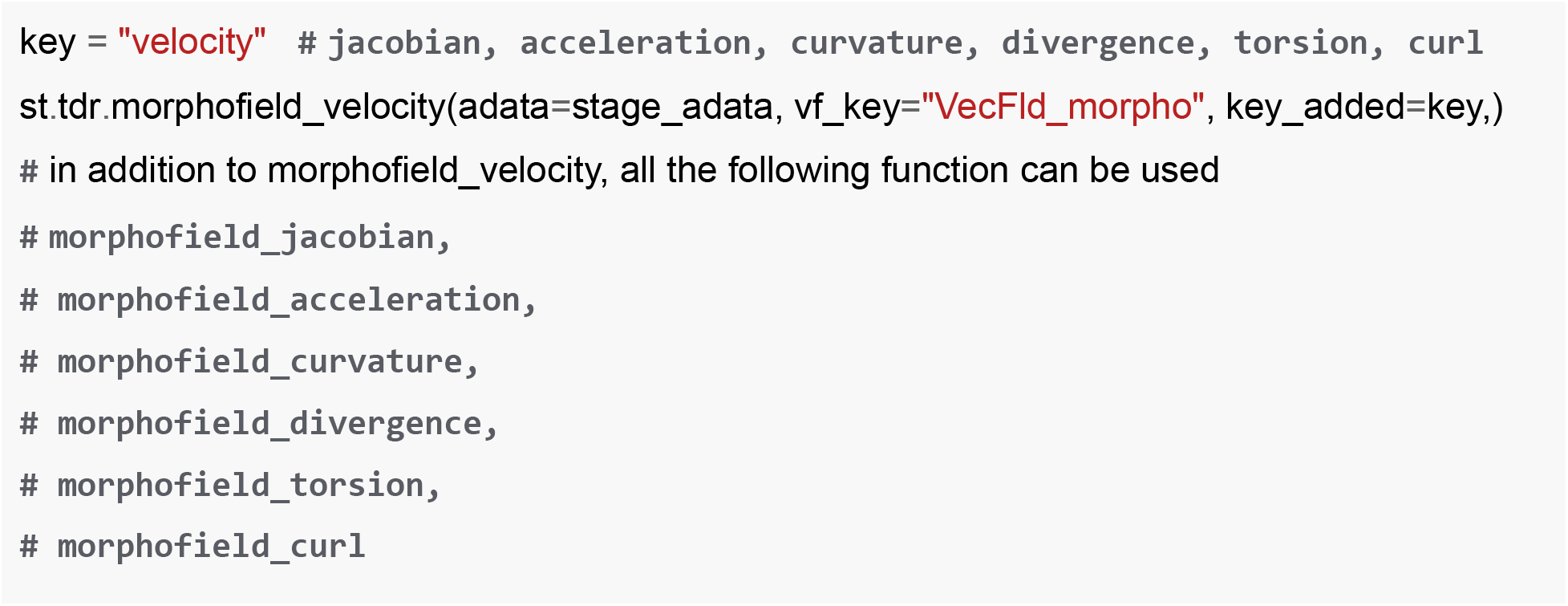

## Supplementary Methods

### Analysis of Slide-seq, seqFISH, MERFISH, STARmap and Seq-Scope data

#### Slide-seq data from the mouse hippocampus

The Slide-seq V2 dataset was downloaded from the Broad Institute single-cell portal (https://singlecell.broadinstitute.org/single_cell/study/SCP815), in the form of a raw expression matrix and array of barcode locations. To classify cell types and cellular subtypes, we applied the RCTD(Cable et al., 2022a) procedure in doublet mode using a single-cell RNA sequencing reference from the adult mouse brain(Saunders et al., 2018), selecting the highest probability cell type as the label for each spot. For the spatial colocalization and gene expression similarity analysis (**Fig. S13b**), we computed the weighted distance matrix in Euclidean space from the 10 nearest neighbors for each spot, and the weighted distance matrix in gene expression space from the 30 nearest neighbors for each spot in the space formed by the top 30 principal components (see **Construct cell type colocalization matrix**). For the spatially-constrained clustering (**Fig. S13c**), we used a resolution of 1.0 and constructed the expression graph from the 30 nearest neighbors in the space formed by the top 30 principal components, and the spatial graph from the eight nearest neighbors in 2D space. For the digitization (**Fig. S13d**), we specified a bin size of 2, kernel size of 5 and minimum area of 100 after scaling all spatial coordinates by a factor of 10. For the ligand-receptor interaction analysis (**Fig. S13e**), all pairwise permutations of ependymal cells, interneurons and choroid plexus epithelial cells were considered. To select potential ligands and receptors of interest for each sending and receiving cell type (respectively), we subsetted to the top 20 highest expressed ligands or receptors, and further subsetted to the five ligand:receptor pairs simultaneously expressed in the highest proportion of all cells of the sending and receiving cell type that are spatially proximal (defined for each spot as being one of the ten nearest neighbors). We repeated for all possible sending and receiving cell types to obtain the final list of unique ligand:receptor interactions of interest, and for each interaction computed the mean ligand:receptor product and *p*-value using 10000 permutations to obtain all necessary information for the plot.

#### seqFISH data from the mouse embryo

The processed seqFISH dataset (following segmentation and cell type identification from clustering in the PCA space) was downloaded from a custom webpage home (https://crukci.shinyapps.io/SpatialMouseAtlas/) and converted to an AnnData object with Seurat(Butler et al., 2018) v4.1.1. For the regression model analysis, we used a list of spatially differentially-expressed genes (DEGs) identified by Lohoff and Ghazanfar *et al.(Lohoff et al.,2022*)and recorded in supplementary table 8 of the original publication.

#### MERFISH data from the mouse hypothalamus

The processed MERFISH dataset was downloaded from Dryad (https://datadryad.org/stash/dataset/doi:10.5061/dryad.8t8s248). We selected one slice from the hypothalamic preoptic region, annotated with “Bregma” and labeled as being the −9^th^ section.

#### STARmap data from the mouse cortex

Details for the raw STARmap dataset can be found in the seminal study (Wang et al., 2018). For the digitization (**Fig. S12f**), we specified a bin size of 1, kernel size of 4 and minimum area of 60 after scaling all spatial coordinates by a factor of 100.

#### Seq-Scope data from the mouse liver

The raw Seq-Scope data was downloaded from the example posted on the Seq-Scope companion repository, STtools (Xi et al., 2022). Using STtools, we converted data from the base format to a Seurat object, and to an AnnData object using Seurat(Butler et al., 2018) v4.1.1. For determining the principal axes (**Fig. S13g**), we fit a two-component principal components analysis (PCA) to the spatial coordinates, with the first corresponding to the horizontal axis and the second corresponding to the vertical axis. To find genes varying along the vertical principal axis, we fit a general linear model with two degrees of freedom, a *q*-value threshold of 0.01 and a negative log-likelihood threshold of-2500. To identify GO term enrichment for identified genes, we used Enrichr(Chen et al., 2013), querying the mouse database and the “GO Biological Process 2021” gene set. For the plot (**Fig. S13h**), we set an adjusted *p*-value cutoff of 5×10^-5^.

